# Beyond Shielding: The Roles of Glycans in SARS-CoV-2 Spike Protein

**DOI:** 10.1101/2020.06.11.146522

**Authors:** Lorenzo Casalino, Zied Gaieb, Jory A. Goldsmith, Christy K. Hjorth, Abigail C. Dommer, Aoife M. Harbison, Carl A. Fogarty, Emilia P. Barros, Bryn C. Taylor, Jason S. McLellan, Elisa Fadda, Rommie E. Amaro

## Abstract

The ongoing COVID-19 pandemic caused by severe acute respiratory syndrome coronavirus 2 (SARS-CoV-2) has resulted in more than 15,000,000 infections and 600,000 deaths worldwide to date. Antibody development efforts mainly revolve around the extensively glycosylated SARS-CoV-2 spike (S) protein, which mediates the host cell entry by binding to the angiotensin-converting enzyme 2 (ACE2). Similar to many other viruses, the SARS-CoV-2 spike utilizes a glycan shield to thwart the host immune response. Here, we built a full-length model of glycosylated SARS-CoV-2 S protein, both in the open and closed states, augmenting the available structural and biological data. Multiple microsecond-long, all-atom molecular dynamics simulations were used to provide an atomistic perspective on the roles of glycans, and the protein structure and dynamics. We reveal an essential structural role of N-glycans at sites N165 and N234 in modulating the conformational dynamics of the spike’s receptor binding domain (RBD), which is responsible for ACE2 recognition. This finding is corroborated by biolayer interferometry experiments, which show that deletion of these glycans through N165A and N234A mutations significantly reduces binding to ACE2 as a result of the RBD conformational shift towards the “down” state. Additionally, end-to-end accessibility analyses outline a complete overview of the vulnerabilities of the glycan shield of SARS-CoV-2 S protein, which may be exploited by therapeutic efforts targeting this molecular machine. Overall, this work presents hitherto unseen functional and structural insights into the SARS-CoV-2 S protein and its glycan coat, providing a strategy to control the conformational plasticity of the RBD that could be harnessed for vaccine development.

## Introduction

COVID-19 is an infectious respiratory disease that started in Wuhan, China, near the end of 2019, and has now spread worldwide as a global pandemic.^1^ This is not the first time that a coronavirus (CoV) has posed a threat to human health. SARS-CoV-2 (the virus that causes COVID-19) is in the same family of viruses, *Coronaviridae*, as severe acute respiratory syndrome (SARS) and Middle East respiratory syndrome (MERS) related coronaviruses, which have resulted in previous epidemics.^2–4^ Owing to the lack of immunity, COVID-19 has already caused a catastrophic loss of human life worldwide^5^ as well as significant economic damage.^6^

Coronaviruses, including SARS-CoV-2, are lipid-enveloped positive-sense RNA viruses. Together with the host-derived membrane, a set of structural proteins provide an organizational scaffold that wraps and contains the positive-sense viral RNA. Among them, the most critical is the spike, or S, protein, which is conserved to varying degrees across the *Coronaviridae* family and plays a key role in the virus’ initial attachment and fusion with the host cell. The S protein is a class I fusion protein, synthesized as a single 1273 amino acid polypeptide chain, which associates as a trimer. Each monomer is made of two subunits, S1 and S2, and can be divided into three main topological domains, namely the head, stalk, and cytoplasmic tail (CT) (**Figure 1A**). One particularly interesting feature of the SARS-CoV-2 S protein is its adoption of a novel furin cleavage site between S1 and S2 (S1/S2), likely cleaved by the TMPRSS2 protease,^7^ and believed to prime the spike for infection.^8,9^ A second proteolytic cleavage at site S2’ releases the fusion peptide (FP), which penetrates the host cell membrane, gearing it up for fusion.^10^ A number of recently published structural studies has provided an atomic or a near-atomic understanding of the head portion of SARS-CoV-2 spike, which comprises multiple domains (**Figures 1A** and **1B**).^11,12^ The S1 subunit contains an N-terminal domain (NTD) and the receptor binding domain (RBD), where the receptor binding motif (RBM) is responsible for the interaction with the angiotensin-converting enzyme 2 (ACE2) receptor to gain entry into the host.^13^ The S2 subunit has been aptly described as “a metastable spring-loaded fusion machine” because it plays a key role in integrating the viral and host cell membranes.^14^ It contains the FP, the central helix (CH), and the connecting domain (CD). Additional domains within the S2 subunit that are not resolved in the pre-fusion state via cryo-EM or X-ray experiments include the heptad repeat 2 (HR2) and the transmembrane (TM) domains forming the stalk, and the CT (**Figures 1A** and **1B**).

**Figure 1.**
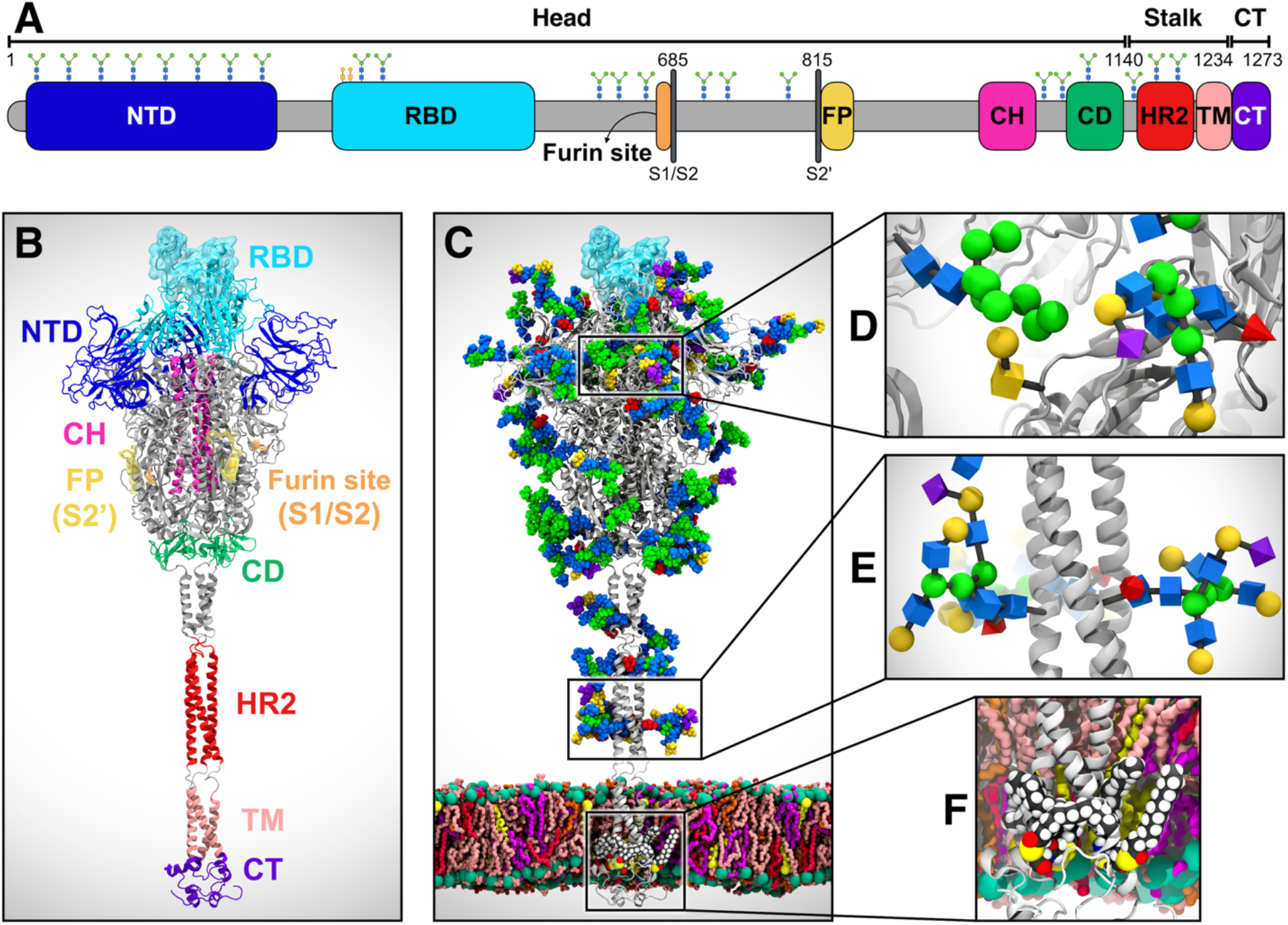
System overview. (**A**) Sequence of the full-length spike (S) protein contains the N-terminal domain (NTD, 16–291), receptor binding domain (RBD, 330–530), furin cleavage site (S1/S2), fusion peptide (FP, 817–834), central helix (CH, 987–1034), connecting domain (CD, 1080–1135), heptad repeat 2 (HR2, 1163–1210) domain, transmembrane domain (TM, 1214–1234), and cytoplasmic tail (CT, 1235–1273). Representative icons for N-glycans (blue and green) and O-glycan (yellow) are also depicted according to their position in the sequence. (**B**) Assembly of the head, stalk, and CT domains into the full-length model of the S protein in the open state. (**C**) Glycosylated and palmitoylated full-length model of the S protein in the open state embedded in a lipid bilayer mimicking the composition of the endoplasmic reticulum-Golgi intermediate compartment. Color code used for lipid tails: POPC (pink), POPE (purple), POPI (orange), POPS (red), cholesterol (yellow). P atoms of the lipid heads are shown with green spheres. Cholesterol’s O3 atoms are shown with yellow spheres. (**D**) Magnified view of S protein head glycosylation, where glycans are depicted using the Symbol Nomenclature for Glycans (SNFG). (**E**) Magnified view of S protein stalk glycosylation. (**F**) Magnified view of S protein S-palmitoylation within CT.

Another key structural feature of the S protein that eludes detailed experimental structural characterization is its extensive glycosylation, shown in **Figure 1C**. Protein glycosylation plays a crucial role in viral pathogenesis,^15–17^ as demonstrated by the characteristically thick N-glycan coating of the viral fusion proteins.^18–21^ In the HIV-1 envelope spike (Env), for example, the protein-accessible surface area is almost entirely covered in N-glycans.^20,22^ These are so densely packed that they account for more than half of the protein’s molecular weight.^23^ The biological roles of the N-glycans expressed on the surface of viral envelope glycoproteins are very diverse^16^ and are all inextricably linked to their nature. Viral entry through membrane fusion is initiated by envelope glycoproteins through molecular recognition events involving cell surface receptors, which are often mediated by specific N-glycan epitopes.^16,24–26^ Moreover, a highly dense coating of non-immunogenic or weakly immunogenic complex carbohydrates on otherwise dangerously exposed viral proteins constitutes a perfect camouflage (or shield) to evade the immune system.^16,18,19,27,28^ To this end, the HIV-1 Env glycan shield, which is largely structured by oligomannose (Man5-9) N-glycans,^18,20,27,29^ has been shown to be quite effective in allowing the virus to thwart the immune system;^15,16^ it has also been found to be responsible for the virus’ interactions with DC-SIGN C-type lectins.^30^ Contrary to HIV-1 Env, the betacoronaviruses SARS and MERS S proteins are not shielded as effectively.^15^ Furthermore, both SARS-CoV and SARS-CoV-2 spikes present a rather different glycosylation pattern from that of HIV-1 Env, with a large presence of complex N-glycans relative to oligomannose type.^11,15,31^ More specifically, the SARS-CoV-2 spike has 22 predicted N-glycosylation sites per protomer,^11,12^ of which at least 17 have been found to be occupied,^11,31^ plus at least two predicted O-glycosylation sites (see **Figures 1C-1E**).^31^

In this work, we present multiple microsecond-long, all-atom, explicitly solvated molecular dynamics (MD) simulations of the full-length SARS-CoV-2 S glycoprotein embedded in the viral membrane, with a complete glycosylation profile consistent with glycomic data.^11,31^ The simulations discussed here augment and extend the available structural and biological data to provide an atomically detailed perspective on the full-length SARS-CoV-2 S protein’s glycan coat, structure, and dynamics. Beyond shielding, our work reveals an essential structural role of N-glycans linked to N165 and N234 in modulating the conformational transitions of the RBD. Deletion of these glycans in our simulations elicits a destabilizing effect on the RBD “up” conformation. These results are corroborated via biolayer interferometry experiments showing a reduction of ACE2-binding, which underlies an increase of the RBD “down” population. Moreover, our simulations highlight how glycans camouflage SARS-CoV-2 S protein for thwarting the host immune response. A detailed analysis of the S protein’s glycan shield discloses a lack of vulnerabilities in the stalk, particularly for large molecules, whereas the head region appears to be a more viable target. Importantly, an in-depth overview of the RBM accessibility indicates a remarkably different extent of glycan shield between the “up” and “down” RBD conformations. Overall, this work provides an atomic-level perspective on the SARS-CoV-2 S protein, highlighting the importance of glycans not only as shielding devices for immune evasion, but also as essential structural elements for virus infectivity. These insights lay the foundations for a possible strategy to modulate the RBD conformational plasticity and virus infectivity, which could be harnessed in the development of therapeutics aimed at fighting the pandemic threat.

## Results & Discussion

### All-Atom MD Simulations of the Full-Length Model of the SARS-CoV-2 S Protein in Open and Closed States

In this work, we built a complete, full-length model of the glycosylated SARS-CoV-2 S protein in both closed and open states, hereafter referred to as “Closed” and “Open,” respectively. Closed is based on cryo-EM structure 6VXX, where all three RBDs are in a “down” conformation.^12^ Open is based on cryo-EM structure 6VSB, where the RBD within chain A (RBD-A) is in an “up” conformation.^32^ A detailed view of chains A, B, and C and the RBD “up/down” conformations as in Open/Closed is shown in **Figure S1** of the Supporting Information (SI). These models were built in three steps, as fully described in the Materials and Methods section in the SI: (i) the “head,” comprising S1/S2 subunits until residue 1140 and based on the above-mentioned experimental coordinates;^12,32^ (ii) the “stalk” (residues 1141–1234), comprising HR2 and TM domains, and constructed through homology modeling using Modeller;^33^ (iii) the CT (residues 1235–1273) built using I-TASSER (**Figures 1A)**.^34^ The modeled constructs (see **Figure 1B** for Open) were fully glycosylated at N-/O-glycosylation sites^11,31^ (**Figures 1C-E**), and further refined with cysteine palmitoylation within the CT (**Figure 1F**).^35,36^ An asymmetric (i.e., not specular across monomers) site-specific glycoprofile totaling 70 glycans (22 × 3_monomers_ N-glycans and 2_chainA_ + 1_chainB_ + 1_chainC_ O-glycans) was derived according to available glycoanalytic data (**Tables S1-S3** for composition).^11,31^ We remark that the saccharides originally solved in the cryo-EM structures have been generally retained and utilized as a basic scaffold, when possible, to build the full glycan structure using CHARMM-GUI (details in the Materials and Methods section in the SI).^37–39^ The full-length structures were embedded into an equilibrated all-atom membrane bilayer with a composition mimicking the endoplasmic reticulum-Golgi intermediate compartment (ERGIC, **Table S4** for composition), where the virus buds. The membrane was generated through CHARMM-GUI.^40^ Subsequently, explicit water molecules and ions were added, affording two final systems each tallying ∼1.7 million atoms (**Figure 1B**). Using CHARMM36 all-atom additive force fields^41,42^ and NAMD 2.14,^43^ multiple replicas of all-atom MD simulations of the Open (6x) and Closed (3x) systems were run on the NSF Frontera computing system at the Texas Advanced Computing Center (TACC), achieving benchmarks of ∼60 ns/day on 256 nodes for cumulative extensive sampling of ∼4.2 and ∼1.7 µs, respectively (**Table S5, Movie S1**). Note that, since RBD-A within Open is captured in a metastable state,^32^ Open was simulated for a longer time than Closed. Visual inspection of trajectories was performed with VMD^44^ that, together with MDtraj,^45^ was also used for analyses.

Topological domain-specific root-mean-square-deviation (RMSD) relative to the starting structures was calculated to examine the structural stability along the simulations (**Figures S2-S3**). Overall, the two systems showed structural convergence of the extracellular topological domains (head and stalk) within ∼400 ns. No significant difference was observed in the RMSD values of the stalk between the systems (**Figure S3**). Besides displaying a replica-specific convergence of RMSD values, the triple-stranded coiled-coil structural motif of the stalk persists throughout the simulations (**Movie S1**). Notably, as a result of secondary structure prediction (see Material and Methods in the SI), the stalk model includes two loops that break the alpha-helix within each strand (**Figure 1B**). Along with linked N-glycans, these loops confer critical dynamic properties to the stalk, allowing it to bend to accommodate the head’s wiggling motions while remaining anchored to the membrane (**Movie S1**). The RMSD values of the CT domain show extremely large deviations and a mixed range of dynamic behaviors across replicas (**Figure S3**). While the first section of CT remains anchored to the inner-leaflet of the lipid bilayer through the palmitoylated cysteines, the rest is highly flexible and solvent exposed.

The root-mean-square-fluctuation (RMSF) values of the glycans were measured to determine their average flexibility around their mean position across the trajectories of all replicas (**Figure S4**). These values indicate a diverse range of mobility for specific sites, which is mostly dependent on solvent exposure, branching, and sequence. Complex-type N-glycans located on the NTD exhibit larger RMSF values than oligomannose and other N-glycans populating the head, including the ones in the RBD at sites N331 and N343. Interestingly, the O-glycans linked in the immediate vicinity of the RBD, namely at T323 and S325, show the lowest RMSF values. This is mostly attributable to their small size; however, their position may be critical for modulating RBD conformational changes, although we note that both positions are not consistently occupied. Finally, the large tetrantennary N-glycans in the stalk region at N1173 and N1194, modeled consistently with experimental data,^11^ show the most extensive fluctuations, suggesting their good shielding potential.

The stability of the membrane bilayer was also monitored by tracking the time evolution of a series of structural properties, including area per lipid, membrane thickness, order parameters, and phospholipid tilt angles. Equilibrium area per lipid and P–P membrane thickness plots are shown in **Figure S5**. These results are in agreement with known POPC and previously reported binary POPC + X trends, where X is any other lipid added to the mix.^46–52^

### N-Glycans at N165 and N234 modulate the RBD Conformational Dynamics

SARS-CoV and SARS-CoV-2 S glycoproteins share 76% sequence identity,^12^ and 18 out of 22 N-glycosylation sites found on SARS-CoV-2 are conserved from SARS-CoV. Based on the analysis of our simulations of the full-length model of SARS-CoV-2 S protein, we identified two N-glycans linked to N165 and N234 in the NTD, which, given their strategic position and structure, we hypothesized may play a role in the RBD conformational dynamics. Our simulations show that in Closed, the N-glycan at N234 within NTD-B, modeled as Man9 in agreement with glycoanalytic data,^11^ is pointing away from the core of the S protein and is directed toward the solvent (**Figure 2A**); in contrast, in Open, where the RBD of chain A is “up,” the same Man9 linked to N234 of NTD-B is directed inward, inserting itself into the large volume left vacant by the opening of RBD-A (**Figures 2B** and **2C**),. Because the glycans were built by structural alignment on the linked GlcNAc (or chitobiose) residues, wherever available, the relative position of the existing GlcNAc or chitobiose may affect the orientation of the entire N-glycan. Indeed, the N-glycan core is quite rigid.^53^ To further assess the possible impact of the force fields and/or of the starting cryo-EM structure on the simulations described here, we performed an additional set of simulations of the open SARS-CoV-2 S protein’s head using AMBER ff14SB/GLYCAM06j-1 force fields^54,55^ and an alternative initial cryo-EM structure (PDB ID: 6VYB),^56^ which presents the GlcNAc linked to N234 in a slightly different orientation. These simulations, detailed in **Section 2** of SI, show that the Man9 glycan at N234 progressively inserts itself to reach the apical core of the trimer through interactions with the surface of the closed RBD domain located across from it (**Movie S2**), adopting a structural arrangement similar to the one described in **Figure 2B**. The highly conserved N-glycan at N165 also inserts itself between NTD-B and RBD-A, either making extensive interactions with RBD-A or occupying the volume left vacant following its opening (**Figure 2C**). This was also observed in the additional simulations described in **Section 2** of the SI.

**Figure 2.**
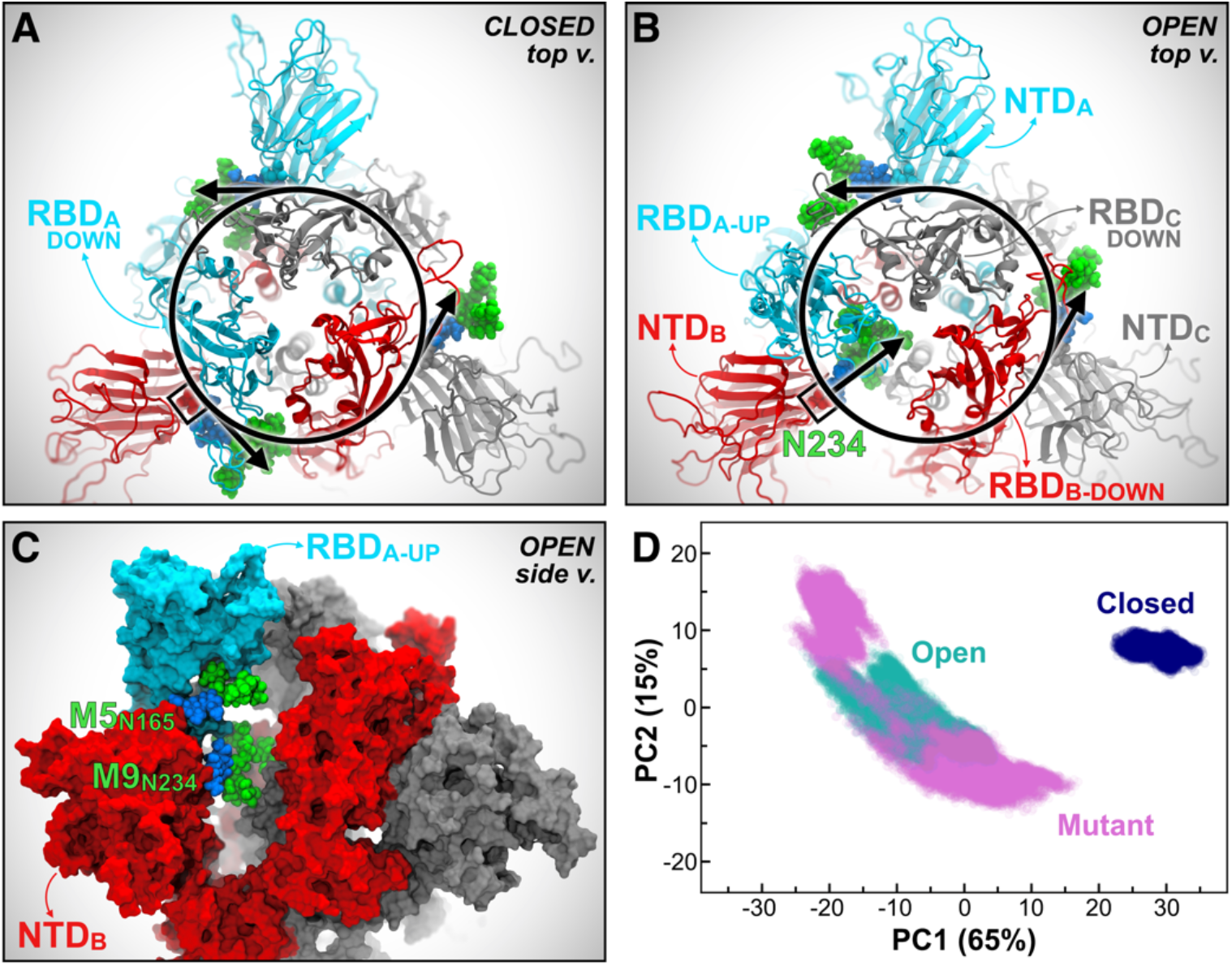
N234A and N165A mutations show increased instability of RBD-A in the “up” state. (**A-B**) Top view of the S protein as in the Closed (**A**) and Open (**B**) systems. Protein is represented with cartoons, colored in cyan, red, and gray for chains A, B and C, respectively. Oligomannose N-glycans at position N234 from all chains are depicted with VdW spheres, where GlcNAc is colored in blue and Man in green. In Closed (**A**), all the N-glycans at N234 are tangential to a hypothetical circle going through N234. In Open (**B**), the N-glycan at N234 of chain B moves inward, filling in the vacancy under RBD-A in the “up” conformation. (**C**) Side view of the S protein (surface representation) in Open, where the RBD of chain A (RBD-A, cyan) is stabilized by N-glycans at N165 and N234 in the “up” conformation. Same color scheme as panels A and B is applied. (**D**) PCA plot showing PC1 vs. PC2 of RBD-A (residues 330–530) in Closed, Open, and Mutant in blue, teal, and magenta, respectively. The amount (%) of variance accounted by each PC is shown between parentheses.

In light of these observations, an additional third system, “Mutant,” was generated from Open by introducing N165A and N234A mutations, which led to glycan deletions at the respective sites. Mutant was then simulated through all-atom MD for six replicas totaling ∼4.2 µs (**Table S5**). To determine the structural consequences of removing these two glycans, we compared the conformational landscape of the RBD in Open, Mutant, and Closed systems using principal component analysis (PCA). Scatter plot projections of RBD-A dynamics onto the first two eigenvectors (PC1 vs. PC2), i.e. the two motions with the largest variance in the trajectories (65% and 15%, respectively), clearly indicate that the RBD-A in Mutant explores a larger conformational space than in Open, which in turn is considerably larger than in Closed (**Figure 2D**). Although the difference between Open and Closed is immediately explained by the RBD-A state (“up/down”), the PCA landscape pinpoints a more stable RBD “up” conformation in the presence of N165 and N234, thus suggesting a structural role for the respective N-linked glycans (**Movie S3**). Detailed analysis of the PCA space explored by RBD-A, including the comparison between Open and Mutant only and the per-replica contribution in all systems, is presented in **Figures S6** and **S7**. Analysis of RMSD along the trajectory further highlights the instability of RBD-A in Mutant (**Figure S2**). This behavior is confirmed even in the case of a single point mutation (N234A), as revealed by the additional independent simulations of the spike head described in **Section 2** of SI. In contrast to RBD-A, RBD-B and RBD-C are in the “down” conformation in all three systems, showing mostly stable overlapping PCA distributions (**Figure S7**).

To obtain further insights into the RBD-A dynamic behavior and examine the differences in the explored conformational spaces revealed by PCA, we monitored the variations along the dynamics of two angles describing RBD-A fluctuations, hereafter referred to as the “lateral-angle” and “axial-angle.” The lateral-angle reports an in-plane motion of the RBD along a hypothetical circle centered on the central helices of the spike (**Figure 3A**). This angle is described by three points corresponding to the (i) center of mass (COM) of RBD-A core β-sheets at frame 0, (ii - vertex) COM of the top section of the CH, and (iii) COM of RBD-A core β-sheets at frame n. The axial-angle identifies a tilting motion of the RBD either away from or toward the CH of the spike (**Figure 3B)**. This angle is defined by three points corresponding to (i) the COM of RBD-A core β-sheets, (ii - vertex) the COM of the CH, and (iii) the COM of the top section of the CH. The distributions of the lateral and axial-angle fluctuations along the trajectories of each system are shown in **Figures 3C** and **3D**. Angle variations were calculated with respect to their initial values at frame 0 of the trajectories. Full details on angle definitions and calculations are outlined in the Material and Methods section in the SI.

**Figure 3.**
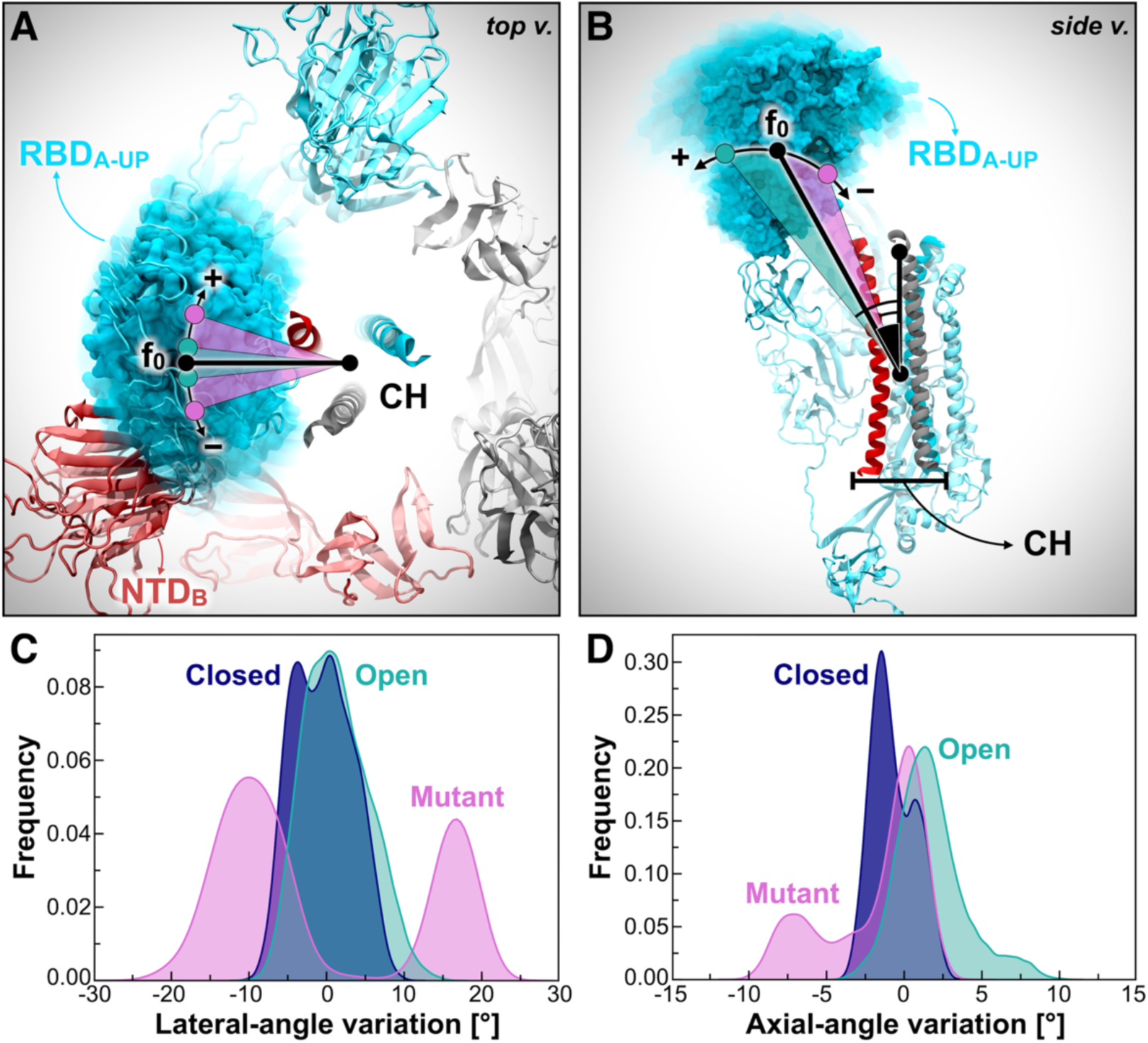
RBD-A lateral and axial-angle fluctuations. (**A, B**) RBD-A lateral-angle (**A**) and axial-angle (**B**), where chains A, B, and C of the spike are represented as ribbons colored in cyan, red, and gray, respectively. Positive and negative variations with respect to the initial frame (0) are indicated with the “+” and “−” symbols, respectively. N-/O-Glycans and some structural domains of the spike are omitted for clarity. (**C, D**) Distributions of RBD-A lateral-angle (**C**) and axial-angle (**D**) fluctuations along the trajectories (across all replicas) in Closed (blue), Open (teal), and Mutant (magenta). Angle variations were calculated with respect to their value at frame 0. Frequencies have been normalized within the respective data sets.

In agreement with the dynamic behavior revealed by PCA (**Figure 2D)**, the RBD-A lateral-angle displays a well-defined single distribution centered on zero in Open, whereas it exhibits a bimodal population in Mutant (**Figure 3C**). This analysis shows that the presence of the NTD-B N-glycans at N165 and N234 is crucial to stabilize RBD-A “up” conformation, preventing it from undergoing disordered motions. Interestingly, the axial-angle analysis results display a similar behavior of RBD-A in Open and Mutant, showing a long tail distribution in both systems (**Figure 3D**). However, Mutant exhibits a significant negative trend, whereas Open shows a positive trend. This suggests that the N-glycans at N165 and N234 may not only play a critical role in stabilizing the RBD “up” conformation, but also in modulating RBD opening and closing, thus potentially affecting binding to ACE2. Indeed, whereas the RBD in the “down” state overlies on the CH of the spike trimer, thus not being accessible to ACE2, when it transitions to the “up” state it detaches from the central axis, protruding into the solvent and becoming available for binding.

To confirm the hypothesis unveiled by our simulations, we sought to experimentally assess the effects of N165 and N234 deletions on the RBD state. To quantify RBD accessibility in solution, we used biolayer interferometry to measure ACE2 binding to S protein containing either N165A or N234A substitutions. As a result, ablation of these N-linked sites reduces the binding responses for N165A or N234A to ∼90% and ∼60% (p=0.0051 and p=0.0002, Student’s t-test) as high as the parent S-2P variant, respectively (**Figure 4**). We remark that the S-2P variant of the S protein bears two consecutive proline substitutions within the S2 subunit that are introduced to stabilize the pre-fusion conformation.^32^ Importantly, a negative control spike (HexaPro), engineered with S383C/D985C mutations to lock all three RBDs in the closed state through a disulfide bond,^57^ shows no binding to ACE2. These experiments corroborate the hypothesis that N165 and N234 glycans stabilize the RBD “up” conformation, therefore facilitating binding to ACE2, with the N234 glycan playing a larger role than N165.

**Figure 4.**
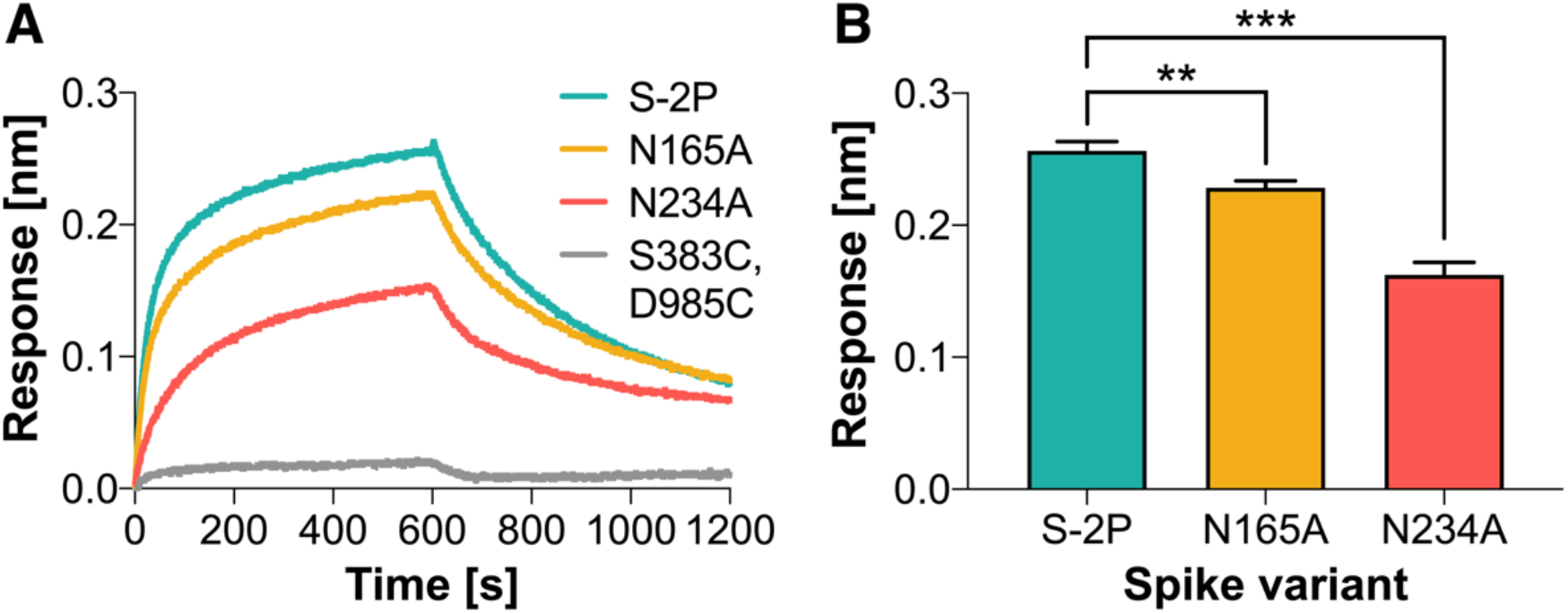
N234A and N165A mutations reduce RBD binding to ACE2. (**A**) Representative biolayer interferometry sensorgrams showing binding of ACE2 to spike variants. (**B**) Binding responses for biolayer interferometry measurements of ACE2 binding to spike variants. Data are shown as mean ± S.D. from three independent experiments for each variant. Asterisks represent statistical significance (Student’s t test; *0.01<p<0.05, **0.001>p>0.01, ***0.0001<p<0.001).

To characterize how the N-glycans at N165 and N234 stabilize the RBD in the “up” state, we examined their interaction with the S protein through hydrogen bond analysis. Man9 linked to N234 on NTD-B deeply extends into the large pocket created by the opening of the RBD-A (**Figure 5C**). Specifically, it largely interconnects with the lower part of RBD-A (H519 in particular), propping it up from underneath, makes stable hydrogen bonds with D198 of NTD-B, and interacts as deep as R983, D985, E988 located within the CH of chain B (**Movie S3**). All of these hydrogen bonds are stable for more than 40% of the 4.2 µs trajectory of the Open system, with the majority of hydrogen bonds formed with the CH of chain B and RBD-A (see **Figure 5A**). The N-glycan at N165 is more exposed to the solvent relative to the Man9 glycan at N234; nevertheless, it extensively engages in interactions with the “up” conformation of RBD-A (∼90% frequency) with a variable hydrogen bonding pattern across simulation replicas (**Figures 5B** and **5D, Figure S8, Movie S3**). The per-replica hydrogen bond networks observed for the N-glycans at N234 and N165 are shown in **Figure S9**.

**Figure 5.**
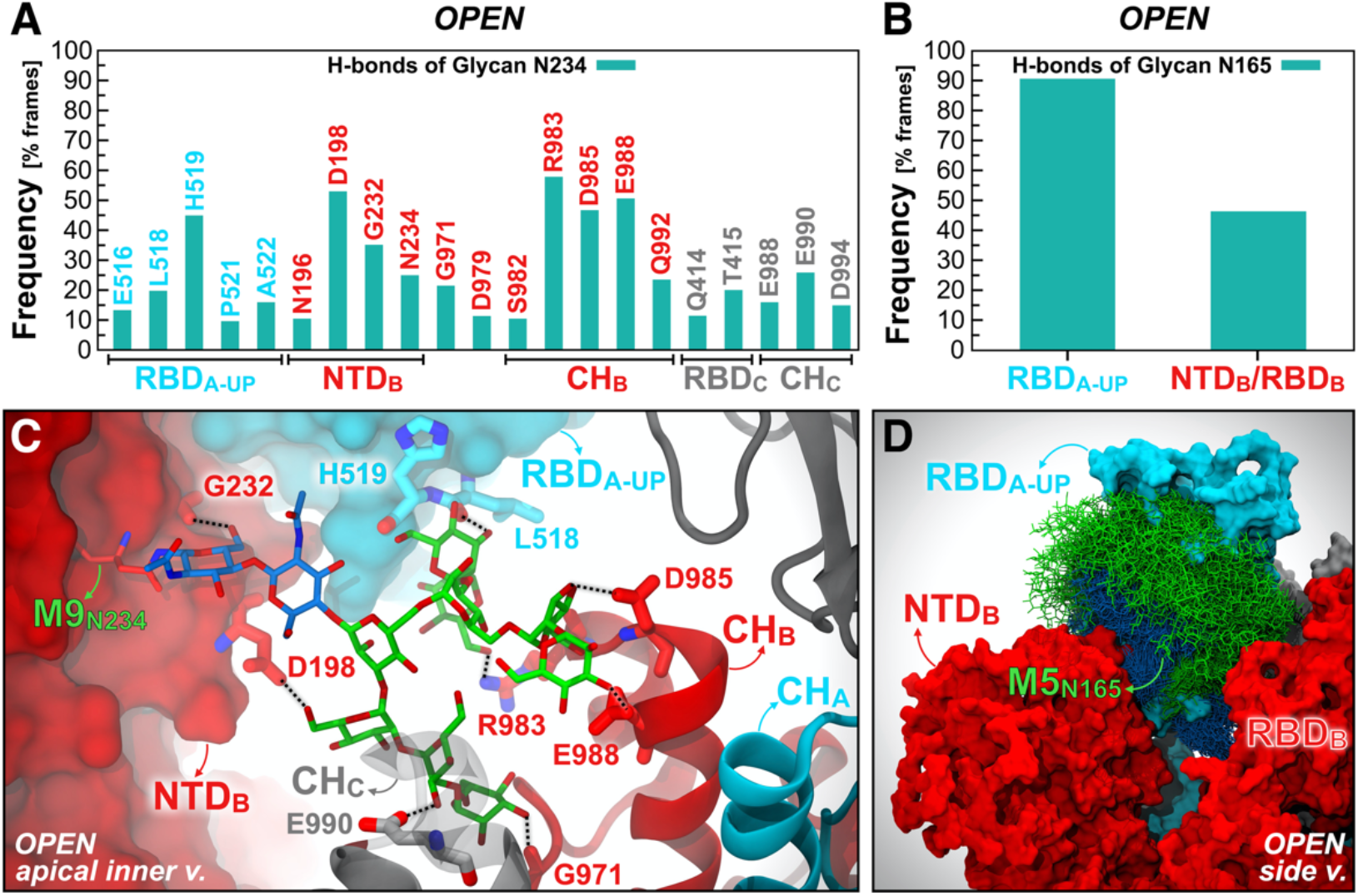
Hydrogen bond interactions of N-glycans at N234 and N165. The main hydrogen bond interactions of N-glycans at N234 (**A**) and N165 (**B**) within the Open system are shown as occupancy across all replicas (% frames). (**C**) A snapshot capturing Man9 glycan at N234 (licorice representation) within NTD-B establishing multiple hydrogen bonds with S protein residues (thicker licorice representation) belonging to RBD-A (cyan surface), NTD-B (red surface), CH-B (red cartoons), and CH-C (gray cartoons). GlcNAc and Man carbons are colored in blue and green, respectively. (**D**) Molecular representation of Man5 glycan at N165 within NTD-B interacting with RBD-A. Multiple (1000) equally interspersed configurations (selected across all replicas) of the glycan at N165 from NTD-B are simultaneously shown. The glycan is represented as colored licorices (GlcNAc in blue, Man in green), whereas RBD-A and NTD-B are represented as cyan and red surfaces, respectively.

Altogether, the heterogeneous conformational dynamics disclosed by PCA and angle analysis, together with the reduced ACE2-binding responses revealed by biolayer interferometry experiments, show that the conformational plasticity of the RBD is affected by the absence of N-glycans at N165 and N234. Notably, two flexible linker loops, connecting the RBD with the receptor-binding C-terminal domain 2 (CTD2) located underneath, prime the RBD to undergo large conformational rearrangements, such as the ACE2-induced hinge motion.^58^ In this scenario, the subtle “up” /” down” equilibrium of the RBD can be dramatically altered by introducing single point mutations within the RBD or nearby regions,^57^ or also by varying pH conditions.^59^ Our work reveals that, similarly to MERS spike’s RBD,^60^ the SARS-CoV-2 spike’s RBD “up” state is metastable within its conformational ensemble. We find that the RBD necessitates two N-glycans located on the adjacent NTD, namely N165 and N234, to “load-and-lock” the “up” conformation and to successfully bind to ACE2 receptors. Although our simulations do not show a full “up-to-down” transition of the RBD, the investigated glycan-deleting mutations cause the RBD in the “up” state to experience a larger conformational freedom and to undergo wider tilting motions. As a consequence, this conformational instability might increase the vulnerability of the virus, as cryptic epitopes located on the RBD might become more exposed to antibody recognition,^61,62^ and, most likely, might decrease the virus infectivity as a result of a possible conformational shift of the RBD towards the closed state. Here, through biolayer interferometry experiments, we show that S protein binding to ACE2 is remarkably reduced for the N234A mutant, and slightly impaired for the N165A variant, whereas is completely abolished in the case of an engineered S protein with all three RBDs locked in the “down” state. As such, these experiments, confirming the computational predictions, advocate for a conformational shift of the RBD population towards the “down” state as a result of the depletion of N234 and N165 glycans. This is consistent with the dynamics of the Man9 glycan linked to N234, which we observe crawling towards the apical portion of the spike’s central axis, filling in the space left vacant by the RBD in the “up” conformation, and engaging in persistent interactions both with the same RBD “up” and the CH. Hence, it is not surprising that this glycan plays a larger role in stabilizing the RBD “up” than the glycan at N165, which is more solvent exposed and whose interaction pattern is less defined. However, we show that N165 steadily interacts with the RBD, therefore also contributing to loading and locking the “up” state, although to a lesser extent.

The SARS-CoV-2 spike’s marked sensitivity to mutations affecting its structural dynamics represents an opportunity for vaccine design,^32,63^ especially when it entails the alteration of the RBD “up” /” down” equilibrium.^57^ In this context, our findings pinpoint the possibility of controlling the RBD conformational plasticity by introducing N165A and N234A mutations. These N-linked sites – N234 in particular – are essential for locking the RBD in the open state, thus contributing to priming the SARS-CoV-2 virus to recognize ACE2 receptors and invade host cells. Excising these glycans might result in less infectious, or “weakened”, viruses bearing S proteins with predominantly “down” RBDs. This scenario would not prevent a humoral immune response targeting the upper side of the RBD,^64^ including the RBM, since the RBDs would still be able to transition to the “up” state, although to a lesser extent. Instead, it might stimulate a larger production of antibodies recognizing other known epitopes located either on the core of the RBD,^65,66^ or on the NTD,^67^ or other S2 regions,^68^ which might be engaged even in the closed state.

### Glycan Shield of the Full-Length SARS-CoV-2 S Protein

In the context of vaccine design, it is critical to consider all the strategies developed by viruses to evade the host immune response. Within this framework, many viruses use a glycan shield to mask the immunogenic epitopes targeted by neutralizing antibodies.^11,15^ SARS-CoV-2 S protein glycosylation is extensive and capable of thwarting the host humoral recognition.^11,15,69^ In addition to the spike head and RBD,^61,70^ another possible attractive target for antibodies is the stalk, as also shown in influenza virus studies.^71^ Whereas 19 N-glycans essentially camouflage the spike head region, only 3 N-glycosylation sites (N1158, N1174, and N1194) are present on the stalk. Exploiting the extensive sampling derived from all-atom simulations of our full-length model of SARS-CoV-2 S glycoprotein, we provide an unprecedented end-to-end overview of the spike’s glycan shield (**Figure 6**). In **Figure 6A**, each blue bush-like structure represents an ensemble of equally interspersed conformations of a single glycan sampled along a 1 μs trajectory for a total of 300 superimposed poses. Considering the different timescales of the dynamics of these N-glycans (nanoseconds), relative to the antibody/spike binding process (microseconds to milliseconds), this representation provides a realistic view of the shielding that molecules targeting the spike might encounter before binding. To quantify this shielding effect, we calculated the S protein’s accessible surface area (ASA) that is covered by glycans on both the head and stalk regions of the Open system. We also screened a wide range of probe radii, from 1.4 to 15 Å, to measure the effectiveness of the shield with respect to molecules of different sizes. A value of 1.4 Å is commonly set to approximate the radius of a water molecule, whereas larger radii can be used to describe larger moieties, ranging from small molecules at 2–5 Å to larger peptide- and antibody-sized molecules at 10–15 Å.^69,72^

**Figure 6.**
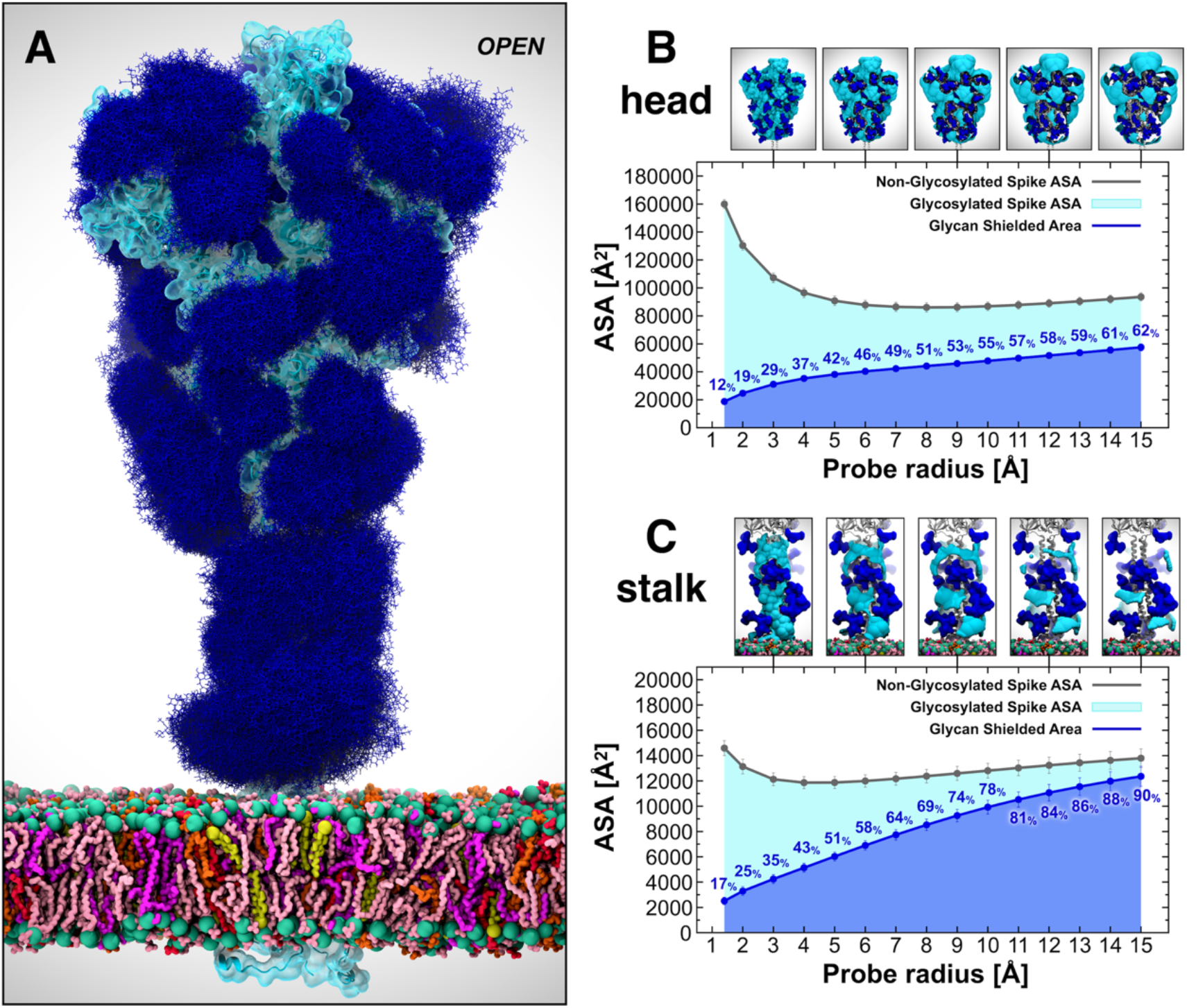
Glycan shield of the SARS-CoV-2 S protein. (**A**) Molecular representation of the Open system. Glycans at several frames (namely, 300 frames, one every 30 ns from one replica) are represented with blue lines, whereas the protein is shown with cartoons and highlighted with a cyan transparent surface. Color code used for lipid tails: POPC (pink), POPE (purple), POPI (orange), POPS (red), cholesterol (yellow). P atoms of the lipid heads are shown with green spheres. Cholesterol’s O3 atoms are shown with yellow spheres. (**B-C**) Accessible surface area of the head (**B**) and stalk (**C**) and the area shielded by glycans at multiple probe radii from 1.4 (water molecule) to 15 Å (antibody-sized molecule). The values have been calculated and averaged across all replicas of Open and are reported with standard deviation. The area shielded by the glycans is presented in blue (rounded % values are reported), whereas the gray line represents the accessible area of the protein in the absence of glycans. Highlighted in cyan is the area that remains accessible in the presence of glycans, which is also graphically depicted on the structure in the panels located above the plots.

Our results indicate that the head is overall less shielded by the glycans than the stalk at all probe radii (**Figures 6B** and **6C**). Interestingly, the stalk is almost completely inaccessible to large molecules such as antibodies, with glycan coverage equal to 90% of the protein accessible area for 15-Å-radius probe. On the contrary, the head represents an easier target as its glycan camouflaging is insufficient (62%) to cover its larger surface area, which includes one RBD in the “up” conformation. When smaller probes are screened (1.4–3 Å radius), the stalk and the head result to be similarly shielded at an average of 26% and 20%, respectively, suggesting that small molecules can equally penetrate either areas. ASA average and standard deviation values for the head and stalk in Open are provided in **Tables S6** and **S7**, respectively. Overall, taking glycosylation into account, the stalk appears as a potentially more difficult therapeutic target than the head, despite being a highly conserved domain among betacoronaviruses. Its smaller surface is well-protected by large sialylated and fucosylated tetrantennary glycans, which are found to be almost 100% effective in shielding large molecules. Interestingly, glycans at N1174 and N1198 have been found to always be tetrantennary, 100% fucosylated (N1174 and N1198), and 100% sialylated (N1198).^11,15^ However, small drugs might still interfere with the fusion process by binding to the HR2 domain of the stalk. In contrast, the head shows a higher vulnerability, which can be harnessed for vaccine development.

### Glycan Shield of the Receptor Binding Domain

As discussed in the previous section, the glycan shield plays a critical role in hiding the S protein surface from molecular recognition. However, to effectively function, the spike needs to recognize and bind to ACE2 receptors as the primary host cell infection route. For this reason, the RBM must become fully exposed and accessible.^73^ In this scenario, the glycan shield works in concert with a large conformational change that allows the RBD to emerge above the N-glycan coverage. Here, we quantify the ASA of the RBM within RBD-A, corresponding to the RBD/ACE2-interacting region (residues 400–508), at various probe radii in both the Open and Closed systems (**Figures 7A** and **7D**, full data in **Tables S8-S10**.). As expected, the ASA plots show a significant difference between the “down” (Closed) and “up” (Open) RBD conformations, with the RBM area covered by glycans being remarkably larger in the former. When RBD-A is in the “up” conformation, its RBM shows an average (across all radii) of only ∼9% surface area covered by glycans, compared with ∼35% in the Closed system (**Figures 7A** and **7D**). This difference is further amplified when considering a larger probe radius of 15 Å, with a maximum of 11% and 46% for Open and Closed, respectively. Interestingly, for smaller probes (1.4–3 Å) the shielding becomes weak in both systems, with an average of 6% and 16% for Open and Closed, respectively.

**Figure 7.**
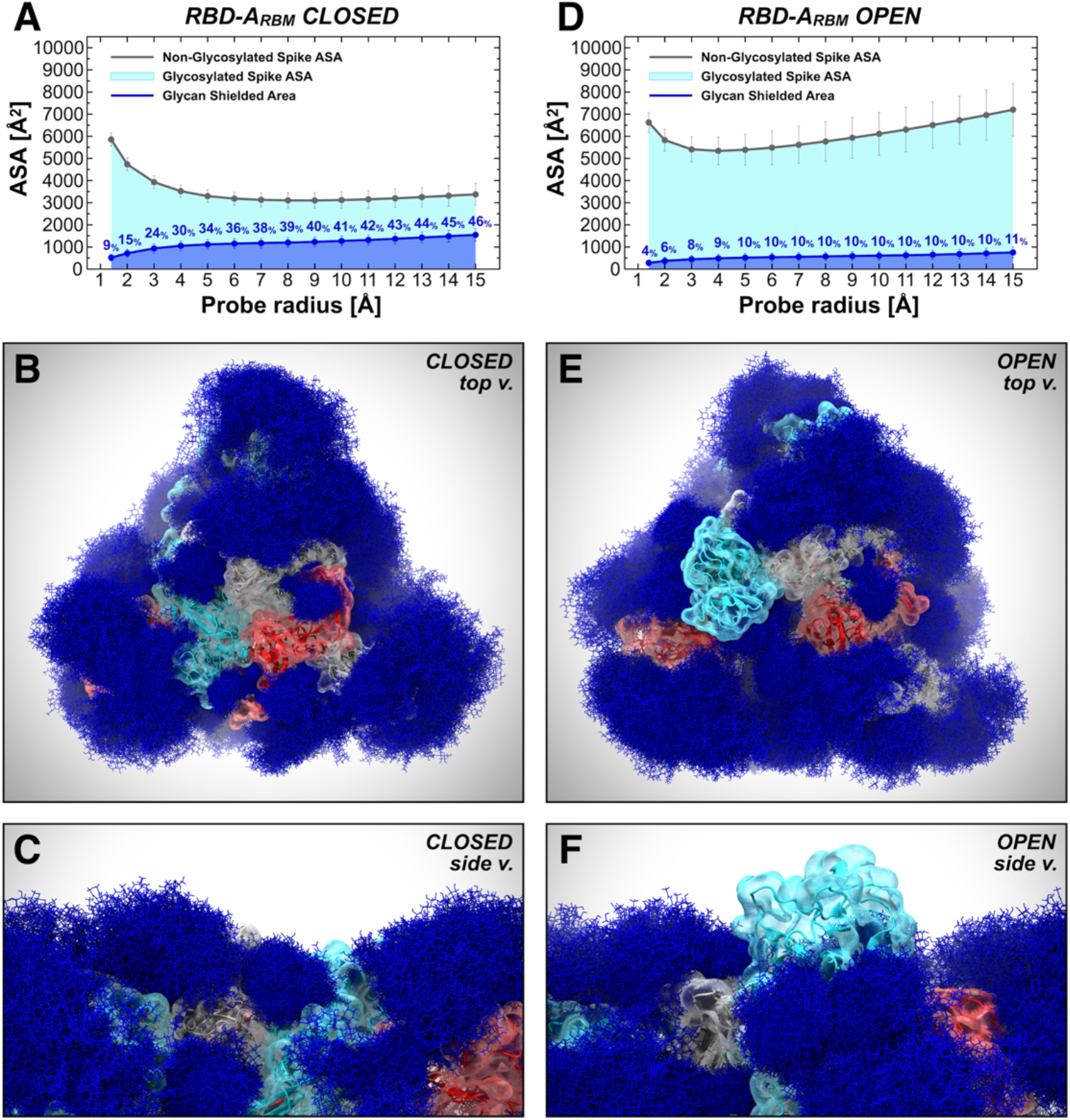
Glycan shield of the RBD ACE2-interacting region. The accessible surface area of the RBM-A and the area shielded by neighboring glycans in the Closed (**A**) and Open (**D**) systems are plotted at multiple probe radii from 1.4 (water molecule) to 15 Å (antibody-sized molecule). The values have been averaged across replicas and are reported with standard deviation. In blue is the area of the RBM-A covered by the glycans (rounded % values are reported), whereas the gray line is the accessible area in the absence of glycans. Highlighted in cyan is the RBM-A area that remains accessible in the presence of glycans, which is also graphically depicted on the structure in the panels located below the plots. (**B-F**) Molecular representation of Closed and Open systems from top (**B** and **E**, respectively) and side (**C** and **F**, respectively) views. Glycans (blue lines) are represented at several frames equally interspersed along the trajectories (300 frames along 0.55 ns for Closed and 1.0 μs for Open), while the protein is shown with colored cartoons and transparent surface (cyan, red and gray for chains A, B and C, respectively). Importantly, in panel **E** and **F**, RBD within chain A (cyan) is in the ‘up’ conformation and emerges from the glycan shield.

Note that the RBD region that does not directly interact with ACE2 remains shielded by the glycans in both “up” and “down” conformations (**Figure S10**). This region is equally protected regardless of the RBD conformation mostly owing to the presence of N-glycans bound to the RBD itself at N331 and N343 and to the N-glycans at N165 and N234 (**Figure S11**). Ultimately, this analysis shows that the RBM is always accessible when RBD is “up”, whereas it is very well-camouflaged when “down” (**Figure 7**). This suggests that the glycan shield of this critical domain is effectively paired with its “down-to-up” conformational change, allowing the RBM to transiently emerge from the glycan shield and bind to ACE2 receptors. Furthermore, while antibody targeting the RBD might be ineffective when the RBD is “down”, small molecules could more easily circumvent the glycan coverage. This is in agreement with structural data reporting the “up” conformation as a requirement for RBD neutralization by host antibodies.^74^ In this respect, several SARS-CoV-2 antibodies targeting the S protein have been identified (**Table S11**).^61,62,64–68,75–79^ The majority of these antibodies recognizes epitopes on the RBD, while only a few have been shown to address other antigenic regions within the NTD and CD (**Figure S12**). Among the RBD antibodies, B38 interacts with the RBM at the RBD/ACE2 interface,^64^ whereas S309 and CR3022 target the side/bottom part of the RBD.^61,62,65^ In addition, 4A8 and 1A9 have been found to engage with the NTD and CD, respectively.^67,68^ Analysis of the glycan shield effective on these epitopes is provided in **Section 3** of SI.

## Conclusions

Our work presents multiple microsecond-long, all-atom, explicitly solvated MD simulations of the full-length model of the glycosylated SARS-CoV-2 S protein embedded in a viral membrane. We show how the time-averaged glycan shield covers a vast amount of the S protein surface area and how it changes depending on the conformational state of the protein between open and closed states. Interestingly, our simulated models reveal a role – beyond shielding – for N-glycans at positions N165 and N234 as modulators of the RBD conformational plasticity. Simulations of the N165A/N234A and of N234A mutants of the S protein highlight their critical structural role in stabilizing the RBD “up” conformation. To confirm our simulation results, biolayer interferometry experiments conducted on N165A and N234A variants show a reduced ACE2 binding when these glycans are removed, revealing an RBD conformational shift towards the “down” state, with a larger effect for N234A. Overall, our work sheds new light on the full structure of this critical target and points to opportunities and challenges for small molecules and vaccine design. We make available through the NSF MolSSI and BioExcel COVID-19 Molecular Structure and Therapeutics site (https://covid.molssi.org/) our models to enable other groups to use and explore this dynamic system in atomic detail.^80^

## Supporting Information

The Supporting Information file (.pdf) contains the following: Material and methods, antibody accessibility analysis, additional simulations of the spike’s head model, supplementary figures and tables, captions of Movies S1-S3.

## Author Contributions

^+^L.C and Z.G. equally contributed to this work. R.E.A. and E.F. designed and oversaw the research project. L.C, Z.G., built the full-length spike models and performed the MD simulations. A.C.D. built the membrane bilayer. L.C., Z.G., R.E.A performed the simulation analyses. A.C.D. performed the analyses of the membrane bilayer. L.C., Z.G. created the figures. L.C. made movies S1 and S3. A.M.H., C.A.F., E.F. built the spike’s head model and performed analyses discussed in Section 3 of SI. A.M.H., C.A.F., E.F. made movie S2. J.S.M. designed and oversaw biolayer interferometry experiments. J.A.G. and C.K.H. performed the experiments and wrote the corresponding results and methods sections. L.C., Z.G., R.E.A., E.F. wrote the paper with contributions from all authors.

## Funding Sources

This work was supported by NIH GM132826, NSF RAPID MCB-2032054, an award from the RCSA Research Corp., a UC San Diego Moore’s Cancer Center 2020 SARS-COV-2 seed grant, and the Irish Research Council. LC is funded by a Visible Molecular Cell Consortium fellowship. This work was supported in part by NIH grant R01-AI127521 to J.S.M.

## Acknowledgements

We are grateful for the efforts of the Texas Advanced Computing Center (TACC) Frontera team and for the compute time made available through a Director’s Discretionary Allocation (made possible by the National Science Foundation award OAC-1818253), and also to the Irish Centre for High-End Computing (ICHEC) for computational resources. We thank Prof. Michael Feig and Dr. Lim Heo (Michigan State University), Prof. Syma Khalid (University of Southampton), Prof. Carlos Simmerling and his research group (SUNY Stony Brook), Prof. Ben Neuman (Texas A&M University), Prof. Greg Voth and Dr. Viviana Monje-Galvan (University of Chicago), Prof. Adrian Mulholland and his research group (University of Bristol), Prof. Julien Michel (University of Edinburgh) and his research group, Prof. Jean-Philip Piquemal (Sorbonne University), and Dr. Reda Rawi (NIH Vaccine Research Center) for system structure checks and helpful discussions.

## For Table of Contents Use Only

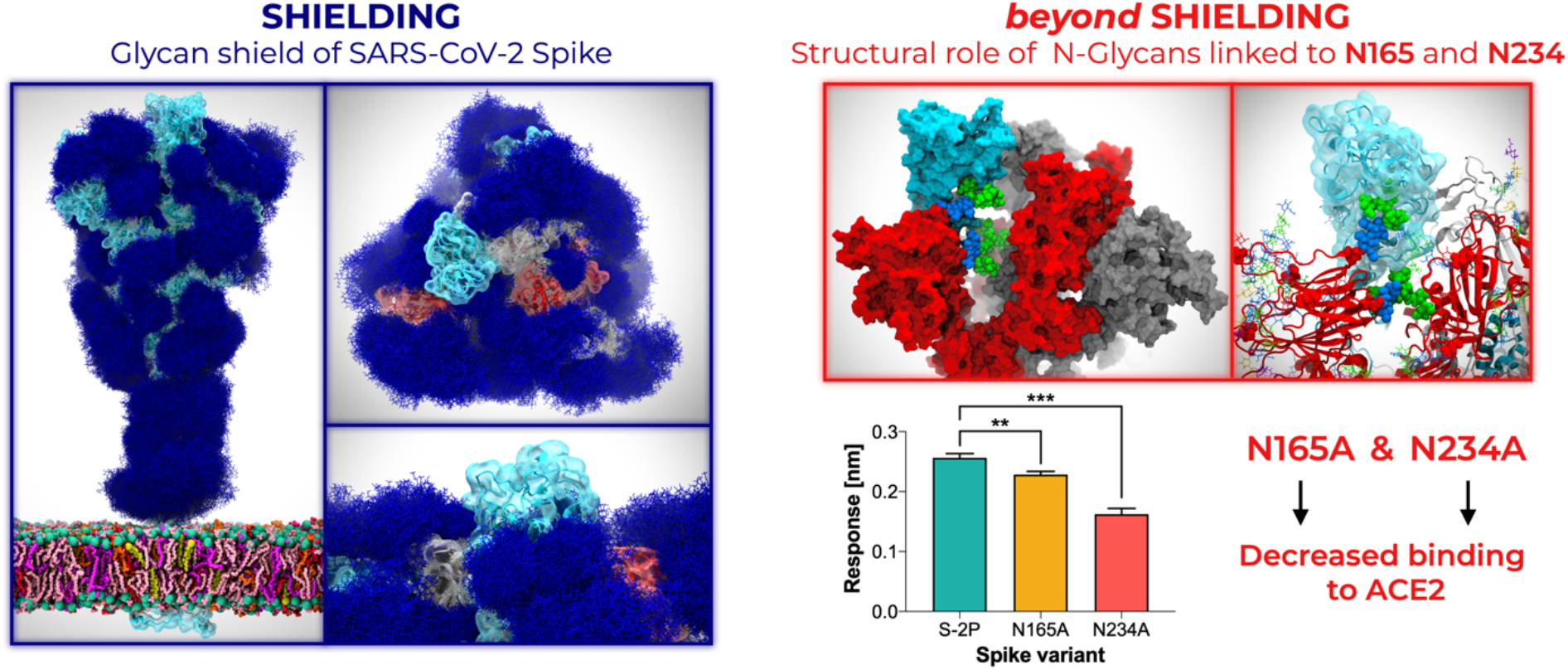

## Synopsis

The glycan shield is a sugary barrier that helps the viral SARS-CoV-2 spikes to evade the immune system. Beyond shielding, two of the spike’s glycans are discovered to prime the virus for infection.

## Notes

https://amarolab.ucsd.edu/covid19.php

https://covid.molssi.org

### Competing Interest Statement

The authors have declared no competing interest.

