## Supporting Information for "Beyond Shielding: The Roles of Glycans in SARS-CoV-2 Spike Protein"

**Table of Contents:**

1. **Materials and Methods**
   - **1.1.** Computational Methods
   - **1.2.** Experimental Methods
2. **Supplementary Simulations: All-Atom MD Simulations of S Protein Head using Amber FF14SB/Glycam06 Force Fields**
   - **2.1.** Computational Methods
   - **2.2.** Results and Discussion
   - **2.3.** Conclusions
3. **Overview of Neutralizing Antibody Epitope Accessibility**
4. **Supplementary Figures (S1 to S16)**
5. **Supplementary Tables (S1 to S15)**
6. **Supplementary Movies (S1 to S3)**
7. **Supplementary References**

### 1. Materials and Methods

#### 1.1. Computational Methods

*Model systems.* Severe acute respiratory syndrome coronavirus 2 (SARS-CoV-2) spike (S) glycoprotein is a large, glycosylated homotrimer, where each of its three identical monomers (residues 16–1273) can be divided into three main topological domains: the "head," comprising S1 and S2 subunits until residue 1140; the "stalk," composed of heptad repeat 2 (HR2) and transmembrane (TM) domains (residues 1141–1234); and the cytoplasmic tail (CT) (residues 1235–1273) (**Figure 1A** main text).[1] Experimental structures of SARS-CoV-2 S have been resolved in two main conformational states, open and closed, that were used in this study to build two complete, fully glycosylated models, referred to in this manuscript as "Open" and "Closed", respectively. The Closed system is based on a cryo-EM structure of the S protein solved at 2.80-Å average resolution (PDB ID: 6VXX),[2] where all receptor binding domains (RBDs) are in the "down" conformation. The Open system is instead built upon a cryo-EM structure of the S protein solved at 3.46-Å average resolution (PDB ID: 6VSB),[1] where only one RBD (chain A) is in the "up" conformation. A third system, called "Mutant", was also generated from the Open system upon mutation of N165 and N234 into alanine within all the three monomers, which ablated the respective N-glycan sequons. Although the cryo-EM structures of the S protein already provide critical information about its structure, they are usually incomplete and/or have been modified to increase protein stability.[1,3] For example, the introduction of two consecutive prolines (S-2P variant) in the central helix and/or of an engineered C-terminal foldon trimerization domain[4] has been adopted as a common strategy to stabilize the S protein for cryo-EM.[1,3] In addition, highly flexible protein regions (loops) and glycans beyond the first three sugars often remain unresolved owing to resolution limits. Therefore, several modeling steps were required to produce a full-length model of the wild type protein as described below.

*Missing loops modeling.* The employed cryo-EM structures of the S protein reveal several missing gaps corresponding to flexible loops ranging from 3 to 38 residues. To generate a complete construct, missing gaps were modeled as disordered loops using Modeller9.19.[5] Keeping the cryo-EM coordinates fixed, 50 models were independently generated for each monomer, from which the top models were selected and reassembled to recreate the full trimeric head. The alignment between the cryo-EM structure and the FASTA sequence of SARS-CoV-2 spike (QHD43416.1)[6] used by Modeller was generated using Clustal Omega.[7] The top models were further visually inspected to discard those in which loops were entangled in a knot or clashed with the rest of the structure. Finally, the stabilizing proline mutations from the cryo-EM structures were mutated back to wild type, and modeling artifacts were detected and corrected prior to simulations.

*HR2 and TM domain (stalk) modeling.* Both cryo-EM structures employed to build our models were stabilized using an engineered C-terminal foldon trimerization domain.[4] Therefore, the stalk region of the S protein from residues 1147–1234, including the HR2 and TM domains, had to be constructed. Using the Jpred4 server,[8] the secondary structure of the stalk sequence was predicted as three helical segments connected by two unstructured loops (**Figure S13**). Given the amphipathic nature of the helical segments, the three chains were assembled into an alpha-helical coiled-coil trimeric bundle using Modeller9.19.[5] A coiled-coil crystal structure, where the smaller and more hydrophobic residues are positioned inside the bundle and the polar residues are solvent-exposed, served as a template (PDB ID: 2WPQ).[9] Each alpha-helix was broken into three segments separated by two loops according to Jpred4 secondary structure predictions.

*CT modeling.* The CT of the S protein (residues 1235–1273) was modeled using the i-TASSER software.[10–12] i-TASSER generated five models with confidence scores of −1.25, −2.65, −3.15, −4.33, and −1.33 (C-score range [−5, 2]). Out of these, model 3 was selected because it revealed a helical domain between residues 1238 and 1245, where two cysteines were shown to be palmitoylated in another betacoronavirus, MHV-A59.[13,14] The corresponding cysteines in SARS-CoV-2 were C1240 and C1241. The remaining sequence of CT was predicted to be intrinsically disordered. Both cysteines (C1240 and C1241) were palmitoylated using lipid-tail functionality available within *Glycan Reader* in CHARMM-GUI.[15,16]

*Glycosylation.* SARS-CoV-2 S protein features 22 N-glycan sequons (N-X-S/T) per monomer, which have been found to be heterogeneously populated in different glycoanalytic studies.[17–19] Interestingly, two O-glycans have also been characterized at positions T323 and S325.[19] Our modeled constructs have been fully N-/O-glycosylated using the *Glycan Reader & Modeler* tool[20] integrated into *Glycan Reader*[15] in CHARMM-GUI.[16] An asymmetric (i.e., not specular across monomers) site-specific glycoprofile has been derived according to glycoanalytic data reported by Watanable et al.[17] for N-glycans and by Shajahan et al.[19] for O-glycans. Detailed per-chain descriptions of the site-specific glycoprofile of the S protein systems simulated in this work are shown in **Tables S1-S3**. In the Mutant system, N165A and N234A mutations were introduced to remove the respective N-glycans. This was performed with PSFGEN during system setup. In summary, 70 glycans ($22 \times 3_{monomers}$ N-glycans and $2_{chainA} + 1_{chainB} + 1_{chainC}$ O-glycans) have been added in the Open and Closed systems, whereas 64 have been added in the Mutant system. Our modeled N-glycans account for oligo-mannose from $Man_5GlcNAc_2$ to $Man_9GlcNAc_2$, complex and hybrid types, displaying one to four antennas. Additional modifications, such as fucosylation and sialylation, have also been site-specifically considered, as reported in Watanabe et al.[17] We remark that the oligosaccharides (GlcNAc-/$GlcNAc_2$-/$ManGlcNAc_2$-) originally solved in the cryo-EM structures have been generally retained or used as a basic scaffold, when possible, to build the full glycans. However, owing to steric clashes arising at particularly buried sites (for example, N122), a manipulation of glycan dihedrals and/or asparagine side chain has been necessary to fit in all the glycans.

*Membrane modeling.* The lipid composition of the membrane patch was selected based on the lipoprofiles[21,22] of the endoplasmic reticulum and trans-Golgi network, organelles in which the coronavirus membranes are known to be constructed.[23,24] A symmetric 225 Å × 225 Å lipid bilayer patch was generated using CHARMM-GUI's input generator.[16,25] The lipids were packed to an area per lipid of 70 $Å^2$ with the following ratio of phospholipids and cholesterol: POPC (47%), POPE (20%), CHL (15%), POPI (11%), and POPS (7%). The IUPAC names corresponding to these abbreviations are given in **Table S4**. The area per lipid value was selected based on the suggested CHARMM-GUI areas; the equilibrium values for this system were calculated and can be found in **Figure S5**.

*System preparation.* Upon functionalization (i.e., glycosylation and palmitoylation) of the glycoproteins through the *Glycan Reader* module available within CHARMM-GUI, further modifications to the structures were necessary and formatting issues were manually solved. Differently from SARS-CoV, where the S1/S2 site is cleaved prior to fusion (i.e., at the host cell surface), in SARS-CoV-2 the cleavage occurs when the virus is assembled.[2] Therefore, all the three constructs were modeled in their cleaved form, i.e. with the furin site cleaved between residues R685 and S686. However, for convenience, S1 and S2 subunits within each protomer have been assigned to the same chain (referred to as A/B/C), following the scheme used in 6VSB,[1] where RBD of chain A is in the "up" conformation. Glycans were attributed segnames from G1 to G70 (G1–G64 for Mutant), as reported in **Tables S1-S3**. Protonation states were assessed using PROPKA3[26] in the presence and absence of glycans, without registering any critical differences. The generated models were parametrized using PSFGEN and CHARMM36 all-atom additive force fields for protein, lipids, and glycans.[27,28] Parameters for palmitoylated cysteine were taken from Jang et al.,[29] whereas ions were treated using Beglov and Roux force fields.[30] The systems were fully solvated with explicit water molecules described using the TIP3P model.[31] The total number of atoms is 1,693,017 for the Open system (size: 225 Å × 225 Å × 367 Å), 1,658,797 for the Closed system (size: 225 Å × 225 Å × 359 Å), and 1,693,069 for the Mutant system (size: 225 Å × 225 Å × 367 Å).

*Molecular dynamics (MD) simulations.* All-atom MD simulations were conducted on the Frontera computing system at the Texas Advanced Computing Center (TACC) using NAMD 2.14.[32] The systems were initially relaxed through a series of minimization, melting (for the membrane), and equilibration cycles. During the first cycle, the protein, glycans, lipid heads (P atom for POPC, POPI, POPE, and POPS and O3 atom for CHL), solvent, and ions were kept fixed and the systems were subjected to an initial minimization of 10000 steps using the conjugate gradient energy approach. Subsequently, to allow the lipids tails to equilibrate, the temperature was incrementally changed from 10 to 310 K for 0.5 ns at 1 fs/step (NVT ensemble). The following simulation cycle was run at 2 fs/step, 1.01325 bar, and 310 K (NPT ensemble). Next, the systems were simulated with only the protein and glycans harmonically restrained at 5 kcal/mol to allow the full environment to relax in 2500 minimization steps and 0.5-ns simulations. Finally, all the restraints were released, and the systems were equilibrated for additional 0.5 ns. From this point, the production run was started, and frames were saved every 100 ps. Production MD simulations were run in triplicates for ~1 µs for Open and Mutant and ~0.6 µs for Closed (**Table S5**). To further explore the conformational space of the RBD in the "up" conformation, additional adaptive sampling simulations were run for Open and Mutant, which were also performed in triplicates for ~0.4 µs. Whereas velocities were randomly reinitialized, the initial coordinates were selected after principal component analysis (PCA) of RBD-A. In detail, the minimum (replica 4), mean (replica 5), and maximum (replica 6) along PC1 were identified and the corresponding frames used a starting point for adaptive sampling simulations.

All simulations were performed using periodic boundary conditions and particle-mesh Ewald[33] electrostatics for long-range electrostatic interactions with maximum grid spacing of 2 Å and evaluation every 3 time steps. Non-bonded van der Waals interactions and short-range electrostatic interactions were calculated with a cutoff of 12 Å and a switching distance of 10 Å. The SHAKE algorithm[34] was employed to fix the length of all hydrogen-containing bonds, enabling the use of 2-fs integration time steps. All simulations were performed under the NPT ensemble using a Langevin thermostat[35] (310 K) and a Nosé-Hoover Langevin barostat[36,37] (1.01325 bar) to achieve temperature and pressure control, respectively.

*Accessible surface area (ASA)*. ASA was calculated using the *measure sasa* command implemented in VMD,[38] which is based on the Shrake and Rupley algorithm,[39] in combination with in-house Tcl scripts. Three separate ASA analyses were conducted by taking into account the S protein head (residues 16–1140), stalk (residues 1141–1234), and receptor binding motif (RBM) of the RBD (residues 400–508), respectively. The area covered by glycans (i.e., the glycan shield) was obtained after the subtraction of the ASA of the considered domain in the absence of glycans with the ASA in the presence of glycans. This value was calculated along the trajectory with a stride of 150, 20, and 20 frames between each assessment for head, stalk, and RBM-A, respectively. For each system (Open and Closed), the values were averaged across all the respective replicas and standard deviation was computed. Apart from the standard 1.4-Å probe, this analysis was repeated for 14 different (1-Å-interspersed) values of probe radius (from 2 to 15 Å). Note that additional ASA analyses on the whole RBD-A (residues 330–530) and on its non-interacting region (residues 330–399 and 509–530) were also analogously performed (see SI). Similarly, epitope-specific ASA analyses were conducted on the chain A of Open and Closed systems using only 7.2 and 18.6 Å as probe radii. ASA evaluations were conducted with a stride of 20 frames. The residues considered for each epitope are listed in **Table S11**.

*Principal Component Analysis (PCA)*. PCA was performed using the sklearn.decomposition.PCA function in the *Scikit-learn* library using python3.6.9.[40] First, all simulations were aligned with *mdtraj*[41] onto the same initial coordinates using Cα atoms of chain-A central scaffold (residues 747–783 and 920–1032). Next, simulation coordinates of RBD-A (residues 330–530) from all systems (Open, Mutant, and Closed) and replicas were concatenated and used to fit the transformation function. Subsequently, the fitted transformation function was applied to reduce the dimensionality of each system simulation RBD-A Cα coordinates. Subsequently, the fitted transformation function was applied to reduce the dimensionality of RBD-A Cα coordinates from each system into the PC space. Note that it is important that the comparative PCA across systems have consistent eigen-basis of the principal components. This was ensured by transforming all systems coordinates into the same PC space. The same procedure was performed for PCA of RBD-B and RBD-C.

*Angles calculation.* The lateral angle and axial angle were calculated using in-house Tcl scripts along with VMD.[38] The axial angle is defined by three points corresponding to (i) the center of mass (COM) of RBD-A β-sheets (residues 394–403, 507–517, and 432–437), (ii - vertex) the COM of the central helices (residues 987–1032), and (iii) the COM of the top section of the central helices (residues 987–993). The lateral angle is described by three points corresponding to the (i) COM of RBD-A at frame 0, (ii - vertex) COM of the top section of the central helices, and (iii) COM of RBD-A β-sheets at frame *n*. We note that when calculating the COM of RBD-A we only considered the core residues defining the β-sheets of the domain, i.e. the most stable part of the RBD, discarding instead the highly flexible, solvent exposed loops (like the receptor binding motif) or the hinges that would have altered the position of the COM, thus biasing the angle calculation. In this way it was possible to keep track of the actual core motions of RBD-A. The other COMs were calculated on the central helices (CH). The CH are three alpha-helices (one for each monomer) located around the central axis of the spike trimer, representing the most rigid backbone of the spike's head as shown in Figure 1 of the main text.

Both angles were evaluated at each frame along the trajectories as a variation (positive or negative) with respect to their initial value. The trajectories were aligned by the S protein central scaffold (residues 747–783 and 920–1032) including the central helices using the coordinates at frame 0 as a reference. Importantly, whereas the axial angle was calculated in a three-dimensional space defined by *xyz* coordinates, the lateral angle was assessed by considering the projection of the COMs onto a two-dimensional space defined only by *xy* coordinates. In this way, the lateral angle only accounts for lateral tilt/shift of the RBD, discarding any other motion along *z*.

*Hydrogen bonds calculation.* Hydrogen bonds were calculated using the *measure hbonds* command implemented in VMD[38] in combination with in-house Tcl scripts. Hydrogen bonds criteria were set as 3.5 Å for distance between heavy atoms and as 45° for angle between Acc-Don-Hyd. All frames across all replicas were considered for this analysis. Occupancy (%) was determined by counting the number of frames in which a specific hydrogen bond was formed with respect to the total number of frames.

*Root-mean-square-deviation (RMSD)*. RMSD of protein Cα atoms was computed using the *measure rmsd* command implemented in VMD[38] in combination with in-house Tcl scripts. Different alignments were done before RMSD calculations using the initial coordinates of Cα atoms as a reference. In particular, for RBD-A RMSDs, Cα atoms of the S protein central scaffold (residues 747–783 and 920–1032) were used as a reference for alignment, whereas for the head, stalk, and CT RMSDs, the trajectories were aligned onto the Cα atoms of the residues of the respective regions.

*Root-mean-square-fluctuations (RMSF)*. RMSF was calculated using in-house python scripts along with *mdtraj*.[41] RMSF was computed for each glycan of every chain across all replicas in Closed, Open, and Mutant. The trajectories were aligned onto the initial coordinates using the Cα atoms of the entire protein as a reference.

**1.2 Experimental Methods**

*Protein expression and purification.* The spike S2P variant was expressed using a previously described mammalian expression vector containing proline substitutions at residues 986 and 987 and C-terminal 8xHis and TwinStrep tags.[1] Alanine substitutions were introduced into S2P to yield the N165A and N234A spike variants. To generate the disulfide-locked spike with all RBDs down, we introduced S383C, D985C substitutions into a previously described stabilized spike construct (HexaPro) containing proline substitutions at positions 817, 892, 899 942, 966 and 987).[42] ACE2 was expressed using a mammalian expression vector encoding residues 1-615 of human ACE2, a

C-terminal HRV 3C cleavage site, a mono-Fc tag, and 8xHis. Plasmids were transiently transfected into FreeStyle 293-F cells using polyethyleneimine and cultured for 4 days (for spike variants) or 6 days (for ACE2). Spike variants were purified by passing filtered cell supernatant over Strep-Tactin resin, and ACE2 was purified using Protein A agarose. ACE2 was subsequently cleaved with 3C protease passed over Protein A to remove the Fc-8xHis fragment.

*Biolayer interferometry.* Anti-foldon IgG was immobilized to an anti-human Fc (AHC) Octet biosensor (FortéBio), which was subsequently dipped into the specified spike ectodomain variant for loading. The biosensor was then dipped into 200 nM ACE2 to measure the RBD-spike association signal before being transferred to a well containing buffer only (10 mM HEPES pH 7.5, 150 mM NaCl, 3 mM EDTA, 0.05% Tween 20 and 1 mg/mL bovine serum albumin) to measure the dissociation signal. The total response at the end of the association phase was recorded for each variant and used to quantify the relative proportions of accessible RBDs. The total response at the end of the association phase, where nearly all the RBDs should be saturated, was recorded for each variant and used to quantify the relative proportions of accessible RBDs. Three independent experiments were run for each of the S2P, N165A, and N234A spike variants. Collected data shown in Figure 4B are reported as mean ± s.d., with the following values: 0.2563 nm ± 0.0070 nm (S2P); 0.2284 nm ± 0.0051 nm (N165A); 0.1623 nm ± 0.0097 nm (N234A). The experiment with the HexaPro variant with all the three RBDs locked in the closed conformation was used as control. Raw data are made available as supporting information.

### 2. Supplementary Simulations: All-Atom MD Simulations of S Protein Head using Amber FF14SB/Glycam06 Force Fields

#### 2.1. Computational Methods

The model of the SARS CoV2 S protein head was built in the open conformation by homology with SWISS MODEL[43] using the cryo-EM structure of the spike trimer in the open state as a template (PDB ID: 6VYB,[44] 3.2 Å resolution) and NCBI YP_009724390.1 as reference sequence. We note that this structure (6VYB) differs from to the one used as template for building the full-length, Open and Mutant models described in the main text (6VSB). In 6VYB, the RBD "up" belongs to chain B (corresponding to chain A in 6VSB), whereas chains C and A corresponds to chains B and C of 6VSB, respectively. The missing loops in the 6VYB cryo-EM structure were built automatically by SWISS MODEL based on structural libraries of backbone fragments from the PDB with similar sequences. The resulting protein structure exhibits 18 glycosylation sites per protomer, for a total of 54 sites per trimer that can be occupied. Glycosylation on these sites was built by aligning equilibrated structures of complex fucosylated (FA2B) and non-fucosylated (A2B) and of oligomannose (Man5 and Man9) from our in-house database[45] to the resolved GlcNAc residues in the cryo-EM structure. Because not all the GlcNAc residues were resolved in the cryo-EM and because two glycosylation sites per protomer are located in loops also not resolved in the cryo-EM structure, we have built the final 54-glycans model in two phases. In the first phase we built models with 46 glycosylation sites and run a 20 ns equilibration to obtain conformations of the rebuilt loops that allowed for the linking of our glycans. The chosen structures were then completed with glycosylation in all 54 sites, leading to the 54-glycans model.

In this additional set of simulations of the SARS-CoV-2 S protein head, we have considered three slightly different glycosylation profiles, shown in **Table S13,** resulting into three models that differs specifically at position N234, occupied either by a Man9 (Man9-N234) or where N234 is mutated into Ala (N234A) leading to glycan depletion, or where N234 is non-glycosylated. We ran two independent trajectories for the 46-glycans model (i.e., Man9-N234) and one for the related N234A mutant. Moreover, we performed one run each for the 54(53)-glycans models (i.e., Man9-N234, the N234A mutant, and non-glycosylated N234) for a total of 6 independent MD runs (see **Table S14**). We remark that the N234 N-linked site monitored in this set of simulation belongs to the NTD of chain C (NTD-C), corresponding to NTD-B of the full-length model described in the main text. In these setups of the spike head based on 6VYB and in the respective MD simulations, the protein was described using AMBER ff14SB parameters,[46] whereas the glycans by the GLYCAM06j-1 version of GLYCAM06.[47] An atmosphere of 200 mM NaCl was also included in all simulations, with ions represented by parameters in the AMBER ff14SB set. Water molecules were represented by the TIP3P model.[31] All simulations were run with v18 of the AMBER software package.[48]

The same system preparation and running protocol was used for all MD simulations. The energy of the system built as described above was minimized in two steps of 50,000 cycles of the steepest descent algorithm. During the first minimization all the heavy atoms were kept harmonically restrained using a potential weight of 5 kcal $mol^{-1}Å^2$, while the solvent, counterions and hydrogen atoms were left unrestrained. During the second minimization step only the protein heavy atoms were kept restrained, while the glycans, solvent, counterions and hydrogens were left unrestrained. After energy minimization the system was equilibrated in the NVT ensemble with the same restraint scheme, where heating was performed in two stages over a total time of 1 ns, from 0 to 100 K (stage 1) and then 100 to 300 K (stage 2). During equilibration the SHAKE algorithm was used to constrain all bonds to hydrogen atoms. The Van der Waals and direct electrostatic interactions were truncated at 11 Å and Particle Mesh Ewald (PME) was used to treat long range electrostatics with B-spline interpolation of order 4. Langevin dynamics with collision frequency of 1.0 $ps^{-1}$ was used to control temperature, which a pseudo-random variable seed to ensure there are no synchronization artefacts. Once the system was brought to 300 K an equilibration phase in the NPT ensemble of 1 ns was used to set the pressure to 1 atm. The pressure was held constant with isotropic pressure scaling and a pressure relaxation time of 2.0 ps. At this point all restraints on the protein heavy atoms were removed, allowing the system to evolve for 15 ns of conformational equilibration before production. The total simulation times, including equilibration, are shown in **Table S14**.

#### 2.2 Results and Discussion

As remarked in the main text, to further assess possible impact of the force fields and/or to the starting cryo-EM structure on the simulations described in the main text, we performed an additional set of simulations of the open SARS-CoV-2 S protein's head (presented here) using AMBER ff14SB/GLYCAM06j-1 force fields[46,47] and an alternative initial cryo-EM structure (PDB ID: 6VYB),[44] which presents the N234 GlcNAc in a slightly different orientation. The results of this set of simulations are described below.

**Man9-N234 model.** We analyzed the relative stability of the RBD domains in chains A, B and C during the MD production by calculating the backbone RMSD values relative to the starting homology model based on the 6VYB template. RMSD values were calculated using different alignments, namely residues 770 to 1255 of chain A, B and C, respectively, to evaluate potential biases. The differences in average RMSD values calculated from different alignments are within the standard deviation values used as error bars, therefore only results from the alignment to chain A (residues 770 to 1255) are shown in **Table S15**. Data were collected after 20 ns of equilibration.

Unless otherwise stated, results are shown for the 54(53)-glycans model systems. All simulations of the Man9-N234 model show that the glycan gradually inserts itself in the space left empty by the lifting-up of RBD-B (chain B in this model, corresponding to chain A in the full-length model) (**Movie S2**). This insertion progresses gradually through the formation of hydrogen bonding interactions between Man9 glycan at N234 and Y369 and N370 of RBD-C that evolve to reach the core of the spike's trimer defined by the location of D405, R408 and E409 of RBD-A, located in the diametrically opposite side across the spike apical center (see **Figure S14)**. Similar interactions have been registered also in the simulations of the full-length model of the spike described in the main text, where Man9 at N234 is found to establish persisent h-bonds with residues of RBD-C (RBD-A here).

The insertion of Man9 into the open pocket and its stable interactions with the trimer core's charged residues D405, R409 and E409 results in a stabilization of the whole structure, which is evident from the backbone RMSD analysis of the three RBD domains, shown in **Figure S15**. As an interesting note, position N357 in SARS-CoV S, corresponding to position N370 in SARS-CoV2 S, is part of an NST sequon and it is glycosylated.[49] The sequon is lost in SARS-CoV2 S with a mutation of the Thr to Ala. Because of the important and stable interactions of the Man9 at N234 with N370 along the reaction coordinate that allows it to reach the core of the trimer, it is reasonable to infer that glycosylation at position N370 in SARS-CoV would interfere with this process, potentially affecting or preventing the insertion of the Man9. To understand the significance of the presence of a large glycan such as Man9 at position N234 of NTD-C and its role in filling the empty space left by the opening of the RBD-B, we decided to remove it and to observe the conformational changes through another set of conventional MD simulations.

**N234A mutant and N234-nogly models.** To understand the role of the Man9 at N234 within NTD-C, we designed two models: a model with an N234A mutation in chain C (N234A) and a model where the N234 position also in chain C is non-glycosylated (N234-nogly). For those we analyzed four independent MD trajectories (see **Table S14)**, two from the 46(45)-glycans model, one from the 54(53)-glycans model sites in the N234A mutant form, and one from the 54(53)-glycans model with N234-nogly. In agreement with the full-length model of SARS-CoV2 S protein described in the main text, we observed a higher degree of dynamics of the open RBD relative to Man9-N234 when the glycan at N234 is missing. As an important note, here the N234A mutant and N234-nogly are missing Man9 only in one site of the trimer, i.e. chain C (chain B in the full-length model), which is a subtler modification relative to the full system in which the mutant is missing the glycans at N234 and N165 in each protomer. As indicated by the average backbone RMSD values shown in **Table S15**, except for the case of the 54(53)-glycans N234A model, for which the open RBD remains in a stable conformation for the length of the trajectory considered here, in the two simulations of the 46(45)-glycans N234A models and in the simulation of the 54(53)-glycans N234-nogly model, the open RBD domain is largely displaced relative to the original homology model, used as reference structure. In agreement with the full model discussed in the main text, the dynamics of the open RBD in the mutants is quite complex and within this simulation framework cannot (and should not) be defined as part of any specific reaction coordinate, such as a domain closing or unfolding. Nevertheless, in the simulations of the 46(45)-glycans N234A model, the open RBD (chain B) can be described as shifting towards the RBD of chain C, with the flexible loop interacting with the Man5 at N343 within chain C. Meanwhile, the conformational change we observed for the open RBD in the simulations of the 54(53)-glycans N234-nogly model corresponds more to a shift away from the RBD of chain C (**Figure S16)**.

#### 2.3. Conclusions

The simulations described in this section of the Supporting Information are presented here as supplementary material in support of the results discussed in the main manuscript based on much larger and complete 3D models of the SARS-CoV2 S glycoprotein. Indeed, despite the differences in the systems sizes, setups, original cryo-EM structure (6VYB vs. 6VSB), force field parameter sets (AMBER vs. CHARMM), MD software packages (Amber vs. NAMD), in the running protocols and details in the models' glycosylation, these two sets of simulations converge in showing that the absence of glycosylation at position N234 and at both positions N234 and N165, causes the open RBD to explore a larger conformational freedom, which may indicate a degree of instability. Therefore, all the simulations combined support the conclusion that the strangely "patchy" glycan shield of the SARS-CoV2 S glycoprotein may very well be engineered to play a structural role in supporting the active structure of the protein, by stabilizing the open or "up" conformation of the RBD. Also, as an important note, a mutation found in SARS-CoV2 S relative to the SARS-CoV S causes the loss of glycosylation at N370. Because of the dynamic process that Man9 follows in accessing the trimer core, which involves stable interactions with N370, a glycan at that location may interfere with insertion of the glycan at N234 into the pocket, which would remain empty, potentially making SARS-CoV S less stable in its open conformation than SARS-CoV2 S.

### 3. Overview of Neutralizing Antibody Epitope Accessibility

Several SARS-CoV-2 antibodies targeting the S protein have been identified (**Table S11**).[50–61] The majority of these antibodies recognize epitopes on the RBD, while only a few have been shown to address the antigenic regions within the NTD and CD (**Figure S12**). Among the RBD antibodies, B38 interacts with the RBM at the RBD/ACE2 interface,[52] whereas S309 and CR3022 target the side/bottom part of the RBD.[50,51,53] In addition, 4A8 and 1A9 have been found to engage with the NTD and CD, respectively.[55,56] To quantify the effects of glycan shielding on these epitopes, we calculated each epitope's ASA at two probe radii, 7.2 and 18.6 Å, which approximate the size of antibody hypervariable loop and variable fragments domains, respectively (**Figures S12A** and **S12B,** full data provided in **Table S12**).[62] In Open (RBD "up"), B38 epitope on the RBD/ACE-2 interface shows large ASA that is minimally shielded by glycans (10%/11%, for 7.2 and 18.6 Å probes, respectively ) (**Figure S12A**). Antibodies in this region exploit the vulnerability of the S protein when RBD is in the "up" conformation. Conversely, in Closed, the shielding of B38 epitope remarkably increases to 47%/62% (**Figure S12B**). When the RBD is in the "down" conformation, the RBM is buried by the other two neighboring RBDs, which already reduce its overall accessibility by ~40%. These values are in agreement with the RBM ASA trends shown in **Figure 7** in the main text.

The S309 epitope, located on the side of the RBD and near the N-glycan at N343, shows an interesting behavior. When including glycan N343 as a shielding factor, the epitope is covered up to 45%/56% of its total area. However, this glycan has been shown to be incorporated into the recognized epitope, which would considerably increase the antigenic region targeted by S309.[53] Interestingly, no substantial differences in shielding are observed between Open and Closed because this epitope is mostly located on the RBD side, which remains exposed even in the "down" conformation. Considering the bottom part of the RBD, the epitope recognized by the CR3022 antibody is found to be almost completely shielded in Open (69%/94%) and not accessible at all in Closed. This is in agreement with structural data showing that the cryptic epitope engaged by CR3022 is only available when the RBD is both “up” and rotated.[50,51] Remarkably, this epitope partially overlaps with VHH72, an antibody found to neutralize SARS-CoV-2 S pseudotyped viruses.[60] Finally, the 4A8 and 1A9 epitopes located within the NTD and CD, respectively, are not affected by the conformational changes of the RBD.[55,56] Whereas the epitope recognized by 4A8 is about 36%/51% shielded by glycans, the one targeted by 1A9 is almost completely covered at 86%/99%. These results probe further questions on 4A8 and 1A9 binding mode in the presence of glycans.

### 4. Supplementary Figures

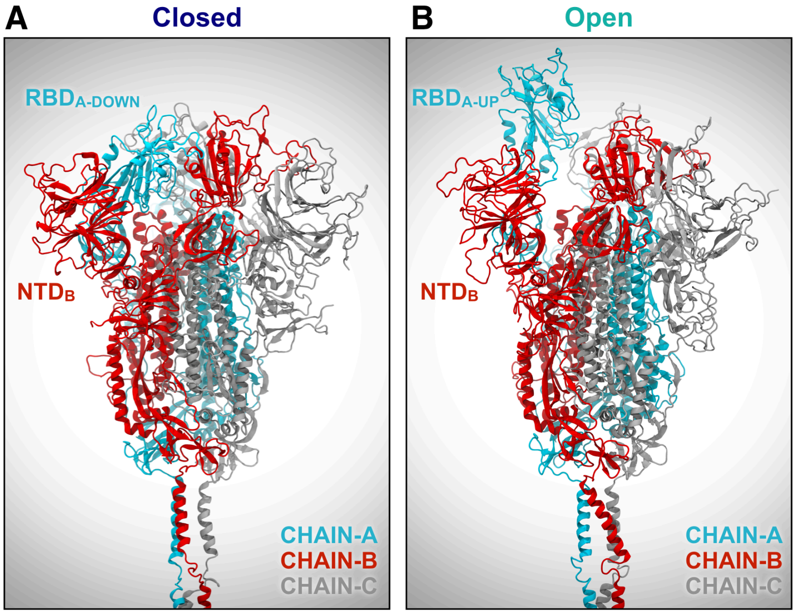

**Figure S1.** Molecular representation of S protein ectodomain in the Closed (A) and Open (B) systems. Protein is shown with cartoons, where chains A-B-C are colored in cyan, red and silver, respectively. Glycans are omitted for clarity**.**

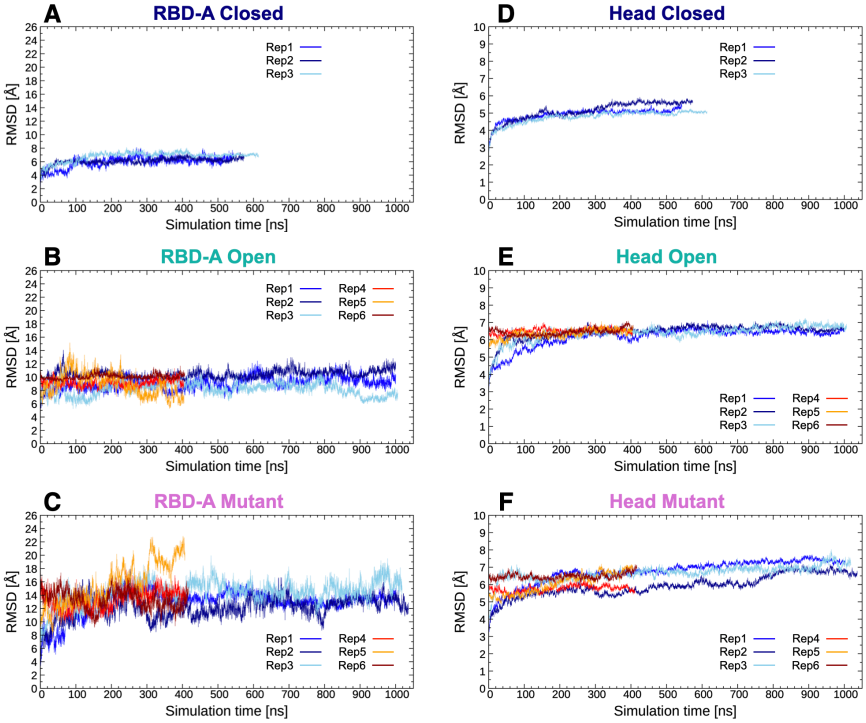

**Figure S2.** RMSD [Å] vs. time [ns] plots of the receptor binding domain of chain A (**A-C**) (RBD-A; residues 330-530) and the head (**D-F**) (residues 16-1140) of the S protein Cα atoms in the Closed, Open, and Mutant systems along each replica. For the RBD-A RMSDs, Cα atoms of the S protein central scaffold (residue 920 to 1032, 747 to 783) was used for alignment, while for head RMSDs the S protein was aligned onto all the head Cα atoms.

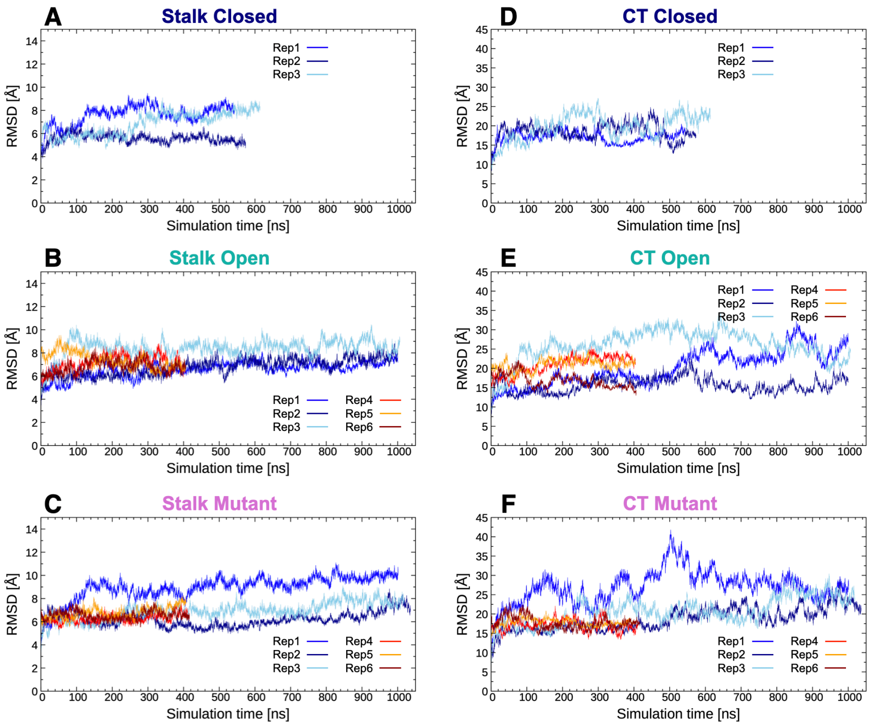

**Figure S3.** RMSD [Å] vs. time [ns] plots of the stalk (**A-C**) (residues 1141-1234) and cytoplasmic tail (**D-F**) (residues 1235-1273) Cα atoms of the S protein in the Closed, Open, and Mutant systems along each replica. For the stalk RMSDs, the stalk Cα atoms were used for alignment. Similarly, the cytoplasmic tail was aligned onto its Cα atoms prior generation of respective RMSDs.

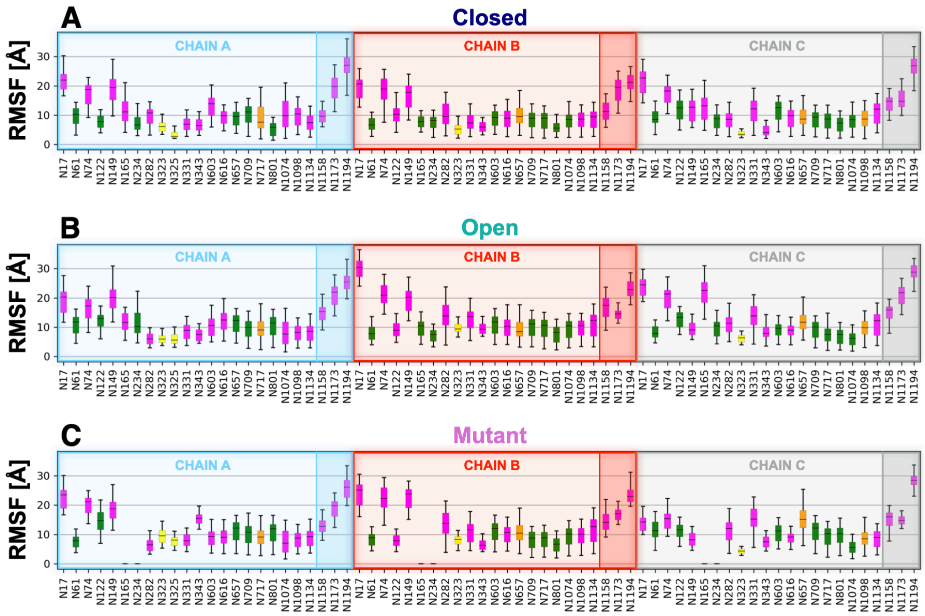

**Figure S4**. RMSF [Å] of each glycan for all chains across all simulations in the Closed (A), Open (B), and Mutant (C) systems. Glycans are colored based on their structure and composition: complex glycans in magenta, oligomannose glycans in green, hybrid glycans in orange, and O-glycans in yellow. Glycans in each chain are decomposed by domain: head (N17 to N1134), and stalk (N1158 to N1194), highlighted with different level of background opacity.

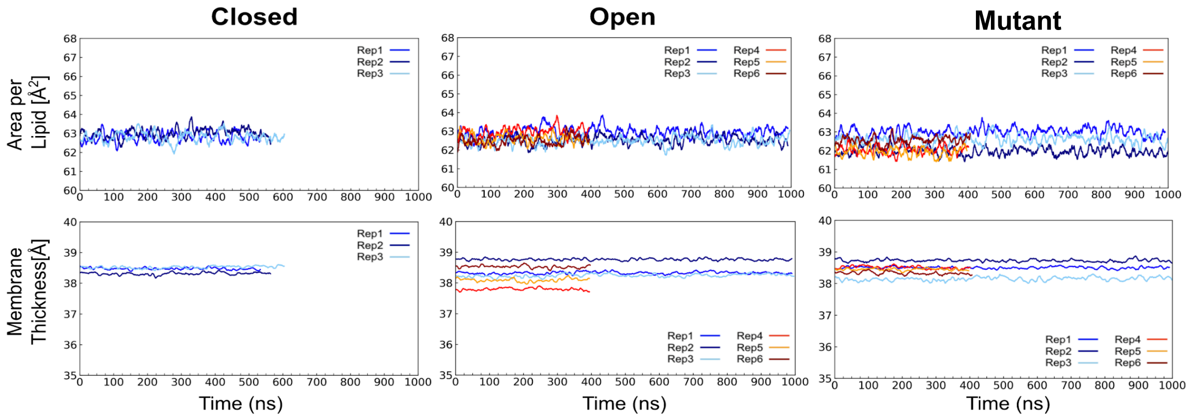

**Figure S5.** Plots of equilibrium area per lipid (top row) and P-P distance indicating membrane thickness (bottom row) of the membranes for the Closed (left), Open (center), and Mutant (right) systems along with each replicate.

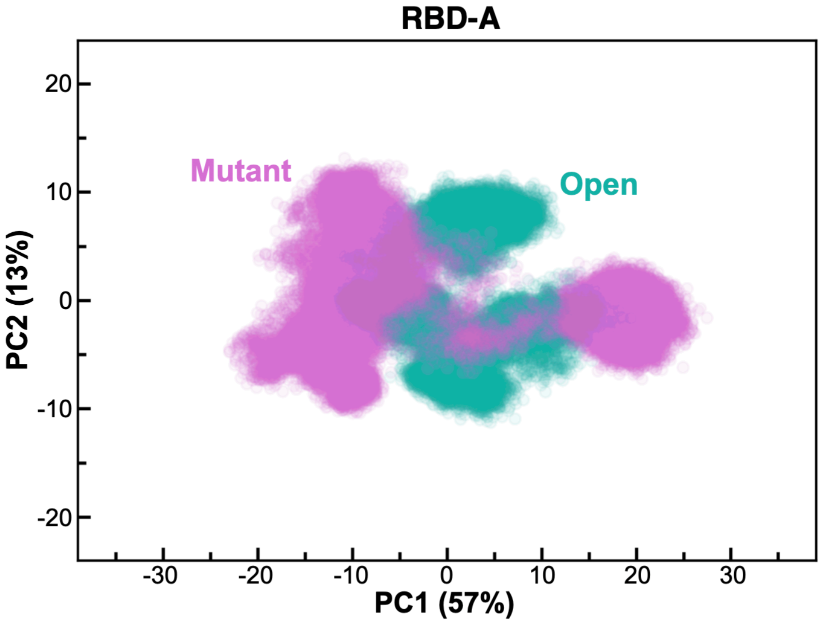

**Figure S6.** PCA plot showing PC1 vs PC2 of RBD-A (residues 330-530) in Open and Mutant in teal and magenta, respectively. The amount (%) of variance accounted by each PC is shown between parentheses.

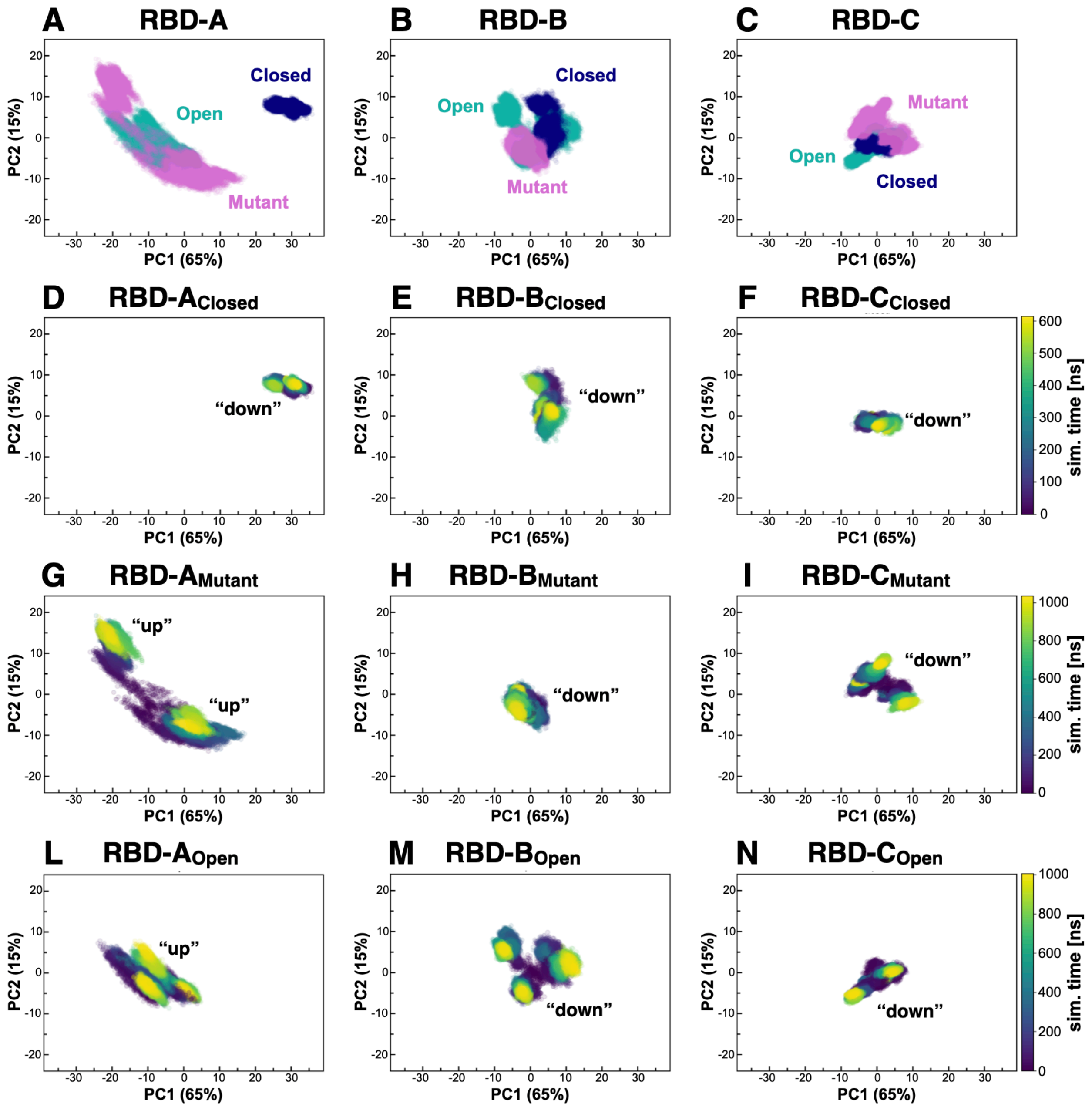

**Figure S7. (A-C)** PCA plots showing PC1 vs PC2 of RBD-A, RBD-B, and RBD-C (residues 330-530) in Closed, Open, and Mutant in blue, teal, and magenta, respectively. (**D-N**) PCA plots showing each system's time evolution for each replica. The time scale is reported as a color bar for each system. We remark that for Open and Mutant, replicas 4, 5, and 6 were simulated for ~400 ns. The RBD state is annotated within each respective plot as "up" or "down." The amount (%) of variance accounted by each PC is shown between parentheses.

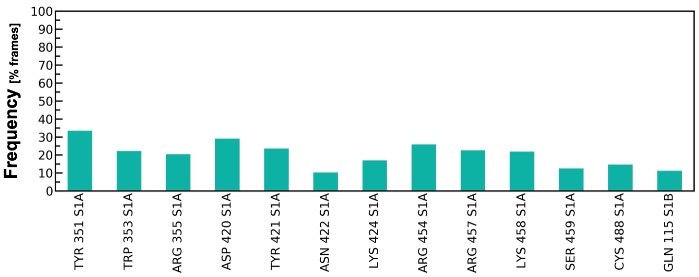

**Figure S8.** The main hydrogen bond interactions of N-glycans at N165 within the Open system are shown as occupancy across all replicas (% frames).

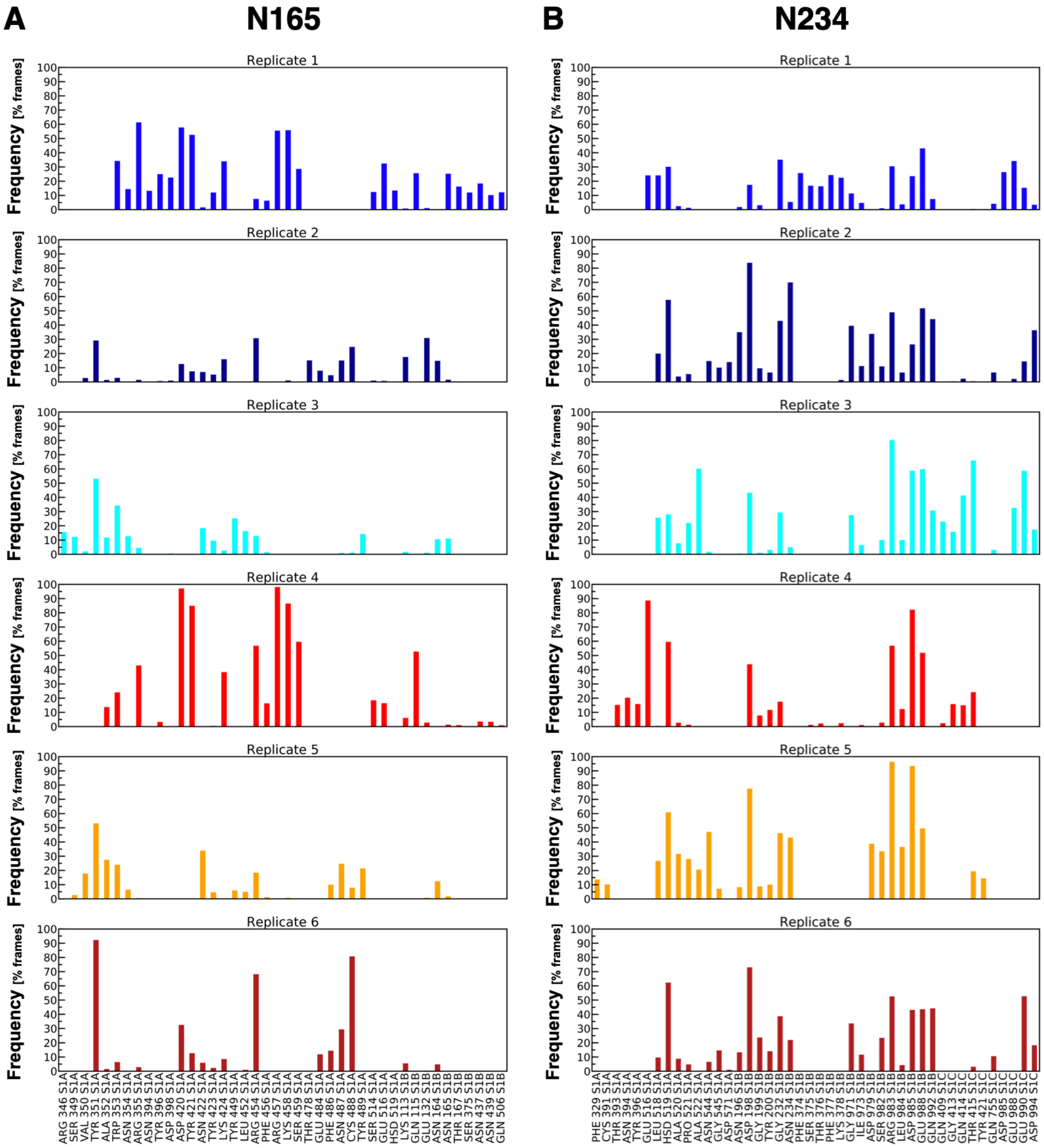

**Figure S9.** Main hydrogen bond interactions of glycan N165 (**A**) and N234 (**B**) within each replica of the Open system are shown as occupancy ($y$, % frames). Residues interacting with the glycans are reported on the $x$ axis.

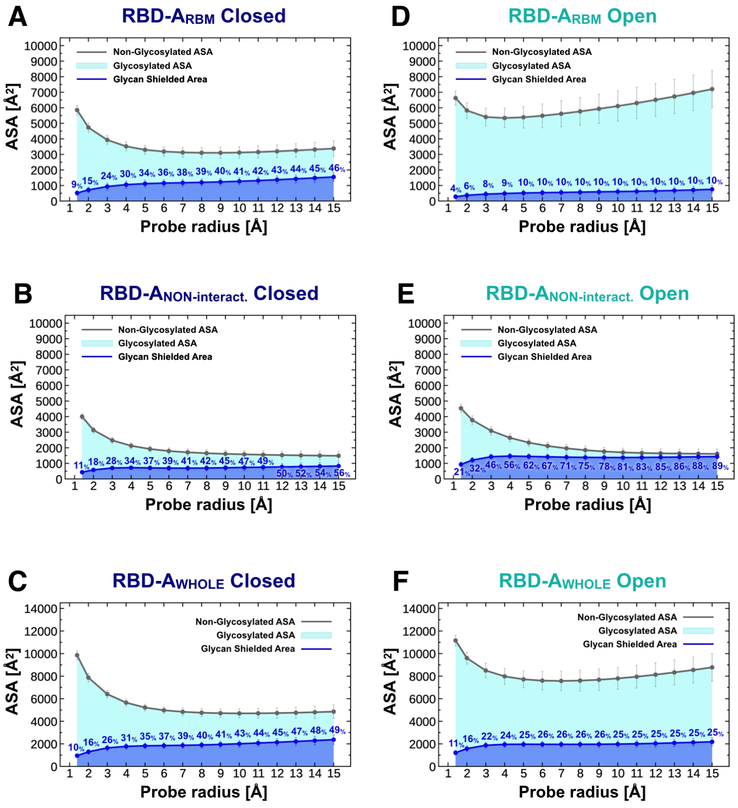

**Figure S10.** The accessible surface area of the RBD-ARBM (residues 400 to 509), RBD-ANON-INTERACTING (residues 330 to 399 and 509 to 530), RBD-AWHOLE (residues 330 to 530) and the area shielded by neighboring glycans in the Closed (**A-C**, respectively) and Open (**D-F**, respectively) systems is plotted at multiple probe radii from 1.4 Å (water molecule) to 15 Å. The values have been averaged across replicates and are reported with standard deviation. In blue is the area covered by the glycans (rounded % are reported), while the grey line is the accessible area in the absence of glycans. Highlighted in cyan is the area that remains accessible in the presence of glycans, which is also graphically depicted on the structure in the panels located below the plots.

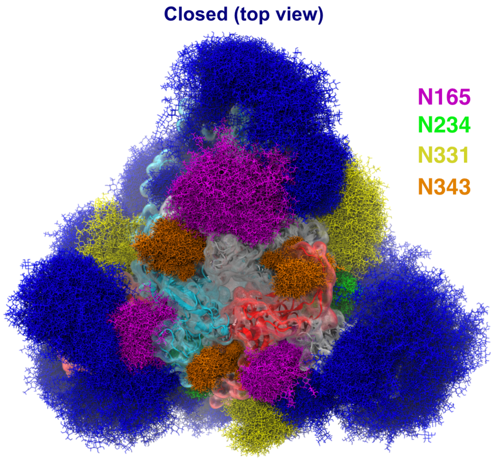

**Figure S11.** Molecular representation of the Closed system from top view. Glycans (blue lines) are represented at several frames equally interspersed along the trajectories (300 frames along 0.55 ns for Closed and 1.0 us for Open), while RBD-A is depicted with cyan cartoons and transparent surface. Chain B and Chain C are shown in red and grey cartoons, respectively, and transparent surface. Glycans at N165, N234, N331 and N343 are colored in purple, green, yellow, and orange, respectively,

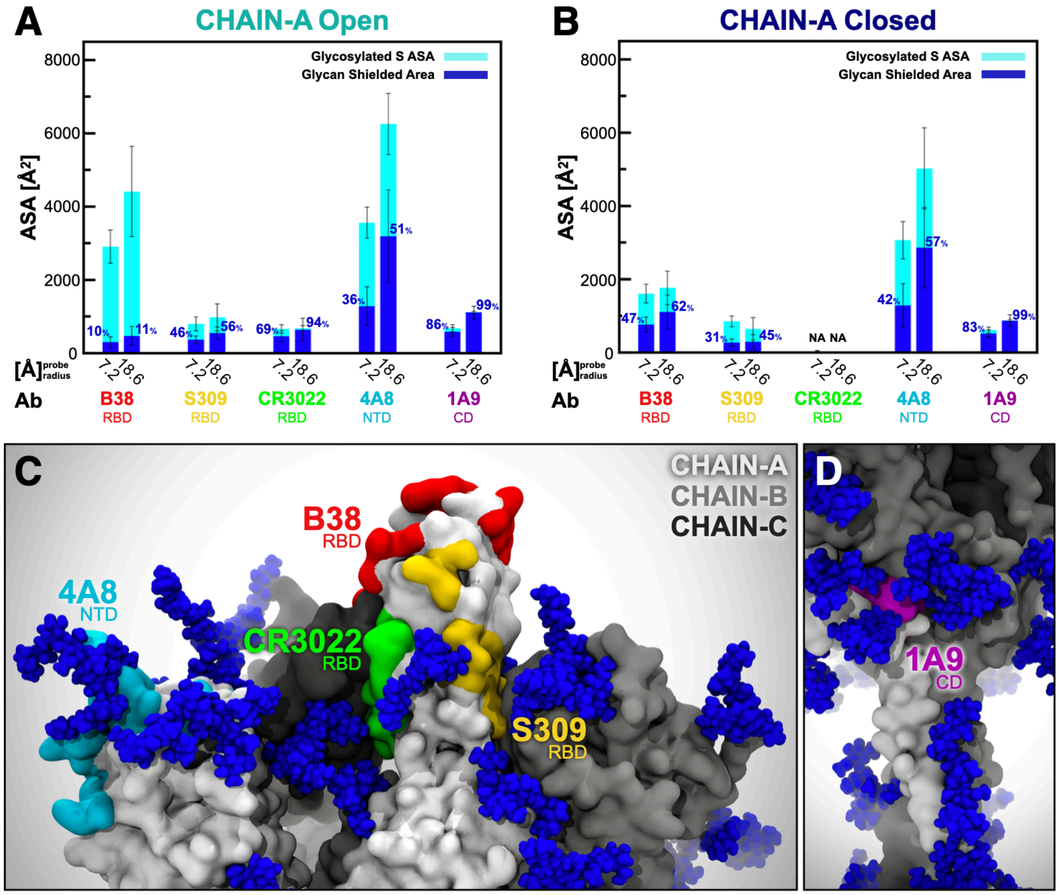

**Figure S12. Accessibility of neutralizing antibody epitopes.** The accessible surface area of antibody epitopes and the area shielded by neighboring glycans are plotted at probe radii 7.2 and 18.6 Å for chain A in Open (**A**) and Closed (**B**) systems. The area of the protein covered by the glycans is depicted in blue (rounded % values are reported), whereas highlighted in cyan is the epitope area that remains accessible in the presence of glycans. The values have been averaged across replicas and are reported with standard deviation. (**C**) Side view of the top region of S protein in the Open system, where the antibody epitopes are highlighted following the color scheme used in panels A and B. (**D**) Magnified view of 1A9 epitope (highlighted in purple) located within CD.

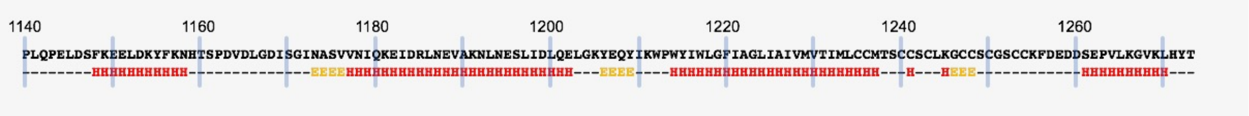

**Figure S13.** Secondary structure of stalk and cytosolic tail regions of the S protein as predicted with Jpred4 server. The secondary structure of the stalk sequence was predicted as three helical segments ("H", red) connected by two unstructured loops.

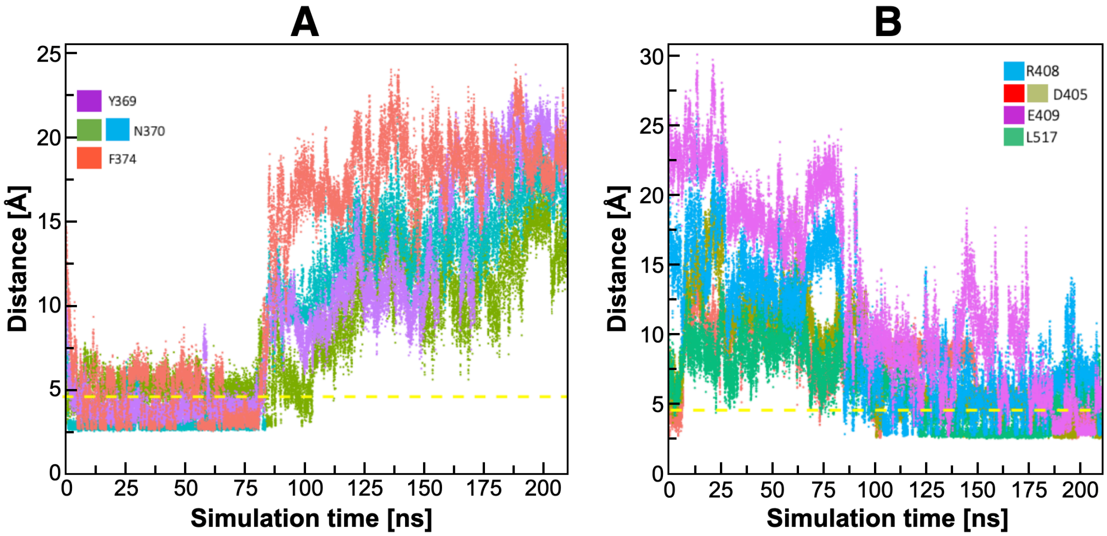

**Figure S14.** (**A-B**) Hydrogen bonding distances (Å) calculated through the 210 ns trajectories of the 46-glycans Man9-N234 spike chosen here for the higher sampling time. (**A**) Interactions with Y369, N370, F374 belonging to RBD-C. (**B**) Interactions with R408, D405, E409, L517 belonging to RBD-A. Interactions with F374 and L517 involve backbone atoms and double colors indicates distances between different atoms on the same residues. Similar hydrogen bonding pattern and behavior have been observed in all other simulations of the Man9-N234 systems.

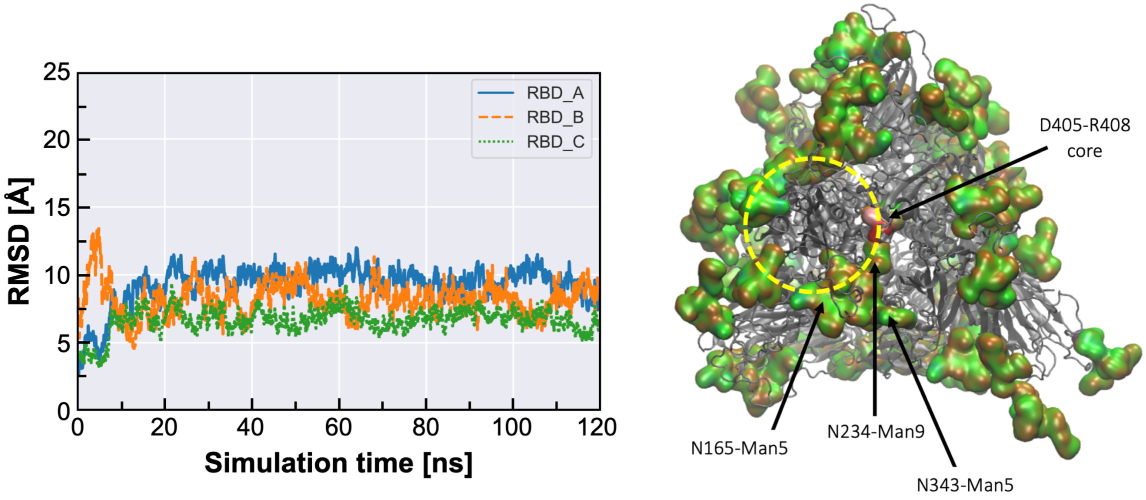

**Figure S15** *Left panel*: Backbone RMSD values calculated for the 54-glycans Man9-N234 system over 120 ns of production. A structural alignment of the protein backbone atoms was done on the stalk residues (resid 770 to 1255) of chain A, see text for more details. *Right panel*: Snapshot of the MD simulation (at 44 ns) showing the insertion of the Man9 at N234 deep into the trimer core highlighted by the residues in red (D405) and white (R408) with labels indicating other important glycans in framing the open conformation of the "up" RBD, highlighted within the yellow circle. All glycans are rendered as Quick Surface (green C atoms) and protein as New Cartoon (all grey). Rendering done with VMD and graphs with *seaborn.pydata.org*.

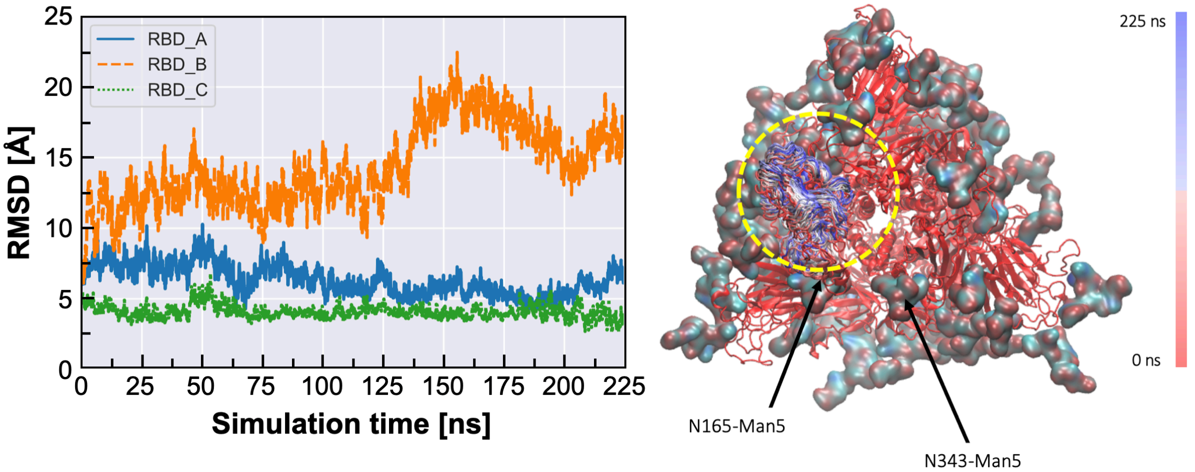

**Figure S16.** *Left panel*: Backbone RMSD values calculated for the 54-glycans N234-nogly system over 225 ns of production. A structural alignment of the protein backbone atoms was done on the stalk residues (resid 770 to 1255) of chain A, see text for more details. *Right panel*: Snapshot from the MD simulation showing a static representation of the protein (red New Cartoon) and the change in the relative position of the open RBD domain (resid 437 to 508) along the trajectory. The coloring indicates the trajectory progression as indicated in the bar on the right-hand side. The yellow circle highlights the position of the open RBD domain. All glycans are shown with Quick Surface (with C atoms in cyan). Rendering done with VMD and graphs with *seaborn.pydata.org*.

### 5. Supplementary Tables

**Table S1**. Glycan compositions for chain-A.

CHAIN A

| # | SITE | TYPE | STRUCTURE | SEQUENCE |
|---|---|---|---|---|
| G1 | N17 | FA2 | 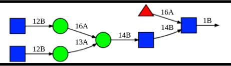 | bDGlcNAc(1→2)aDMan(1→6)[bDGlcNAc(1→2)aDMan(1→3)]bDMan(1→4)bDGlcNAc(1→4)<br>[aLFuc(1→6)]bDGlcNAc(1→)PROA-17 |
| G2 | N61 | M5 | 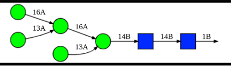 | aDMan(1→6)[aDMan(1→3)]aDMan(1→6)<br>[aDMan(1→3)]bDMan(1→4)bDGlcNAc(1→4)bDGlcNAc(1→)PROA-61 |
| G3 | N74 | A3 | 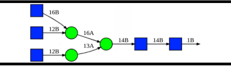 | bDGlcNAc(1→6)[bDGlcNAc(1→2)]aDMan(1→6)<br>[bDGlcNAc(1→2)aDMan(1→3)]bDMan(1→4)bDGlcNAc(1→4)bDGlcNAc(1→)PROA-74 |
| G4 | N122 | M5 | 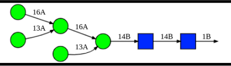 | aDMan(1→6)[aDMan(1→3)]aDMan(1→6)<br>[aDMan(1→3)]bDMan(1→4)bDGlcNAc(1→4)bDGlcNAc(1→)PROA-122 |
| G5 | N149 | FA2G2S1 | 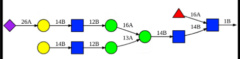 | aDNeu5Ac(2→6)bDGal(1→4)bDGlcNAc(1→2)aDMan(1→6)<br>[bDGal(1→4)bDGlcNAc(1→2)aDMan(1→3)]bDMan(1→4)bDGlcNAc(1→4)<br>[aLFuc(1→6)]bDGlcNAc(1→)PROA-149 |
| G6 | N165 | FA2G2S2 | 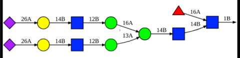 | xaDNeu5Ac(2→6)bDGal(1→4)bDGlcNAc(1→2)aDMan(1→6)<br>[aDNeu5Ac(2→6)bDGal(1→4)bDGlcNAc(1→2)aDMan(1→3)]bDMan(1→4)bDGlcNAc(1→4)<br>[aLFuc(1→6)]bDGlcNAc(1→)PROA-165 |
| G7 | N234 | M8 | 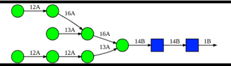 | aDMan(1→2)aDMan(1→6)[aDMan(1→3)]aDMan(1→6)<br>[aDMan(1→2)aDMan(1→2)aDMan(1→3)]bDMan(1→4)bDGlcNAc(1→4)bDGlcNAc(1→)PROA-234 |
| G8 | N282 | FA3 | 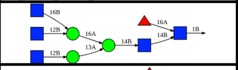 | bDGlcNAc(1→6)[bDGlcNAc(1→2)]aDMan(1→6)<br>[bDGlcNAc(1→2)aDMan(1→3)]bDMan(1→4)bDGlcNAc(1→4)[aLFuc(1→6)]bDGlcNAc(1→)PROA-282 |
| G9 | N331 | FA2 | 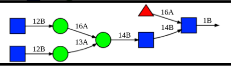 | bDGlcNAc(1→2)aDMan(1→6)[bDGlcNAc(1→2)aDMan(1→3)]bDMan(1→4)bDGlcNAc(1→4)<br>[aLFuc(1→6)]bDGlcNAc(1→)PROA-331 |
| G10 | N343 | FA2 | 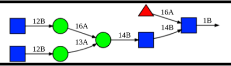 | bDGlcNAc(1→2)aDMan(1→6)[bDGlcNAc(1→2)aDMan(1→3)]bDMan(1→4)bDGlcNAc(1→4)<br>[aLFuc(1→6)]bDGlcNAc(1→)PROA-343 |
| G11 | N603 | FA2 | 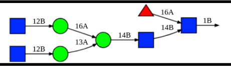 | bDGlcNAc(1→2)aDMan(1→6)[bDGlcNAc(1→2)aDMan(1→3)]bDMan(1→4)bDGlcNAc(1→4)<br>[aLFuc(1→6)]bDGlcNAc(1→)PROA-603 |
| G12 | N616 | A2 | 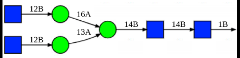 | bDGlcNAc(1→2)aDMan(1→6)<br>[bDGlcNAc(1→2)aDMan(1→3)]bDMan(1→4)bDGlcNAc(1→4)bDGlcNAc(1→)PROA-616 |
| G13 | N657 | M5 | 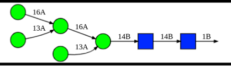 | aDMan(1→6)[aDMan(1→3)]aDMan(1→6)<br>[aDMan(1→3)]bDMan(1→4)bDGlcNAc(1→4)bDGlcNAc(1→)PROA-657 |
| G14 | N709 | M6 | 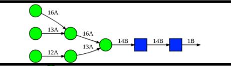 | aDMan(1→6)[aDMan(1→3)]aDMan(1→6)<br>[aDMan(1→2)aDMan(1→3)]bDMan(1→4)bDGlcNAc(1→4)bDGlcNAc(1→)PROA-709 |
| G15 | N717 | Hybrid G1 | 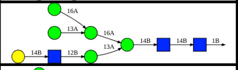 | bDGal(1→4)bDGlcNAc(1→2)aDMan(1→3)[aDMan(1→6)<br>[aDMan(1→3)]aDMan(1→6)]bDMan(1→4)bDGlcNAc(1→4)bDGlcNAc(1→)PROA-717 |
| G16 | N801 | M6 | 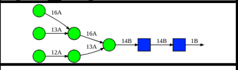 | aDMan(1→6)[aDMan(1→3)]aDMan(1→6)<br>[aDMan(1→2)aDMan(1→3)]bDMan(1→4)bDGlcNAc(1→4)bDGlcNAc(1→)PROA-801 |
| G17 | N1074 | FA2G2S1 | 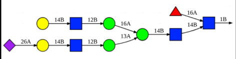 | aDNeu5Ac(2→6)bDGal(1→4)bDGlcNAc(1→2)aDMan(1→3)<br>[bDGal(1→4)bDGlcNAc(1→2)aDMan(1→6)]bDMan(1→4)bDGlcNAc(1→4)<br>[aLFuc(1→6)]bDGlcNAc(1→)PROA-1074 |
| G18 | N1098 | FA2 | 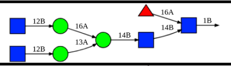 | bDGlcNAc(1→2)aDMan(1→6)[bDGlcNAc(1→2)aDMan(1→3)]bDMan(1→4)bDGlcNAc(1→4)<br>[aLFuc(1→6)]bDGlcNAc(1→)PROA-1098 |
| G19 | N1134 | FA1 | 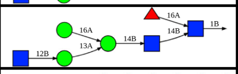 | bDGlcNAc(1→2)aDMan(1→3)[aDMan(1→6)]bDMan(1→4)bDGlcNAc(1→4)<br>[aLFuc(1→6)]bDGlcNAc(1→)PROA-1134 |
| G20 | N1158 | A2 | 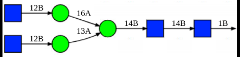 | bDGlcNAc(1→2)aDMan(1→6)<br>[bDGlcNAc(1→2)aDMan(1→3)]bDMan(1→4)bDGlcNAc(1→4)bDGlcNAc(1→)PROA-1158 |
| G21 | N1173 | FA4 | 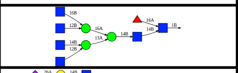 | bDGlcNAc(1→6)[bDGlcNAc(1→2)]aDMan(1→6)[bDGlcNAc(1→4)<br>[bDGlcNAc(1→2)]aDMan(1→3)]bDMan(1→4)bDGlcNAc(1→4)[aLFuc(1→6)]bDGlcNAc(1→)PROA-1173 |
| G22 | N1194 | FA4G4S1 | 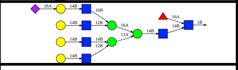 | aDNeu5Ac(2→6)bDGal(1→4)bDGlcNAc(1→6)[bDGal(1→4)bDGlcNAc(1→2)]aDMan(1→6)<br>[bDGal(1→4)bDGlcNAc(1→4)<br>[bDGal(1→4)bDGlcNAc(1→2)]aDMan(1→3)]bDMan(1→4)bDGlcNAc(1→4)<br>[aLFuc(1→6)]bDGlcNAc(1→)PROA-1194 |
| G23 | T323 | O-glycan | 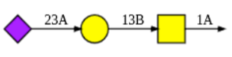 | aDNeu5Ac(2→3)bDGal(1→3)aDGalNAc(1→)PROA-323 |
| G24 | S325 | O-glycan | 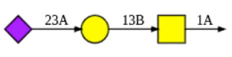 | aDNeu5Ac(2→3)bDGal(1→3)aDGalNAc(1→)PROA-325 |

**Table S2**. Glycan compositions for chain-B.

CHAIN B

| # | SITE | TYPE | STRUCTURE | SEQUENCE |
|---|---|---|---|---|
| G25 | N17 | FA3 | 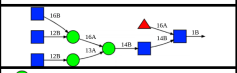 | bDGlcNAc(1→6)[bDGlcNAc(1→2)]aDMan(1→6)<br>[bDGlcNAc(1→2)aDMan(1→3)]bDMan(1→4)bDGlcNAc(1→4)[aLFuc(1→6)]bDGlcNAc(1→)PROB-17 |
| G26 | N61 | M5 | 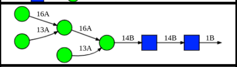 | aDMan(1→6)[aDMan(1→3)]aDMan(1→6)<br>[aDMan(1→3)]bDMan(1→4)bDGlcNAc(1→4)bDGlcNAc(1→)PROB-61 |
| G27 | N74 | FA3G3S2 | 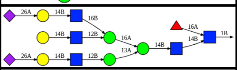 | aDNeu5Ac(2→6)bDGal(1→4)bDGlcNAc(1→6)[bDGal(1→4)bDGlcNAc(1→2)]aDMan(1→6)<br>[aDNeu5Ac(2→6)bDGal(1→4)bDGlcNAc(1→2)aDMan(1→3)]bDMan(1→4)bDGlcNAc(1→4)<br>[aLFuc(1→6)]bDGlcNAc(1→)PROB-74 |
| G28 | N122 | FA2 | 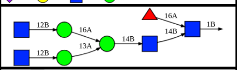 | bDGlcNAc(1→2)aDMan(1→6)[bDGlcNAc(1→2)aDMan(1→3)]bDMan(1→4)bDGlcNAc(1→4)<br>[aLFuc(1→6)]bDGlcNAc(1→)PROB-122 |
| G29 | N149 | FA3 | 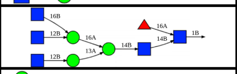 | bDGlcNAc(1→6)[bDGlcNAc(1→2)]aDMan(1→6)<br>[bDGlcNAc(1→2)aDMan(1→3)]bDMan(1→4)bDGlcNAc(1→4)[aLFuc(1→6)]bDGlcNAc(1→)PROB-149 |
| G30 | N165 | M5 | 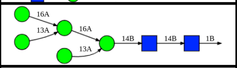 | aDMan(1→6)[aDMan(1→3)]aDMan(1→6)<br>[aDMan(1→3)]bDMan(1→4)bDGlcNAc(1→4)bDGlcNAc(1→)PROB-165 |
| G31 | N234 | M9 | 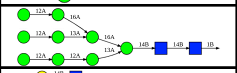 | aDMan(1→2)aDMan(1→6)[aDMan(1→2)aDMan(1→3)]aDMan(1→6)<br>[aDMan(1→2)aDMan(1→2)aDMan(1→3)]bDMan(1→4)bDGlcNAc(1→4)bDGlcNAc(1→)PROB-234 |
| G32 | N282 | FA3G3S1 | 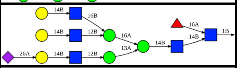 | aDNeu5Ac(2→6)bDGal(1→4)bDGlcNAc(1→2)aDMan(1→3)[bDGal(1→4)bDGlcNAc(1→6)<br>[bDGal(1→4)bDGlcNAc(1→2)]aDMan(1→6)]bDMan(1→4)bDGlcNAc(1→4)<br>[aLFuc(1→6)]bDGlcNAc(1→)PROB-282 |
| G33 | N331 | FA2 | 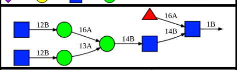 | bDGlcNAc(1→2)aDMan(1→6)[bDGlcNAc(1→2)aDMan(1→3)]bDMan(1→4)bDGlcNAc(1→4)<br>[aLFuc(1→6)]bDGlcNAc(1→)PROB-331 |
| G34 | N343 | FA1 | 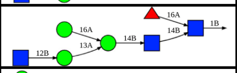 | bDGlcNAc(1→2)aDMan(1→3)[aDMan(1→6)]bDMan(1→4)bDGlcNAc(1→4)<br>[aLFuc(1→6)]bDGlcNAc(1→)PROB-343 |
| G35 | N603 | M5 | 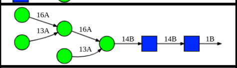 | aDMan(1→6)[aDMan(1→3)]aDMan(1→6)<br>[aDMan(1→3)]bDMan(1→4)bDGlcNAc(1→4)bDGlcNAc(1→)PROB-603 |
| G36 | N616 | FA2 | 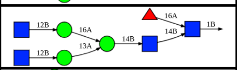 | bDGlcNAc(1→2)aDMan(1→6)[bDGlcNAc(1→2)aDMan(1→3)]bDMan(1→4)bDGlcNAc(1→4)<br>[aLFuc(1→6)]bDGlcNAc(1→)PROB-616 |
| G37 | N657 | Hybrid G1 | 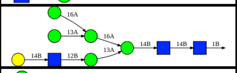 | bDGal(1→4)bDGlcNAc(1→2)aDMan(1→3)[aDMan(1→6)<br>[aDMan(1→3)]aDMan(1→6)]bDMan(1→4)bDGlcNAc(1→4)bDGlcNAc(1→)PROB-657 |
| G38 | N709 | M5 | 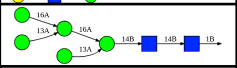 | aDMan(1→6)[aDMan(1→3)]aDMan(1→6)<br>[aDMan(1→3)]bDMan(1→4)bDGlcNAc(1→4)bDGlcNAc(1→)PROB-709 |
| G39 | N717 | M5 | 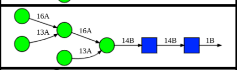 | aDMan(1→6)[aDMan(1→3)]aDMan(1→6)<br>[aDMan(1→3)]bDMan(1→4)bDGlcNAc(1→4)bDGlcNAc(1→)PROB-717 |
| G40 | N801 | M7 | 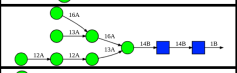 | aDMan(1→2)aDMan(1→2)aDMan(1→3)[aDMan(1→6)<br>[aDMan(1→3)]aDMan(1→6)]bDMan(1→4)bDGlcNAc(1→4)bDGlcNAc(1→)PROB-801 |
| G41 | N1074 | M5 | 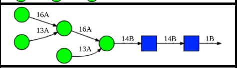 | aDMan(1→6)[aDMan(1→3)]aDMan(1→6)<br>[aDMan(1→3)]bDMan(1→4)bDGlcNAc(1→4)bDGlcNAc(1→)PROB-1074 |
| G42 | N1098 | A2 | 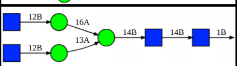 | bDGlcNAc(1→2)aDMan(1→6)<br>[bDGlcNAc(1→2)aDMan(1→3)]bDMan(1→4)bDGlcNAc(1→4)bDGlcNAc(1→)PROB-1098 |
| G43 | N1134 | FA3 | 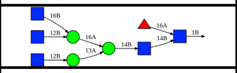 | bDGlcNAc(1→6)[bDGlcNAc(1→2)]aDMan(1→6)<br>[bDGlcNAc(1→2)aDMan(1→3)]bDMan(1→4)bDGlcNAc(1→4)[aLFuc(1→6)]bDGlcNAc(1→)PROB-1134 |
| G44 | N1158 | FA2G2S1 | 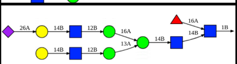 | aDNeu5Ac(2→6)bDGal(1→4)bDGlcNAc(1→2)aDMan(1→6)<br>[bDGal(1→4)bDGlcNAc(1→2)aDMan(1→3)]bDMan(1→4)bDGlcNAc(1→4)<br>[aLFuc(1→6)]bDGlcNAc(1→)PROB-1158 |
| G45 | N1173 | FA4 | 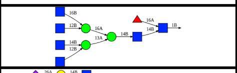 | bDGlcNAc(1→6)[bDGlcNAc(1→2)]aDMan(1→6)[bDGlcNAc(1→4)<br>[bDGlcNAc(1→2)]aDMan(1→3)]bDMan(1→4)bDGlcNAc(1→4)[aLFuc(1→6)]bDGlcNAc(1→)PROB-1173 |
| G46 | N1194 | FA4G4S1 | 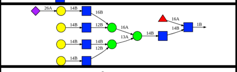 | aDNeu5Ac(2→6)bDGal(1→4)bDGlcNAc(1→6)[bDGal(1→4)bDGlcNAc(1→2)]aDMan(1→6)<br>[bDGal(1→4)bDGlcNAc(1→4)<br>[bDGal(1→4)bDGlcNAc(1→2)]aDMan(1→3)]bDMan(1→4)bDGlcNAc(1→4)<br>[aLFuc(1→6)]bDGlcNAc(1→)PROB-1194 |
| G47 | T323 | O-glycan | 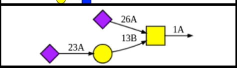 | aDNeu5Ac(2→3)bDGal(1→3)[aDNeu5Ac(2→6)]aDGalNAc(1→)PROB-323 |

**Table S3**. Glycan compositions for chain-C.

CHAIN C

| # | SITE | TYPE | STRUCTURE | SEQUENCE |
|---|---|---|---|---|
| G48 | N17 | FA3 | 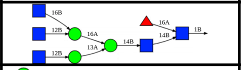 | bDGlcNAc(1→6)[bDGlcNAc(1→2)]aDMan(1→6)<br>[bDGlcNAc(1→2)aDMan(1→3)]bDMan(1→4)bDGlcNAc(1→4)[aLFuc(1→6)]bDGlcNAc(1→)PROC-17 |
| G49 | N61 | M5 | 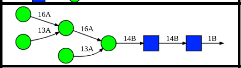 | aDMan(1→6)[aDMan(1→3)]aDMan(1→6)<br>[aDMan(1→3)]bDMan(1→4)bDGlcNAc(1→4)bDGlcNAc(1→)PROC-61 |
| G50 | N74 | A2 | 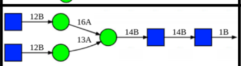 | bDGlcNAc(1→2)aDMan(1→6)<br>[bDGlcNAc(1→2)aDMan(1→3)]bDMan(1→4)bDGlcNAc(1→4)bDGlcNAc(1→)PROC-74 |
| G51 | N122 | M5 | 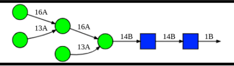 | aDMan(1→6)[aDMan(1→3)]aDMan(1→6)<br>[aDMan(1→3)]bDMan(1→4)bDGlcNAc(1→4)bDGlcNAc(1→)PROC-122 |
| G52 | N149 | FA2 | 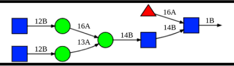 | bDGlcNAc(1→2)aDMan(1→6)[bDGlcNAc(1→2)aDMan(1→3)]bDMan(1→4)bDGlcNAc(1→4)<br>[aLFuc(1→6)]bDGlcNAc(1→)PROC-149 |
| G53 | N165 | FA2G2S1 | 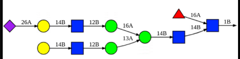 | aDNeu5Ac(2→6)bDGal(1→4)bDGlcNAc(1→2)aDMan(1→6)<br>[bDGal(1→4)bDGlcNAc(1→2)aDMan(1→3)]bDMan(1→4)bDGlcNAc(1→4)<br>[aLFuc(1→6)]bDGlcNAc(1→)PROC-165 |
| G54 | N234 | M9 | 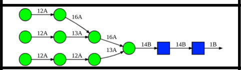 | aDMan(1→2)aDMan(1→6)[aDMan(1→2)aDMan(1→3)]aDMan(1→6)<br>[aDMan(1→2)aDMan(1→2)aDMan(1→3)]bDMan(1→4)bDGlcNAc(1→4)bDGlcNAc(1→)PROC-234 |
| G55 | N282 | A2 | 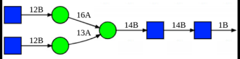 | bDGlcNAc(1→2)aDMan(1→6)<br>[bDGlcNAc(1→2)aDMan(1→3)]bDMan(1→4)bDGlcNAc(1→4)bDGlcNAc(1→)PROC-282 |
| G56 | N331 | FA3G3S1 | 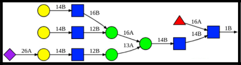 | aDNeu5Ac(2→6)bDGal(1→4)bDGlcNAc(1→2)aDMan(1→3)[bDGal(1→4)bDGlcNAc(1→6)<br>[bDGal(1→4)bDGlcNAc(1→2)]aDMan(1→6)]bDMan(1→4)bDGlcNAc(1→4)<br>[aLFuc(1→6)]bDGlcNAc(1→)PROC-331 |
| G57 | N343 | FA2 | 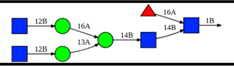 | bDGlcNAc(1→2)aDMan(1→6)[bDGlcNAc(1→2)aDMan(1→3)]bDMan(1→4)bDGlcNAc(1→4)<br>[aLFuc(1→6)]bDGlcNAc(1→)PROC-343 |
| G58 | N603 | M5 | 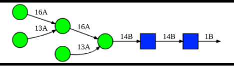 | aDMan(1→6)[aDMan(1→3)]aDMan(1→6)<br>[aDMan(1→3)]bDMan(1→4)bDGlcNAc(1→4)bDGlcNAc(1→)PROC-603 |
| G59 | N616 | FA2 | 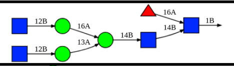 | bDGlcNAc(1→2)aDMan(1→6)[bDGlcNAc(1→2)aDMan(1→3)]bDMan(1→4)bDGlcNAc(1→4)<br>[aLFuc(1→6)]bDGlcNAc(1→)PROC-616 |
| G60 | N657 | Hybrid G1 | 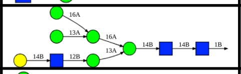 | bDGal(1→4)bDGlcNAc(1→2)aDMan(1→3)[aDMan(1→6)<br>[aDMan(1→3)]aDMan(1→6)]bDMan(1→4)bDGlcNAc(1→4)bDGlcNAc(1→)PROC-657 |
| G61 | N709 | M5 | 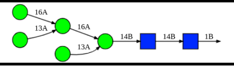 | aDMan(1→6)[aDMan(1→3)]aDMan(1→6)<br>[aDMan(1→3)]bDMan(1→4)bDGlcNAc(1→4)bDGlcNAc(1→)PROC-709 |
| G62 | N717 | M6 | 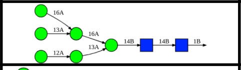 | aDMan(1→6)[aDMan(1→3)]aDMan(1→6)<br>[aDMan(1→2)aDMan(1→3)]bDMan(1→4)bDGlcNAc(1→4)bDGlcNAc(1→)PROC-717 |
| G63 | N801 | M5 | 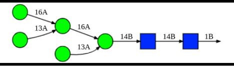 | aDMan(1→6)[aDMan(1→3)]aDMan(1→6)<br>[aDMan(1→3)]bDMan(1→4)bDGlcNAc(1→4)bDGlcNAc(1→)PROC-801 |
| G64 | N1074 | M5 | 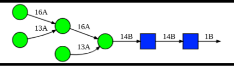 | aDMan(1→6)[aDMan(1→3)]aDMan(1→6)<br>[aDMan(1→3)]bDMan(1→4)bDGlcNAc(1→4)bDGlcNAc(1→)PROC-1074 |
| G65 | N1098 | Hybrid G1S1 | 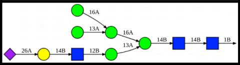 | aDNeu5Ac(2→6)bDGal(1→4)bDGlcNAc(1→2)aDMan(1→3)[aDMan(1→6)<br>[aDMan(1→3)]aDMan(1→6)]bDMan(1→4)bDGlcNAc(1→4)bDGlcNAc(1→)PROC-1098 |
| G66 | N1134 | FA2 | 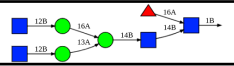 | bDGlcNAc(1→2)aDMan(1→6)[bDGlcNAc(1→2)aDMan(1→3)]bDMan(1→4)bDGlcNAc(1→4)<br>[aLFuc(1→6)]bDGlcNAc(1→)PROC-1134 |
| G67 | N1158 | A2 | 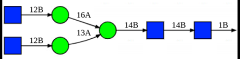 | bDGlcNAc(1→2)aDMan(1→6)<br>[bDGlcNAc(1→2)aDMan(1→3)]bDMan(1→4)bDGlcNAc(1→4)bDGlcNAc(1→)PROC-1158 |
| G68 | N1173 | FA4 | 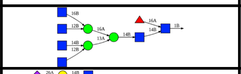 | bDGlcNAc(1→6)[bDGlcNAc(1→2)]aDMan(1→6)[bDGlcNAc(1→4)<br>[bDGlcNAc(1→2)]aDMan(1→3)]bDMan(1→4)bDGlcNAc(1→4)[aLFuc(1→6)]bDGlcNAc(1→)PROC-1173 |
| G69 | N1194 | FA4G4S1 | 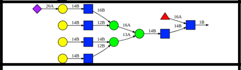 | aDNeu5Ac(2→6)bDGal(1→4)bDGlcNAc(1→6)[bDGal(1→4)bDGlcNAc(1→2)]aDMan(1→6)<br>[bDGal(1→4)bDGlcNAc(1→4)<br>[bDGal(1→4)bDGlcNAc(1→2)]aDMan(1→3)]bDMan(1→4)bDGlcNAc(1→4)<br>[aLFuc(1→6)]bDGlcNAc(1→)PROC-1194 |
| G70 | T323 | O-glycan | 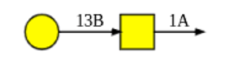 | bDGal(1→3)aDGalNAc(1→)PROC-323 |

**Table S4**. Membrane lipids, their percentages in the membrane patch, and corresponding IUPAC names.

| Lipid | Percentage | IUPAC Name |
|---|---|---|
| POPC | 47% | 1-palmitoyl-2-oleoyl-sn-glycero-3-phosphocholine |
| POPE | 20% | 1-palmitoyl-2-oleoyl-sn-glycero-3-phosphoethanolamine |
| CHL | 15% | (3β)-cholest-5-en-3-ol |
| POPI | 11% | 1-palmitoyl-2-oleoyl-sn-glycero-3-phosphoinositol |
| POPS | 7% | 1-palmitoyl-2-oleoyl-sn-glycero-3-phospho-L-serine |

**Table S5**. Summary of the full-length S protein all-atom MD simulations.

| Replica # | Closed | Open | Mutant |
|---|---|---|---|
| **Rep 1** | 543.60 ns | 1000.50 ns | 1001.30 ns |
| **Rep 2** | 573.80 ns | 1000.30 ns | 1036.20 ns |
| **Rep 3** | 614.10 ns | 1006.30 ns | 1018.40 ns |
| **Rep 4** | - | 404.30 ns | 411.70 ns |
| **Rep 5** | - | 404.30 ns | 407.00 ns |
| **Rep 6** | - | 406.50 ns | 416.60 ns |
| **Total** | **1731.50 ns** | **4222.20 ns** | **4291.20 ns** |

**Table S6**. Accessible Surface Area (ASA) values for protein S' head in Open (**A**) and Closed (**B**). Glycan shielded area is the area covered by glycans. Glycosylated P ASA is the area effectively accessible in the presence of glycans. Non-Glycosylated P ASA is the accessible area in the absence of glycans (i.e. of the nude protein). AVG is average, ST.DEV is standard deviation. A full description is provided in Material and Methods section.

**A** **OPEN**

| Probe Radius [Å] | Glycan shielded area AVG [$Å^2$] | Glycan shielded area ST.DEV. [±$Å^2$] | Non-Glyc. P ASA AVG [$Å^2$] | Non-Glyc. P ASA ST.DEV. [±$Å^2$] | Glyc. P. ASA AVG [$Å^2$] | Glyc. P ASA ST.DEV. [±$Å^2$] |
|---|---|---|---|---|---|---|
| 1.4 | 18779.8 | 1571.4 | 159922.7 | 3174.9 | 141142.9 | 4145.5 |
| 2.0 | 24698.4 | 1665.1 | 130357.0 | 3444.4 | 105658.6 | 4398.0 |
| 3.0 | 31100.5 | 1434.8 | 107239.5 | 3488.8 | 76138.9 | 4272.8 |
| 4.0 | 35301.9 | 1075.7 | 96504.0 | 3490.1 | 61202.1 | 3955.2 |
| 5.0 | 38191.1 | 822.2 | 90792.7 | 3412.8 | 52601.6 | 3498.2 |
| 6.0 | 40356.3 | 986.2 | 87786.2 | 3345.6 | 47429.9 | 3006.9 |
| 7.0 | 42281.5 | 1328.9 | 86439.4 | 3325.4 | 44157.8 | 2699.9 |
| 8.0 | 44131.3 | 1613.1 | 86039.7 | 3301.1 | 41908.4 | 2567.3 |
| 9.0 | 45968.6 | 1808.5 | 86208.7 | 3195.1 | 40240.1 | 2499.6 |
| 10.0 | 47841.9 | 1921.5 | 86837.2 | 3074.8 | 38995.3 | 2502.5 |
| 11.0 | 49744.9 | 2002.0 | 87790.2 | 2951.8 | 38045.4 | 2530.3 |
| 12.0 | 51696.8 | 2098.7 | 89020.7 | 2884.3 | 37324.0 | 2602.2 |
| 13.0 | 53652.4 | 2229.5 | 90407.6 | 2870.1 | 36755.2 | 2710.7 |
| 14.0 | 55595.5 | 2382.7 | 91942.5 | 2864.8 | 36347.0 | 2831.9 |
| 15.0 | 57532.3 | 2558.8 | 93593.0 | 2881.6 | 36060.8 | 2962.7 |

**HEAD**

**B** **CLOSED**

| Probe Radius [Å] | Glycan shielded area AVG [$Å^2$] | Glycan shielded area ST.DEV. [±$Å^2$] | Non-Glyc. P ASA AVG [$Å^2$] | Non-Glyc. P ASA ST.DEV. [±$Å^2$] | Glyc. P. ASA AVG [$Å^2$] | Glyc. P ASA ST.DEV. [±$Å^2$] |
|---|---|---|---|---|---|---|
| 1.4 | 18521.6 | 1571.9 | 157977.0 | 2438.0 | 139455.3 | 3746.0 |
| 2.0 | 24625.2 | 1744.2 | 128117.3 | 2254.9 | 103492.1 | 3617.8 |
| 3.0 | 31256.1 | 1547.3 | 104823.6 | 1963.9 | 73567.5 | 2961.4 |
| 4.0 | 35675.1 | 1210.0 | 93827.7 | 1860.4 | 58152.6 | 2343.5 |
| 5.0 | 38897.3 | 1038.3 | 88215.8 | 1722.4 | 49318.5 | 1767.4 |
| 6.0 | 41537.9 | 1192.1 | 85642.1 | 1590.7 | 44104.2 | 1467.4 |
| 7.0 | 43882.8 | 1438.8 | 84670.6 | 1487.4 | 40787.8 | 1393.2 |
| 8.0 | 45990.7 | 1660.3 | 84545.2 | 1431.6 | 38554.6 | 1501.2 |
| 9.0 | 47982.3 | 1879.2 | 84845.5 | 1393.0 | 36863.2 | 1628.8 |
| 10.0 | 49919.4 | 2037.1 | 85500.4 | 1345.2 | 35581.0 | 1744.0 |
| 11.0 | 51813.2 | 2212.0 | 86438.5 | 1373.7 | 34625.4 | 1861.0 |
| 12.0 | 53749.4 | 2315.6 | 87602.2 | 1393.9 | 33852.7 | 2005.5 |
| 13.0 | 55669.8 | 2427.7 | 88927.4 | 1392.7 | 33257.6 | 2132.1 |
| 14.0 | 57634.5 | 2520.2 | 90436.6 | 1424.9 | 32802.0 | 2235.4 |
| 15.0 | 59581.6 | 2574.8 | 92011.2 | 1422.9 | 32429.5 | 2333.3 |

**Table S7**. Accessible Surface Area values for protein S' stalk in Open (**A**) and Closed (**B**). Same notations as Table S2.

| **A** | | | **OPEN** | | | | | **B** | | | **CLOSED** | | | |
|---|---|---|---|---|---|---|---|---|---|---|---|---|---|---|
| **Probe Radius [Å]** | **Glycan shielded area AVG [$Å^2$]** | **Glycan shielded area ST.DEV. [$±Å^2$]** | **Non-Glyc. P ASA AVG [$Å^2$]** | **Non-Glyc. P ASA ST.DEV. [$±Å^2$]** | **Glyc. P. ASA AVG [$Å^2$]** | **Glyc. P ASA ST.DEV. [$±Å^2$]** | | **Probe Radius [Å]** | **Glycan shielded area AVG [$Å^2$]** | **Glycan shielded area ST.DEV. [$±Å^2$]** | **Non-Glyc. P ASA AVG [$Å^2$]** | **Non-Glyc. P ASA ST.DEV. [$±Å^2$]** | **Glyc. P. ASA AVG [$Å^2$]** | **Glyc. P ASA ST.DEV. [$±Å^2$]** |
| 1.4 | 2514.0 | 294.1 | 14591.3 | 590.3 | 12077.2 | 645.3 | | 1.4 | 2302.4 | 246.1 | 14791.3 | 696.8 | 12488.9 | 666.3 |
| 2.0 | 3292.5 | 348.5 | 13132.1 | 582.5 | 9839.6 | 675.3 | | 2.0 | 3002.9 | 270.0 | 13272.3 | 647.1 | 10269.4 | 618.6 |
| 3.0 | 4244.9 | 359.5 | 12129.2 | 506.5 | 7884.3 | 612.3 | | 3.0 | 3977.4 | 297.4 | 12284.7 | 533.3 | 8307.3 | 528.7 |
| 4.0 | 5139.4 | 359.8 | 11865.1 | 476.1 | 6725.8 | 566.3 | | 4.0 | 4946.9 | 336.3 | 12073.8 | 484.5 | 7126.9 | 495.8 |
| 5.0 | 6030.3 | 366.8 | 11872.6 | 476.2 | 5842.4 | 540.1 | | 5.0 | 5913.4 | 386.2 | 12137.6 | 471.5 | 6224.2 | 481.5 |
| 6.0 | 6903.7 | 382.7 | 11994.4 | 489.5 | 5090.7 | 524.3 | | 6.0 | 6863.0 | 442.9 | 12318.8 | 478.3 | 5455.7 | 468.7 |
| 7.0 | 7741.0 | 412.3 | 12170.6 | 509.2 | 4429.6 | 511.0 | **STALK** | 7.0 | 7777.2 | 503.6 | 12553.7 | 490.7 | 4776.5 | 459.4 |
| 8.0 | 8529.4 | 450.4 | 12374.3 | 531.2 | 3844.9 | 499.5 | | 8.0 | 8630.6 | 558.3 | 12800.5 | 500.1 | 4169.8 | 454.0 |
| 9.0 | 9258.8 | 496.9 | 12591.7 | 558.3 | 3332.9 | 489.5 | | 9.0 | 9419.1 | 612.3 | 13057.7 | 517.9 | 3638.6 | 453.3 |
| 10.0 | 9927.0 | 549.6 | 12810.0 | 584.9 | 2882.9 | 481.9 | | 10.0 | 10138.2 | 661.8 | 13313.4 | 536.9 | 3175.2 | 450.1 |
| 11.0 | 10527.8 | 601.0 | 13026.0 | 611.3 | 2498.2 | 475.9 | | 11.0 | 10787.2 | 705.1 | 13563.3 | 552.6 | 2776.1 | 454.3 |
| 12.0 | 11066.5 | 653.6 | 13233.3 | 641.6 | 2166.8 | 468.9 | | 12.0 | 11362.0 | 744.6 | 13799.7 | 568.7 | 2437.7 | 461.3 |
| 13.0 | 11548.5 | 700.9 | 13432.7 | 674.2 | 1884.3 | 464.0 | | 13.0 | 11872.3 | 786.3 | 14017.8 | 584.6 | 2145.5 | 472.2 |
| 14.0 | 11979.2 | 745.6 | 13621.2 | 702.9 | 1642.0 | 454.4 | | 14.0 | 12332.3 | 828.3 | 14228.7 | 604.4 | 1896.4 | 484.4 |
| 15.0 | 12360.8 | 789.2 | 13794.6 | 735.9 | 1433.8 | 444.1 | | 15.0 | 12740.7 | 857.2 | 14426.7 | 624.0 | 1686.0 | 496.8 |

**Table S8**. Accessible Surface Area values for protein S' $RBD_{RBM}$ in Open (**A**) and Closed (**B**). Same notations as Table S2.

| **A** | | | **OPEN** | | | | | **B** | | | **CLOSED** | | | |
|---|---|---|---|---|---|---|---|---|---|---|---|---|---|---|
| **Probe Radius [Å]** | **Glycan shielded area AVG [$Å^2$]** | **Glycan shielded area ST.DEV. [$±Å^2$]** | **Non-Glyc. P ASA AVG [$Å^2$]** | **Non-Glyc. P ASA ST.DEV. [$±Å^2$]** | **Glyc. P. ASA AVG [$Å^2$]** | **Glyc. P ASA ST.DEV. [$±Å^2$]** | | **Probe Radius [Å]** | **Glycan shielded area AVG [$Å^2$]** | **Glycan shielded area ST.DEV. [$±Å^2$]** | **Non-Glyc. P ASA AVG [$Å^2$]** | **Non-Glyc. P ASA ST.DEV. [$±Å^2$]** | **Glyc. P. ASA AVG [$Å^2$]** | **Glyc. P ASA ST.DEV. [$±Å^2$]** |
| 1.4 | 281.2 | 133.9 | 6625.4 | 439.8 | 6344.2 | 461.7 | | 1.4 | 517.9 | 142.2 | 5855.3 | 301.2 | 5337.4 | 332.1 |
| 2.0 | 368.9 | 162.1 | 5825.7 | 497.9 | 5456.8 | 535.8 | | 2.0 | 716.3 | 176.1 | 4732.7 | 291.5 | 4016.4 | 315.5 |
| 3.0 | 445.9 | 175.3 | 5408.3 | 560.6 | 4962.4 | 577.1 | | 3.0 | 934.3 | 215.0 | 3931.5 | 284.7 | 2997.2 | 294.3 |
| 4.0 | 486.0 | 181.9 | 5341.8 | 621.8 | 4855.9 | 615.7 | | 4.0 | 1053.5 | 225.7 | 3520.0 | 277.8 | 2466.5 | 284.2 |
| 5.0 | 517.7 | 186.3 | 5389.5 | 692.0 | 4871.8 | 667.9 | | 5.0 | 1117.5 | 222.6 | 3297.2 | 280.6 | 2179.7 | 285.2 |
| 6.0 | 538.2 | 187.4 | 5486.8 | 767.1 | 4948.7 | 723.3 | | 6.0 | 1152.0 | 221.2 | 3184.6 | 293.3 | 2032.6 | 295.8 |
| 7.0 | 554.5 | 190.8 | 5615.6 | 837.2 | 5061.1 | 776.8 | **RBD** | 7.0 | 1176.1 | 220.4 | 3129.0 | 312.7 | 1953.0 | 316.9 |
| 8.0 | 572.9 | 194.1 | 5766.7 | 891.4 | 5193.9 | 827.9 | **RBM** | 8.0 | 1200.1 | 218.6 | 3104.0 | 332.4 | 1903.9 | 342.5 |
| 9.0 | 591.6 | 196.2 | 5932.8 | 931.2 | 5341.1 | 876.7 | | 9.0 | 1231.0 | 220.8 | 3103.0 | 352.1 | 1872.0 | 371.2 |
| 10.0 | 608.3 | 197.5 | 6110.5 | 966.6 | 5502.3 | 928.3 | | 10.0 | 1269.6 | 223.6 | 3119.3 | 371.5 | 1849.7 | 400.4 |
| 11.0 | 626.4 | 200.6 | 6301.9 | 1004.4 | 5675.5 | 982.9 | | 11.0 | 1315.8 | 229.8 | 3151.8 | 391.6 | 1836.1 | 429.1 |
| 12.0 | 649.2 | 207.3 | 6509.5 | 1046.2 | 5860.3 | 1041.2 | | 12.0 | 1369.7 | 240.9 | 3198.3 | 412.9 | 1828.6 | 455.9 |
| 13.0 | 678.1 | 217.7 | 6727.7 | 1090.6 | 6049.6 | 1099.9 | | 13.0 | 1426.2 | 254.3 | 3253.1 | 440.8 | 1826.9 | 482.9 |
| 14.0 | 713.2 | 232.1 | 6958.3 | 1137.5 | 6245.1 | 1159.7 | | 14.0 | 1486.2 | 268.6 | 3312.8 | 467.6 | 1826.6 | 509.1 |
| 15.0 | 753.3 | 247.7 | 7199.9 | 1188.8 | 6446.6 | 1222.3 | | 15.0 | 1546.5 | 286.4 | 3377.0 | 499.1 | 1830.4 | 537.4 |

**Table S9**. Accessible Surface Area values for protein S' $RBD_{NON-INTERACTING.}$ in Open (**A**) and Closed (**B**). Same notations as Table S2.

| **A** | | | **OPEN** | | | | | **B** | | | **CLOSED** | | | |
|---|---|---|---|---|---|---|---|---|---|---|---|---|---|---|
| **Probe Radius [Å]** | **Glycan shielded area AVG [$Å^2$]** | **Glycan shielded area ST.DEV. [$±Å^2$]** | **Non-Glyc. P ASA AVG [$Å^2$]** | **Non-Glyc. P ASA ST.DEV. [$±Å^2$]** | **Glyc. P. ASA AVG [$Å^2$]** | **Glyc. P ASA ST.DEV. [$±Å^2$]** | | **Probe Radius [Å]** | **Glycan shielded area AVG [$Å^2$]** | **Glycan shielded area ST.DEV. [$±Å^2$]** | **Non-Glyc. P ASA AVG [$Å^2$]** | **Non-Glyc. P ASA ST.DEV. [$±Å^2$]** | **Glyc. P. ASA AVG [$Å^2$]** | **Glyc. P ASA ST.DEV. [$±Å^2$]** |
| 1.4 | 930.9 | 156.2 | 4536.2 | 257.8 | 3605.3 | 299.0 | | 1.4 | 433.9 | 72.6 | 4000.4 | 195.1 | 3566.5 | 184.4 |
| 2.0 | 1206.9 | 175.4 | 3776.9 | 274.7 | 2570.0 | 302.9 | | 2.0 | 577.8 | 90.2 | 3137.9 | 195.7 | 2560.0 | 183.4 |
| 3.0 | 1428.6 | 167.7 | 3093.0 | 272.7 | 1664.4 | 305.1 | | 3.0 | 704.7 | 104.1 | 2476.8 | 190.2 | 1772.1 | 169.3 |
| 4.0 | 1466.4 | 171.4 | 2641.0 | 264.8 | 1174.5 | 292.1 | | 4.0 | 723.9 | 125.8 | 2130.8 | 191.3 | 1406.9 | 161.4 |
| 5.0 | 1445.1 | 176.7 | 2332.6 | 245.3 | 887.5 | 260.7 | | 5.0 | 708.7 | 135.7 | 1921.7 | 193.6 | 1213.0 | 164.2 |
| 6.0 | 1418.8 | 190.1 | 2118.9 | 241.7 | 700.2 | 230.0 | | 6.0 | 696.1 | 145.3 | 1792.3 | 201.8 | 1096.2 | 172.7 |
| 7.0 | 1399.2 | 206.9 | 1964.4 | 247.3 | 565.2 | 204.9 | **RBD** | 7.0 | 692.8 | 158.8 | 1708.1 | 213.1 | 1015.3 | 184.4 |
| 8.0 | 1383.7 | 220.4 | 1848.1 | 256.5 | 464.4 | 185.8 | **NON-int** | 8.0 | 700.8 | 169.9 | 1651.0 | 220.5 | 950.2 | 194.9 |
| 9.0 | 1375.5 | 231.8 | 1763.1 | 261.5 | 387.6 | 171.8 | | 9.0 | 717.5 | 182.0 | 1609.4 | 230.2 | 891.8 | 204.0 |
| 10.0 | 1373.8 | 243.6 | 1703.9 | 268.2 | 330.1 | 160.7 | | 10.0 | 737.9 | 195.9 | 1579.4 | 241.3 | 841.4 | 211.6 |
| 11.0 | 1380.3 | 250.9 | 1666.6 | 272.6 | 286.3 | 152.1 | | 11.0 | 755.7 | 208.5 | 1553.0 | 251.2 | 797.3 | 218.5 |
| 12.0 | 1392.4 | 259.3 | 1643.7 | 280.0 | 251.2 | 145.9 | | 12.0 | 773.8 | 220.2 | 1532.4 | 261.8 | 758.6 | 225.3 |
| 13.0 | 1407.8 | 267.4 | 1630.4 | 290.0 | 222.6 | 140.9 | | 13.0 | 791.7 | 233.1 | 1513.6 | 270.9 | 721.9 | 232.8 |
| 14.0 | 1421.0 | 275.9 | 1618.6 | 301.5 | 197.5 | 136.6 | | 14.0 | 809.6 | 246.8 | 1497.7 | 279.3 | 688.1 | 240.5 |
| 15.0 | 1432.5 | 286.1 | 1608.7 | 314.9 | 176.2 | 132.5 | | 15.0 | 828.3 | 260.0 | 1485.3 | 290.0 | 657.0 | 247.2 |

**Table S10**. Accessible Surface Area values for protein S' $RBD_{WHOLE}$ in Open (**A**) and Closed (**B**). Same notations as Table S2.

**A** — **OPEN** (WHOLE RBD)

| Probe Radius [Å] | Glycan shielded area AVG [Å$^2$] | Glycan shielded area ST.DEV. [±Å$^2$] | Non-Glyc. P ASA AVG [Å$^2$] | Non-Glyc. P ASA ST.DEV. [±Å$^2$] | Glyc. P. ASA AVG [Å$^2$] | Glyc. P ASA ST.DEV. [±Å$^2$] |
|---|---|---|---|---|---|---|
| 1.4 | 1211.6 | 200.4 | 11159.1 | 477.4 | 9947.5 | 551.2 |
| 2.0 | 1574.9 | 239.8 | 9599.5 | 526.3 | 8024.6 | 636.4 |
| 3.0 | 1874.3 | 246.7 | 8499.3 | 644.6 | 6625.0 | 731.0 |
| 4.0 | 1953.5 | 257.6 | 7980.0 | 708.5 | 6026.5 | 768.9 |
| 5.0 | 1965.1 | 249.8 | 7719.0 | 749.4 | 5753.9 | 805.0 |
| 6.0 | 1959.1 | 244.3 | 7601.5 | 809.0 | 5642.5 | 843.4 |
| 7.0 | 1955.1 | 255.7 | 7573.2 | 877.6 | 5618.1 | 880.6 |
| 8.0 | 1957.4 | 271.2 | 7604.7 | 929.4 | 5647.3 | 916.7 |
| 9.0 | 1964.8 | 282.1 | 7679.9 | 952.9 | 5715.1 | 952.0 |
| 10.0 | 1979.1 | 291.1 | 7797.0 | 972.1 | 5817.9 | 996.1 |
| 11.0 | 2004.9 | 301.5 | 7950.0 | 1003.3 | 5945.1 | 1045.8 |
| 12.0 | 2040.0 | 314.8 | 8131.4 | 1047.0 | 6091.4 | 1100.4 |
| 13.0 | 2084.0 | 329.7 | 8333.2 | 1096.7 | 6249.2 | 1156.6 |
| 14.0 | 2132.6 | 346.2 | 8549.3 | 1149.6 | 6416.7 | 1213.2 |
| 15.0 | 2181.7 | 366.1 | 8775.7 | 1207.9 | 6594.0 | 1272.6 |

**B** — **CLOSED** (WHOLE RBD)

| Probe Radius [Å] | Glycan shielded area AVG [Å$^2$] | Glycan shielded area ST.DEV. [±Å$^2$] | Non-Glyc. P ASA AVG [Å$^2$] | Non-Glyc. P ASA ST.DEV. [±Å$^2$] | Glyc. P. ASA AVG [Å$^2$] | Glyc. P ASA ST.DEV. [±Å$^2$] |
|---|---|---|---|---|---|---|
| 1.4 | 952.4 | 157.6 | 9855.1 | 400.7 | 8902.7 | 425.7 |
| 2.0 | 1295.6 | 199.2 | 7869.4 | 369.2 | 6573.8 | 393.8 |
| 3.0 | 1641.7 | 236.5 | 6407.7 | 327.0 | 4766.0 | 349.0 |
| 4.0 | 1778.3 | 248.0 | 5651.2 | 299.5 | 3872.8 | 311.8 |
| 5.0 | 1826.7 | 246.0 | 5220.2 | 287.9 | 3393.5 | 294.5 |
| 6.0 | 1848.2 | 250.3 | 4978.4 | 296.0 | 3130.1 | 296.5 |
| 7.0 | 1868.6 | 259.4 | 4838.8 | 317.5 | 2970.2 | 313.1 |
| 8.0 | 1901.1 | 262.6 | 4756.5 | 342.3 | 2855.4 | 334.6 |
| 9.0 | 1949.7 | 274.5 | 4714.0 | 373.7 | 2764.2 | 360.6 |
| 10.0 | 2008.5 | 290.5 | 4699.2 | 405.7 | 2690.7 | 389.1 |
| 11.0 | 2072.5 | 304.7 | 4704.3 | 437.7 | 2631.8 | 420.3 |
| 12.0 | 2142.2 | 323.8 | 4727.4 | 467.7 | 2585.2 | 447.9 |
| 13.0 | 2213.6 | 347.2 | 4759.0 | 500.4 | 2545.5 | 479.4 |
| 14.0 | 2290.7 | 373.4 | 4803.3 | 536.8 | 2512.6 | 514.5 |
| 15.0 | 2368.4 | 404.4 | 4854.0 | 576.6 | 2485.5 | 546.4 |

**Table S11**. Summary of principal antibody epitopes reported in the literature. Antibodies denoted by * refer to partial epitopes identified through mutational studies.

| Antibody | Spike Domain | Epitope Residue Numbers | References |
|---|---|---|---|
| COVA1-22 | NTD | 141-156, 246-260 | 57 |
| 4A8 | | 141-156, 246-260 | 55 |
| B38 | RBD | 403-409, 415-421, 455-459, 473-479, 486-505 | 52 |
| 47D11 | | RBD core (338-437, 507-527) | 54 |
| S309 | | 337-344, 356-361, 440-444, glycan at N343 | 53 |
| CR3022 | | 369-392, 427-430, 515-517 | 50,51 |
| VHH-72 | | partial overlap with CR3022 | 60 |
| COV2-2196* | | F486, N487 | 58 |
| COV2-2165* | | F486, N487 | 58 |
| COV2-2130* | | K444, G447 | 58 |
| MAb362* | | Y449, F456, Y489 | 61 |
| 18F3* | | D405, V407 | 59 |
| 7B11* | | L441, S443, L452 | 59 |
| n3021* | | T500, N501, G502 | 63 |
| n3113* | | N354 | 63 |
| n3130* | | D428, F429, E516 | 63 |
| 1A9 | CD | 1111-1130 | 56 |

**Table S12**. Accessible Surface Area values for protein S’ epitopes in Open (**A**) and Closed (**B**). Same notations as Table S2.

| A: OPEN Probe Radius [Å] | Glycan shielded area AVG [$Å^2$] | Glycan shielded area ST.DEV. [±$Å^2$] | Non-Glyc. P ASA AVG [$Å^2$] | Non-Glyc. P ASA ST.DEV. [±$Å^2$] | Glyc. P. ASA AVG [$Å^2$] | Glyc. P ASA ST.DEV. [±$Å^2$] | | B: CLOSED Probe Radius [Å] | Glycan shielded area AVG [$Å^2$] | Glycan shielded area ST.DEV. [±$Å^2$] | Non-Glyc. P ASA AVG [$Å^2$] | Non-Glyc. P ASA ST.DEV. [±$Å^2$] | Glyc. P. ASA AVG [$Å^2$] | Glyc. P ASA ST.DEV. [±$Å^2$] |
|---|---|---|---|---|---|---|---|---|---|---|---|---|---|---|
| 7.2 | 293.8 | 149.9 | 2899.4 | 420.9 | 2605.6 | 452.1 | **B38** | 7.2 | 757.8 | 205.3 | 1599.4 | 297.9 | 841.6 | 256.9 |
| 18.6 | 465.8 | 257.2 | 4406.9 | 1272.7 | 3941.1 | 1232.0 | **B38** | 18.6 | 1098.1 | 462.7 | 1757.4 | 692.6 | 659.3 | 454.0 |
| 7.2 | 368.5 | 94.6 | 795.4 | 193.7 | 426.9 | 193.1 | **S309** | 7.2 | 264.0 | 101.3 | 844.3 | 202.9 | 580.4 | 152.6 |
| 18.6 | 542.8 | 169.7 | 970.3 | 412.0 | 427.5 | 370.2 | **S309** | 18.6 | 287.2 | 190.7 | 643.6 | 405.2 | 356.4 | 305.9 |
| 7.2 | 454.4 | 158.7 | 654.5 | 208.1 | 200.0 | 113.0 | **CR3022** | 7.2 | 23.9 | 24.0 | 25.4 | 24.9 | 1.5 | 4.1 |
| 18.6 | 636.3 | 309.2 | 674.8 | 335.6 | 38.5 | 71.7 | **CR3022** | 18.6 | 0.0 | 0.0 | 0.0 | 0.0 | 0.0 | 0.0 |
| 7.2 | 1279.9 | 526.7 | 3557.6 | 610.8 | 2277.7 | 427.5 | **4A8** | 7.2 | 1277.1 | 600.2 | 3060.0 | 748.5 | 1783.0 | 510.3 |
| 18.6 | 3186.0 | 1264.5 | 6252.3 | 1297.9 | 3066.2 | 835.3 | **4A8** | 18.6 | 2859.2 | 1097.1 | 5020.3 | 1542.7 | 2161.1 | 1108.9 |
| 7.2 | 577.3 | 120.2 | 675.1 | 80.2 | 97.8 | 93.1 | **1A9** | 7.2 | 508.2 | 100.5 | 609.9 | 81.9 | 101.7 | 80.1 |
| 18.6 | 1106.7 | 172.0 | 1113.9 | 175.0 | 7.2 | 24.2 | **1A9** | 18.6 | 866.3 | 159.7 | 872.8 | 159.9 | 6.6 | 26.0 |

**Table S13.** Site specific glycosylation in the two 54-glycans models of SARS-CoV2 S head (~60,000 atoms), i.e. Man9-N234 and N234A. The asterisk on position 234 chain C indicates the critical region where the glycans support the open RBD (chain B in 6VYB) by filling the empty space. We note that the third model is based on these 54-glycans models, but it was not glycosylated at position N234 within chain C.

| chain | resid | glycan | chain | resid | glycan | chain | resid | glycan |
|---|---|---|---|---|---|---|---|---|
| **A** | 61 | Man5 | **B** | 61 | Man5 | **C** | 61 | Man5 |
| **A** | 74 | FA2G | **B** | 74 | FA2G | **C** | 74 | FA2G |
| **A** | 122 | Man5 | **B** | 122 | Man5 | **C** | 122 | Man5 |
| **A** | 149 | FA2G | **B** | 149 | FA2G | **C** | 149 | FA2G |
| **A** | 165 | Man5 | **B** | 165 | Man5 | **C** | 165 | Man5 |
| **A** | 234 | Man9 | **B** | 234 | Man9 | **C** | 234* | Man9/ N234A |
| **A** | 282 | FA2G | **B** | 282 | FA2G | **C** | 282 | FA2G |
| **A** | 331 | FA2G | **B** | 331 | FA2G | **C** | 331 | Man5 |
| **A** | 343 | Man5 | **B** | 343 | Man5 | **C** | 343 | Man5 |
| **A** | 603 | Man5 | **B** | 603 | Man5 | **C** | 603 | Man5 |
| **A** | 616 | A2G | **B** | 616 | A2G | **C** | 616 | A2G |
| **A** | 657 | Man5 | **B** | 657 | Man5 | **C** | 657 | FA2G |
| **A** | 709 | Man5 | **B** | 709 | Man5 | **C** | 709 | Man5 |
| **A** | 717 | Man5 | **B** | 717 | Man5 | **C** | 717 | Man5 |
| **A** | 801 | Man5 | **B** | 801 | Man5 | **C** | 801 | Man5 |
| **A** | 1074 | Man5 | **B** | 1074 | Man5 | **C** | 1074 | Man5 |
| **A** | 1098 | A2G | **B** | 1098 | A2G | **C** | 1098 | A2G |
| **A** | 1134 | FA2G | **B** | 1134 | FA2G | **C** | 1134 | FA2G |

**Table S14.** MD production times (ns) used for the data analysis of the SARS-CoV2 S glycoprotein head. MD1 and MD2 refer to two independent MD trajectories started from different velocities.

| Systems | 46 glycan MD1 | 46 glycan MD2 | 54 glycan |
|---|---|---|---|
| **Man9-N234** | 210 | - | 120 |
| **N234A** | 240 | 240 | 210 |
| **N234-nogly** | - | - | 240 |

**Table S15.** Average backbone RMSD values (Å) calculated for the SARS-CoV2 S protein head models for the RBD domains along the conventional MD trajectories of length indicated in Table 1. Standard deviation values are shown in brackets.

| Systems | RBD (chain A) | RBD (chain B, open) | RBD (chain C) |
|---|---|---|---|
| **Man9-N234 (46-glycans)** | 6.9 (0.7) | 9.9 (1.2) | 8.0 (0.8) |
| **Man9-N234 (54-glycans)** | 9.3 (1.5) | 8.2 (1.3) | 6.7 (1.0) |
| **N234A (45-glycans MD1)** | 7.2 (0.9) | 15.1 (2.9) | 6.5 (1.0) |
| **N234A (45-glycans MD2)** | 7.4 (0.8) | 15.4 (3.0) | 5.8 (0.8) |
| **N234A (53-glycans)** | 6.5 (0.8) | 6.6 (0.9) | 6.2 (1.3) |
| **N234-nogly (53-glycan)** | 6.3 (1.1) | 14.3 (2.9) | 4.0 (0.5) |

### 6. Supplementary Movies

**Movie S1. Glycosylated full-length model of SARS-CoV-2 spike protein.** The movie shows, in the first part, the structure of the glycosylated full-length model of SARS-CoV-2 S protein in the open state (i.e. with 1 RBD in the "up" conformation, namely RBD-A) referred to as "Open" in the main text. The different domains and the color code used for lipids and glycans are indicated in the movie. In the second part, the movie shows the MD dynamics (with CHARMM36 force fields) of the same Open system. Only one MD replica was selected for illustrative purposes.

**Movie S2. N-glycan at N234 progressively inserts itself into the cavity left empty upon the lifting-up of the open RBD.** The movie shows the MD dynamics of the glycosylated head-only model of SARS-CoV-2 S protein described in Section 2 of the Supporting Information. These simulations were conducted with AMBER and GLYCAM force fields. N-glycan at N234 (highlighted with a magenta surface) progressively inserts in the space left empty by the lifting-up of the open RBD. We remark that in this system, based on a different cryo-EM structure (PDB ID: 6VYB), the open RBD is within chain B, here highlighted with an orange transparent surface. N-glycans at N165 and N343 are depicted with steel blue and cornflower blue surfaces, respectively. All the remaining glycans are shown with an admiral blue surface, whereas the protein is represented with gray cartoons.

**Movie S3. N-glycans at N165 and N234 "lock-and-load" the open RBD for infection.** The movie illustrates the structural role of the N-glycans at N165 and N234 in modulating the RBD conformational plasticity. By means of a closed-up view, the movie shows the MD dynamics of the open RBD (i.e., RBD-A) within the glycosylated full-length model of the SARS-CoV-2 S protein (wild-type, referred to as "Open" in the main text). Only one MD replica was selected for illustrative purpose. N-glycans at N165 and N234, and the RBD, are indicated with respective labels in the movie. All the remaining glycans are depicted with a per-residue colored licorice representation (GlcNAc in blue, Fucose in red, Galactose in yellow, Sialic Acid in purple, Mannose in green). Chain A, B and C of the spike trimer are depicted with cyan, red and gray cartoons, respectively.

(13) Shulla, A.; Gallagher, T. Role of Spike Protein Endodomains in Regulating Coronavirus

Entry. *J. Biol. Chem.* **2009**, *284* (47), 32725–32734. https://doi.org/10.1074/jbc.M109.043547.

(23) Krijnse-Locker, J.; Ericsson, M.; Rottier, P. J. M.; Grittiths, G. *Characterization of the Budding Compartment of Mouse Hepatitis Virus: Evidence That Transport from the RER to the Golgi Complex Requires Only One Vesicular Transport Step*.

(24) Klumperman, J.; Krijnse Locker, J.; Meijer, A.; Horzinek, M. C.; Geuze,' And, H. J.; Rottier2, P. J. M. *Coronavirus M Proteins Accumulate in the Golgi Complex beyond the Site of Virion Budding*; 1994.

(25) Wu, E. L.; Cheng, X.; Jo, S.; Rui, H.; Song, K. C.; Dávila-Contreras, E. M.; Qi, Y.; Lee, J.; Monje-Galvan, V.; Venable, R. M.; Klauda, J. B.; Im, W. CHARMM-GUI Membrane Builder toward Realistic Biological Membrane Simulations. *Journal of Computational Chemistry*. 2014. https://doi.org/10.1002/jcc.23702.

(26) Olsson, M. H. M.; SØndergaard, C. R.; Rostkowski, M.; Jensen, J. H. PROPKA3: Consistent Treatment of Internal and Surface Residues in Empirical p K a Predictions. *J. Chem. Theory Comput.* **2011**. https://doi.org/10.1021/ct100578z.

(27) Huang, J.; Mackerell, A. D. CHARMM36 All-Atom Additive Protein Force Field: Validation Based on Comparison to NMR Data. *J. Comput. Chem.* **2013**.

https://doi.org/10.1002/jcc.23354.
(28) Guvench, O.; Hatcher, E.; Venable, R. M.; Pastor, R. W.; MacKerell, A. D. CHARMM Additive All-Atom Force Field for Glycosidic Linkages between Hexopyranoses. *J. Chem. Theory Comput.* **2009**, *5* (9), 2353–2370. https://doi.org/10.1021/ct900242e.
(29) Jang, H.; Muratcioglu, S.; Gursoy, A.; Keskin, O.; Nussinov, R. Membrane-Associated Ras Dimers Are Isoform-Specific: K-Ras Dimers Differ from H-Ras Dimers. *Biochem. J.* **2016**. https://doi.org/10.1042/BCJ20160031.
(30) Beglov, D.; Roux, B. Finite Representation of an Infinite Bulk System: Solvent Boundary Potential for Computer Simulations. *J. Chem. Phys.* **1994**. https://doi.org/10.1063/1.466711.
(31) Jorgensen, W. L.; Chandrasekhar, J.; Madura, J. D.; Impey, R. W.; Klein, M. L. Comparison of Simple Potential Functions for Simulating Liquid Water. *J. Chem. Phys.* **1983**. https://doi.org/10.1063/1.445869.
(32) Phillips, J. C.; Braun, R.; Wang, W.; Gumbart, J.; Tajkhorshid, E.; Villa, E.; Chipot, C.; Skeel, R. D.; Kalé, L.; Schulten, K. Scalable Molecular Dynamics with NAMD. *Journal of Computational Chemistry*. 2005. https://doi.org/10.1002/jcc.20289.
(33) Darden, T.; York, D.; Pedersen, L. Particle Mesh Ewald: An N·log(N) Method for Ewald Sums in Large Systems. *J. Chem. Phys.* **1993**, *98* (12), 10089–10092. https://doi.org/10.1063/1.464397.
(34) Ryckaert, J. P.; Ciccotti, G.; Berendsen, H. J. C. Numerical Integration of the Cartesian Equations of Motion of a System with Constraints: Molecular Dynamics of n-Alkanes. *J. Comput. Phys.* **1977**, *23* (3), 327–341. https://doi.org/10.1016/0021-9991(77)90098-5.
(35) Turq, P.; Lantelme, F.; Friedman, H. L.; Lantelme, F. Brownian Dynamics with Hydrodynamic Interactions. *J. Chem. Phys.* **1977**, *66*, 3045. https://doi.org/10.1063/1.434317.
(36) Martyna, G. J.; Tobias, D. J.; Klein, M. L. Constant Pressure Molecular Dynamics Algorithms. *J. Chem. Phys.* **1994**, *101* (5), 4177–4189. https://doi.org/10.1063/1.467468.
(37) Feller, S. E.; Zhang, Y.; Pastor, R. W.; Brooks, B. R. Constant Pressure Molecular Dynamics Simulation: The Langevin Piston Method. *J. Chem. Phys.* **1995**, *103* (11), 4613–4621. https://doi.org/10.1063/1.470648.
(38) Humphrey, W.; Dalke, A.; Schulten, K. VMD: Visual Molecular Dynamics. *J. Mol. Graph.* **1996**. https://doi.org/10.1016/0263-7855(96)00018-5.
(39) Shrake, A.; Rupley, J. A. Environment and Exposure to Solvent of Protein Atoms. Lysozyme and Insulin. *J. Mol. Biol.* **1973**. https://doi.org/10.1016/0022-2836(73)90011-9.
(40) Pedregosa, F.; Michel, V.; Grisel, O.; Blondel, M.; Prettenhofer, P.; Weiss, R.; Vanderplas, J.; Cournapeau, D.; Pedregosa, F.; Varoquaux, G.; Gramfort, A.; Thirion, B.; Grisel, O.; Dubourg, V.; Passos, A.; Brucher, M.; Perrot, M.; Duchesnay, É. *Scikit-Learn: Machine Learning in Python*; 2011; Vol. 12.
(41) McGibbon, R. T.; Beauchamp, K. A.; Harrigan, M. P.; Klein, C.; Swails, J. M.; Hernández, C. X.; Schwantes, C. R.; Wang, L. P.; Lane, T. J.; Pande, V. S. MDTraj: A Modern Open Library for the Analysis of Molecular Dynamics Trajectories. *Biophys. J.* **2015**, *109* (8), 1528–1532. https://doi.org/10.1016/j.bpj.2015.08.015.
(42) Hsieh, C.-L.; Goldsmith, J. A.; Schaub, J. M.; DiVenere, A. M.; Kuo, H.-C.; Javanmardi, K.; Le, K. C.; Wrapp, D.; Lee, A. G.; Liu, Y.; Chou, C.-W.; Byrne, P. O.; Hjorth, C. K.; Johnson, N. V.; Ludes-Meyers, J.; Nguyen, A. W.; Park, J.; Wang, N.; Amengor, D.; Lavinder, J. J.; Ippolito, G. C.; Maynard, J. A.; Finkelstein, I. J.; McLellan, J. S. Structure-

Based Design of Prefusion-Stabilized SARS-CoV-2 Spikes. *Science (80-. ).* **2020**, eabd0826. https://doi.org/10.1126/science.abd0826.
(43) Waterhouse, A.; Bertoni, M.; Bienert, S.; Studer, G.; Tauriello, G.; Gumienny, R.; Heer, F. T.; De Beer, T. A. P.; Rempfer, C.; Bordoli, L.; Lepore, R.; Schwede, T. SWISS-MODEL: Homology Modelling of Protein Structures and Complexes. *Nucleic Acids Res.* **2018**, *46*. https://doi.org/10.1093/nar/gky427.
(44) Walls, A. C.; Park, Y. J.; Tortorici, M. A.; Wall, A.; McGuire, A. T.; Veesler, D. Structure, Function, and Antigenicity of the SARS-CoV-2 Spike Glycoprotein. *Cell* **2020**, *181* (2), 281-292.e6. https://doi.org/10.1016/j.cell.2020.02.058.
(45) Harbison, A. M.; Brosnan, L. P.; Fenlon, K.; Fadda, E. Sequence-to-Structure Dependence of Isolated IgG Fc Complex Biantennary N-Glycans: A Molecular Dynamics Study. *Glycobiology* **2019**, *29* (1), 94–103. https://doi.org/10.1093/glycob/cwy097.
(46) Maier, J. A.; Martinez, C.; Kasavajhala, K.; Wickstrom, L.; Hauser, K. E.; Simmerling, C. Ff14SB: Improving the Accuracy of Protein Side Chain and Backbone Parameters from Ff99SB. *J. Chem. Theory Comput.* **2015**, *11* (8), 3696–3713. https://doi.org/10.1021/acs.jctc.5b00255.
(47) Kirschner, K. N.; Yongye, A. B.; Tschampel, S. M.; González-Outeiriño, J.; Daniels, C. R.; Foley, B. L.; Woods, R. J. GLYCAM06: A Generalizable Biomolecular Force Field. Carbohydrates. *J. Comput. Chem.* **2008**, *29* (4), 622–655. https://doi.org/10.1002/jcc.20820.
(48) Case, D.; Ben-Shalom, I.; Brozell, S.; … D. C.-… S. T. article is an open. AMBER 2018; University of California: San Francisco, CA, USA, 2018.© 2019 by the Authors.
(49) Watanabe, Y.; Berndsen, Z. T.; Raghwani, J.; Seabright, G. E.; Allen, J. D.; Pybus, O. G.; McLellan, J. S.; Wilson, I. A.; Bowden, T. A.; Ward, A. B.; Crispin, M. Vulnerabilities in Coronavirus Glycan Shields despite Extensive Glycosylation. *Nat. Commun.* **2020**, *11* (1), 2688. https://doi.org/10.1038/s41467-020-16567-0.
(50) Yuan, M.; Wu, N. C.; Zhu, X.; Lee, C.-C. D.; So, R. T. Y.; Lv, H.; Mok, C. K. P.; Wilson, I. A. A Highly Conserved Cryptic Epitope in the Receptor-Binding Domains of SARS-CoV-2 and SARS-CoV. *Science (80-. ).* **2020**. https://doi.org/10.1126/science.abb7269.
(51) Huo, J.; Zhao, Y.; Ren, J.; Zhou, D.; Duyvesteyn, H. M. E.; Ginn, H. M.; Carrique, L.; Malinauskas, T.; Ruza, R. R.; Shah, P. N. M.; Tan, T. K.; Rijal, P.; Coombes, N.; Bewley, K. R.; Tree, J. A.; Radecke, J.; Paterson, N. G.; Supasa, P.; Mongkolsapaya, J.; Screaton, G. R.; Carroll, M.; Townsend, A.; Fry, E. E.; Owens, R. J.; Stuart, D. I. Neutralization of SARS-CoV-2 by Destruction of the Prefusion Spike. *Cell Host Microbe* **2020**. https://doi.org/10.1016/j.chom.2020.06.010.
(52) Wu, Y.; Wang, F.; Shen, C.; Peng, W.; Li, D.; Zhao, C.; Li, Z.; Li, S.; Bi, Y.; Yang, Y.; Gong, Y.; Xiao, H.; Fan, Z.; Tan, S.; Wu, G.; Tan, W.; Lu, X.; Fan, C.; Wang, Q.; Liu, Y.; Zhang, C.; Qi, J.; Gao, G. F.; Gao, F.; Liu, L. A Noncompeting Pair of Human Neutralizing Antibodies Block COVID-19 Virus Binding to Its Receptor ACE2. *Science (80-. ).* **2020**, eabc2241. https://doi.org/10.1126/science.abc2241.
(53) Pinto, D.; Park, Y.; Beltramello, M.; Walls, A. C.; Tortorici, M. A.; Bianchi, S.; Jaconi, S.; Culap, K.; Zatta, F.; Marco, A. De; Peter, A.; Guarino, B.; Spreafico, R.; Cameroni, E.; Case, J. B.; Chen, R. E.; Havenar-daughton, C.; Snell, G.; Telenti, A.; Virgin, H. W.; Lanzavecchia, A.; Diamond, M. S.; Fink, K.; Veesler, D.; Corti, D. Cross-Neutralization of SARS-CoV-2 by a Human Monoclonal SARS-CoV Antibody. *Nature* **2020**, 1–10. https://doi.org/10.1038/s41586-020-2349-y.

(54) Wang, C.; Li, W.; Drabek, D.; Okba, N. M. A.; van Haperen, R.; Osterhaus, A. D. M. E.; van Kuppeveld, F. J. M.; Haagmans, B. L.; Grosveld, F.; Bosch, B. J. A Human Monoclonal Antibody Blocking SARS-CoV-2 Infection. *Nat. Commun.* **2020**, *11* (1), 1–6. https://doi.org/10.1038/s41467-020-16256-y.
(55) Chi, X.; Yan, R.; Zhang, J.; Zhang, G.; Zhang, Y.; Hao, M.; Zhang, Z.; Fan, P.; Dong, Y.; Yang, Y.; Chen, Z.; Guo, Y.; Zhang, J.; Li, Y.; Song, X.; Chen, Y.; Xia, L.; Fu, L.; Hou, L.; Xu, J.; Yu, C.; Li, J.; Zhou, Q.; Chen, W. A Neutralizing Human Antibody Binds to the N-Terminal Domain of the Spike Protein of SARS-CoV-2. *Science (80-. ).* **2020**, eabc6952. https://doi.org/10.1126/science.abc6952.
(56) Zheng, Z.; Monteil, V. M.; Maurer-Stroh, S.; Yew, C. W.; Leong, C.; Mohd-Ismail, N. K.; Cheyyatraivendran Arularasu, S.; Chow, V. T. K.; Lin, R. T. P.; Mirazimi, A.; Hong, W.; Tan, Y.-J. Monoclonal Antibodies for the S2 Subunit of Spike of SARS-CoV-1 Cross-React with the Newly-Emerged SARS-CoV-2. *Eurosurveillance* **2020**, *25* (28), 2000291. https://doi.org/10.2807/1560-7917.ES.2020.25.28.2000291.
(57) Brouwer, P. J. M.; Caniels, T. G.; van der Straten, K.; Snitselaar, J. L.; Aldon, Y.; Bangaru, S.; Torres, J. L.; Okba, N. M. A.; Claireaux, M.; Kerster, G.; Bentlage, A. E. H.; van Haaren, M. M.; Guerra, D.; Burger, J. A.; Schermer, E. E.; Verheul, K. D.; van der Velde, N.; van der Kooi, A.; van Schooten, J.; van Breemen, M. J.; Bijl, T. P. L.; Sliepen, K.; Aartse, A.; Derking, R.; Bontjer, I.; Kootstra, N. A.; Wiersinga, W. J.; Vidarsson, G.; Haagmans, B. L.; Ward, A. B.; de Bree, G. J.; Sanders, R. W.; van Gils, M. J. Potent Neutralizing Antibodies from COVID-19 Patients Define Multiple Targets of Vulnerability. *Science (80-. ).* **2020**, eabc5902. https://doi.org/10.1126/science.abc5902.
(58) Zost, S. J.; Gilchuk, P.; Case, J. B.; Binshtein, E.; Chen, R. E.; Nkolola, J. P.; Schäfer, A.; Reidy, J. X.; Trivette, A.; Nargi, R. S.; Sutton, R. E.; Suryadevara, N.; Martinez, D. R.; Williamson, L. E.; Chen, E. C.; Jones, T.; Day, S.; Myers, L.; Hassan, A. O.; Kafai, N. M.; Winkler, E. S.; Fox, J. M.; Shrihari, S.; Mueller, B. K.; Meiler, J.; Chandrashekar, A.; Mercado, N. B.; Steinhardt, J. J.; Ren, K.; Loo, Y. M.; Kallewaard, N. L.; McCune, B. T.; Keeler, S. P.; Holtzman, M. J.; Barouch, D. H.; Gralinski, L. E.; Baric, R. S.; Thackray, L. B.; Diamond, M. S.; Carnahan, R. H.; Crowe, J. E. Potently Neutralizing and Protective Human Antibodies against SARS-CoV-2. *Nature* **2020**, 1–7. https://doi.org/10.1038/s41586-020-2548-6.
(59) Tai, W.; Zhang, X.; He, Y.; Jiang, S.; Du, L. Identification of SARS-CoV RBD-Targeting Monoclonal Antibodies with Cross-Reactive or Neutralizing Activity against SARS-CoV-2. *Antiviral Res.* **2020**, *179*, 104820. https://doi.org/10.1016/j.antiviral.2020.104820.
(60) Wrapp, D.; De Vlieger, D.; Corbett, K. S.; Schepens, B.; Saelens, X.; Mclellan, J. S. Article Structural Basis for Potent Neutralization of Betacoronaviruses by Single-Domain Camelid Antibodies. *Cell* **2020**, *181*, 1004-1015.e15. https://doi.org/10.1016/j.cell.2020.04.031.
(61) Ejemel, M.; Li, Q.; Hou, S.; Schiller, Z. A.; Wallace, A. L.; Amcheslavsky, A.; Yilmaz, N. K.; Toomey, J. R.; Schneider, R.; Close, B. J.; Chen, D.-Y.; Conway, H. L.; Saeed, M.; Cavacini, L. A.; Klempner, M. S.; Schiffer, C. A.; Wang, Y. IgA MAb Blocks SARS-CoV-2 Spike-ACE2 Interaction Providing Mucosal Immunity. *bioRxiv* **2020**, 2020.05.15.096719. https://doi.org/10.1101/2020.05.15.096719.
(62) Grant, O. C.; Montgomery, D.; Ito, K.; Woods, R. J. Analysis of the SARS-CoV-2 Spike Protein Glycan Shield: Implications for Immune Recognition. https://doi.org/10.1101/2020.04.07.030445.

(63) Wu, Y.; Li, C.; Xia, S.; Tian, X.; Kong, Y.; Wang, Z.; Gu, C.; Zhang, R.; Tu, C.; Xie, Y.; Yang, Z.; Lu, L.; Jiang, S.; Ying, T. Identification of Human Single-Domain Antibodies against SARS-CoV-2. *Cell Host Microbe* **2020**. https://doi.org/10.1016/j.chom.2020.04.023.
